# A distinct CD115^-^ erythro-myeloid precursor present at the maternal-embryonic interface and in the bone marrow of adult mice

**DOI:** 10.1101/2021.07.24.453629

**Authors:** Shweta Tikoo, Rohit Jain, Brendon Martinez, Renhua Song, Matthias Wielscher, Simone Rizzetto, Lisa E Shaw, Andrew J Mitchell, Maria Elizabeth Torres-Pacheco, Fabio Luciani, Matthias Farlik, Justin JL Wong, Steffen Jung, Stuart T Fraser, Wolfgang Weninger

**Author notes:** These authors contributed equally to this work (Shweta Tikoo, Rohit Jain). Co-corresponding authors: Rohit Jain, The Centenary Institute, Locked Bag No. 6, Newtown, NSW 2042, Australia, Wolfgang Weninger, M.D., The Centenary Institute, Locked Bag No. 6, Newtown, NSW 2042, Australia.

## Abstract

During ontogeny, macrophages develop from CD115^+^ precursors, including erythro-myeloid progenitors (EMP). EMP arise in the embryonic yolk sac, the primary site of early haematopoiesis. In adults, CD115^+^ bone marrow-derived monocytes represent essential macrophage precursors. Herein, we identify a CD115^-^ macrophage precursor within the adult bone marrow that is unrelated to the classical monocyte lineage but rather shares transcriptomic and functional characteristics of embryonic EMP. These EMPROR (for <u>E</u>rythro <u>M</u>yeloid <u>Pr</u>ecurs<u>or</u>) cells are capable of efficiently generating macrophages in disease settings. During early development, EMPROR cells were largely absent from the yolk sac but were instead found at the embryonic-maternal interface in the uterine wall. Unexpectedly, the latter site contains robust haematopoietic activity and harbours defined embryonic haematopoietic progenitor cells, including classical CD115^+^ EMP. Our data suggest the existence of an alternative pathway of macrophage generation in the adult. Further, we uncover a hitherto unknown site of earliest blood cell development.

## Introduction

Macrophages represent a heterogeneous family of immune cells involved in the regulation of tissue homeostasis and inflammatory responses following injury, pathogen invasion and neoplastic transformation (*1–4*). It is now well accepted that macrophages develop by two distinct pathways depending on the organ and inflammatory state (*2, 5, 6*). During homeostasis, most tissue-resident macrophages are established prenatally and self-renew throughout adulthood independent of the bone marrow (BM) supply. During embryonic development, macrophages arise from distinct precursors, the earliest of which are the yolk sac-derived macrophage precursors (MP) and erythro-myeloid progenitors (EMP) (*5–10*). While MP and EMP-derived macrophages seed all embryonic organs, the precursors themselves are believed to be consumed during development and have never been identified in adult tissues (*5, 8*). Post-partum, haematopoietic stem cell (HSC)-derived monocytes replenish several macrophage niches resulting in ontogenic heterogenity within macrophage lineages in selected tissues (*7, 11, 12*). Similarly, in the setting of inflammation or within tumours in the adult, circulating Ly6C^hi^ monocytes infiltrate tissues where they differentiate into macrophages (*13–15*). Although monocyte-derived macrophages acquire gene expression profiles and features reminiscent of tissue-resident macrophages (*16, 17*), whether they attain all functional characteristics of tissue-resident macrophages is still unclear. In addition, macrophage populations within a given microenvironment are not uniform; rather, functionally specialised sub-populations are present depending on their anatomical localisation (*12, 18–21*). While during inflammation and cancer development, monocytes have been reported to be the source of macrophages, the existence of other precursors harbouring macrophage potential has remained elusive (*2*).

## Results

### Identification of a novel macrophage precursor in adult bone marrow

We have previously described a distinct population of tissue-resident macrophages localising to the perivascular space of post-capillary venules in the dermis (*22*) (perivascular macrophages, PVM). As part of the perivascular extravasation unit, PVM mediate the recruitment of leukocytes into the skin during inflammation (*23*). The selective identification of PVM is facilitated by the DPE-GFP transgenic mouse strain (*24*), in which the green fluorescent protein (GFP) is driven by regulatory elements of the CD4 locus (*22*). In addition to dermis (*22*), CD11b^+^CD64^+^F4/80^+^GFP^+^ PVM are present in other organs including the mammary fat pad, the meninges and the superficial layer of CNS cortex of DPE-GFP mice (**Suppl. Fig 1A** and **Suppl. Fig 2A**). Similar to the skin, confocal microscopy of inguinal mammary fat pads revealed that GFP^+^ cells lined blood vessels (**Suppl. Fig. 1B, Left panel**). Co-staining with F4/80 antibody confirmed the identity of these cells to be macrophages (**Suppl. Fig. 1B, right panel**). Intravital multi-photon microscopy (*25*) illustrated that these cells were sessile with dendrites that actively probe the surrounding tissue microenvironment (**Suppl. Fig. 1C** and **Suppl. Video 1**). In contrast to PVM, other tissue-resident macrophages such as microglia, Kupffer cells, and lung macrophages did not express GFP in DPE-GFP mice (**Suppl. Fig. 2A-C**).

To determine the turnover kinetics of GFP^+^ PVM in normal tissues, we performed BM chimera studies where irradiated, congenic PTPRC^a^ (CD45.1) mice received BM of DPE-GFP:RAG-1^-/-^ mice (CD45.2) mixed with 1% BM from wildtype C57BL/6 mice (to reconstitute the adaptive immune system). Seven to eight months post-reconstitution, very few, if any, GFP^+^ cells were present in the ear skin, circulation, spleen, or mammary fat pads, consistent with the idea that PVM are long-lived and/or self-renew **(Fig. 1A).** Nevertheless, after 15 months, approximately 5% of PVM in the skin were GFP^+^, revealing slow replenishment from the BM (**Suppl. Fig. 1D**).

**Figure 1:**
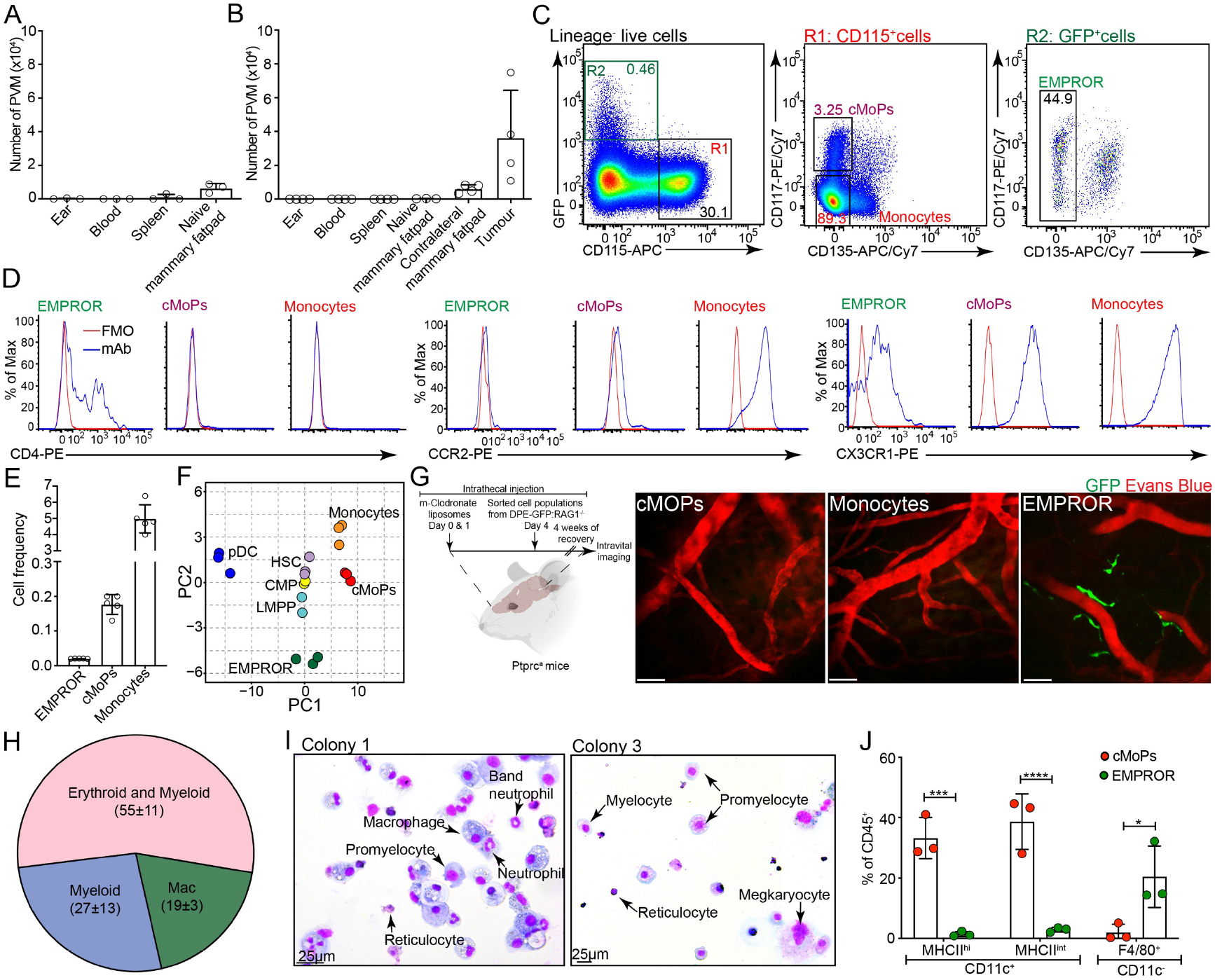
Identification, characterisation and lineage potential of EMPROR cells. **A**, Quantification of donor-derived PVM in various tissues under homeostasis, 7-8 months post bone-marrow reconstitution, n=3 mice, data is representative of two independent experiments. Bars represent mean ± S.D.; **B**, Enumeration of donor-derived PVM in various tissues post-tumour implantation, 10 weeks after bone marrow reconstitution, n=4 mice for all tissues except naïve mammary fat pad, n=3 mice. Bars represent mean ± S.D. Data is representative of one of two independent experiments; **C**, Flow cytometric analysis of bone marrow of DPE-GFP:RAG1^-/-^ mice highlighting the presence of cMoPs, monocytes and EMPROR cells. Lineage markers consist of CD3e, Ly6G, NK1.1, CD19 and Siglec-H. Representative flow plots from one of five independent experiments are shown; **D**, Histogram plots revealing the surface expression of CD4, CCR2 and CX3CR1 on cMoPs, monocytes and EMPROR cells. Representative flow plots from the analysis of one of three independent experiments are shown; **E**, Quantification of the cell frequency of cMoPs, monocytes and EMPROR cells within the adult bone marrow. Data are cumulative of five independent experiments. Bars represent mean ± S.D.; **F**, 2D-PCA plot depicting the transcriptional diversity between HSC (purple), LMPP (light blue), CMP (yellow), cMoPs (red), monocytes (orange), plasmacytoid dendritic cells (pDC, dark blue) and EMPROR cells (green). The data for cMoPs, monocytes, pDC and EMPROR cells was generated from the sequencing of sorted precursors from three independent sort experiments (bone marrow from n=5-7 mice were pooled for individual sorts). Publicly available datasets were used for carrying out comparisons with HSC, LMPP and CMP. ComBat^1, 2^, based on the empirical Bayes method, was applied to the values normalised by variance stabilising transformation from DESeq2 for the batch effect adjustment between samples^1, 2^; **G**, Intravital imaging of the meningeal vasculature in mice reconstituted intrathecally with sorted cMoPs, monocytes and EMPROR cells. Intravital microscopy confirmed the presence of GFP^+^ PVM post-adoptive transfer of EMPROR cells within the CNS. Data representative of three independent experiments wherein the CNS was repleted with EMPROR cells (n=3 mice), cMoPs and monocytes (n=4) sorted from the bone marrow cells of DPE-GFP:RAG1^-/-^ mice; **H**, Pie chart displaying the proportion of macrophage (Mac), myeloid (macrophage and neutrophil) and erythro-myeloid (erythroid with macrophage and neutrophil lineages) colonies generated post-culture of sorted EMPROR cells in MethoCult™ GF M3434 containing transferrin, SCF, IL-3, IL-6, EPO, and supplemented with TPO, GM-CSF and M-CSF. Data are cumulative of three independent sort and cell culture experiments with a total of 34 individual colonies analysed by flow cytometry. The values represent % of colonies ± S.D. for each population; **I**, Giemsa staining of colonies illustrating erythroid as well as myeloid lineage potential generated post-culture of sorted EMPROR cells in MethoCult™ GF M3434 containing transferrin, SCF, IL-3, IL-6, EPO, and supplemented with TPO, GM-CSF and M-CSF. Representative images from two of 20 separate colonies from four independent experiments; **J,** Dendritic cell lineage potential of EMPROR cells and cMoPs. Data are cumulative of three independent sort and culture experiments. Bars represent mean ± S.D. One-way ANOVA was employed; ****P ≤0.0001, ***P ≤0.001, **P ≤0.01, S.D. -Standard deviation.

In pathological settings, such as the tumour milieu, increased numbers of PVM can be found at sites of neoangiogenesis (*26*). In order to investigate the origin of PVM under such conditions, we utilised a murine breast cancer model. Following implantation of syngeneic E0771 medullary breast adenocarcinoma cells (*27, 28*) into the inguinal mammary fat pad of DPE-GFP:RAG-1^-/-^ mice, we observed large numbers of GFP^+^ CD163^+^ macrophages within tumours, particularly in areas of pronounced vascular development at the tumour periphery (**Suppl. Fig. 1E**). Phenotypically, these GFP^+^ macrophages were CD45^+^CD3e^-^B220^-^CD11b^+^CD64^+^F4/80^+^ (**Suppl. Fig. 1F**). Intravital multiphoton microscopy confirmed the presence of GFP^+^ PVM in close proximity to the tumour vasculature (**Suppl. Fig. 1G** and **Suppl. Video 2**).

When E0771 breast cancer cells were implanted into [DPE-GFP:RAG-1^-/-^→PTPRC^a^] BM chimeras (reconstituted for 6-8 weeks), GFP^+^ macrophages developed within the tumours, but not in other organs (**Fig. 1B** and **Suppl. Fig. 3A**). The BM origin of *de novo* developing PVM at inflammatory sites was further studied in a macrophage depletion model wherein [DPE-GFP:RAG-1^-/-^→PTPRC^a^] BM chimeras received mannosylated clodronate liposomes intrathecally (i.t.) in order to deplete meningeal macrophages **(Suppl. Fig. 4A)**. Four weeks post-injection, intravital multiphoton microscopy in the CNS revealed the presence of GFP^+^ PVM lining the post-capillary venules within the meningeal space **(Suppl. Fig. 4A**), indicating that donor BM precursors have the capacity to replenish the PVM pool in the CNS. This was further corroborated by direct i.t. transfer of bulk BM cells from DPE-GFP x RAG-1^-/-^ into PTPRC^a^ mice that were previously depleted of meningeal macrophages **(Suppl. Fig. 4B).** After allowing four weeks of reconstitution, intravital microscopy revealed the presence of GFP^+^ PVM lining the post-capillary venules within the meningeal space **(Suppl. Fig. 4B)**. Together, these data indicate that BM-derived precursors of DPE-GFP mice can generate GFP^+^ PVM in the setting of tumourigenesis or inflammation in adult animals.

As PVM arose from the adult BM, the most likely PVM precursor candidate would be circulating Ly6C^hi^CD11b^+^ monocytes. However, GFP-expressing monocytes were never observed in DPE-GFP:RAG-1^-/-^→PTPRC^a^ BM chimeras (**Suppl. Fig. 3B and C**). While this does not exclude monocytes as PVM precursors, as they may turn on GFP expression during their differentiation into macrophages, it is also plausible that PVM may be derived from an alternative cell source. Indeed, flow cytometric analysis of BM of DPE-GFP:RAG-1^-/-^ mice revealed the presence of GFP^+^ cells (**Suppl. Fig. 3D**). Moreover, when DPE-GFP:RAG-1^-/-^ BM was fractionated into GFP^+^ and GFP^-^ cells and transferred i.t. into meningeal macrophage-depleted mice, only the GFP^+^ fraction gave rise to a network of GFP^+^ macrophages lining blood vessels (**Suppl. Fig. 4C**), morphologically reminiscent of PVM in DPE-GFP mice. GFP^+^ cells readily accumulated i.t. administered TRITC-dextran (**Suppl. Fig. 4C**), indicating that these macrophages are functionally mature. These findings demonstrate that a precursor of GFP^+^ perivascular macrophages resides in the adult BM.

Flow cytometric analysis of the BM of DPE-GFP and DPE-GFP:RAG-1^-/-^ mice showed that lin^-^CD115^+^CD117(cKit)^-^CD135(Flt3)^-^ monocytes and their precursors, including lin^-^CD115^+^CD117^+^CD135^+^ monocyte dendritic cell precursors (MDP) (*29, 30*) and lin^-^ CD115^+^CD117^+^CD135^-^ common monocyte precursors (cMoPs) (*31*), did not express GFP (**Fig. 1C**). Rather, GFP expression within the lineage-negative fraction was restricted to a small but distinct population of CD115^-^CD117^int^CD135^-^ cells (**Fig. 1C** and **Suppl. Fig. 5A and B**). A second population of lin^-^GFP^+^ cells which also expresses CD135^+^ represents lymphocyte precursors (**Fig. 1C, Suppl. Fig. 5A and B** and data not shown). These cells were not considered further in the present work. For simplicity, we refer to lin^-^GFP^+^CD115^-^CD117^int^CD135^-^ cells henceforth as EMPROR (for **<u>E</u>**rythro-**<u>M</u>**yeloid **<u>Pr</u>**ecurs**<u>or</u>**) cells (**Fig. 1C** and **Suppl. Fig. 5A**), based on the evidence that they display phenotypic and functional properties previously ascribed to embryonic EMP cells (detailed below). Lack of Sca-1 and CD150 expression (*32, 33*) distinguishes EMPROR cells from HSC (**Suppl. Fig. 5C and D**). Additional analysis revealed that *bona fide* CMP (*34*) did not express GFP in DPE-GFP mice (**Suppl. Fig. 5E**). A proportion of EMPROR cells were found to be positive for CD4, CD11b and Ly6C (**Fig. 1D** and **Suppl. Fig. 5F**), but negative for CCR2 and low/intermediate for CX3CR1 expression **(Fig. 1D)**. The low CD34 expression further distinguished EMPROR cells from common myeloid progenitors (CMP) (**Suppl. Fig. 5G)**. Moreover, EMPROR cells did not express mature macrophage markers including F4/80, CD64, CD206 and CD209b (**Suppl. Fig. 5G**), and were rare as compared with cMoPs and monocytes (**Fig. 1E**).

### EMPROR cells are distinct from the classical monocyte lineage

We next performed comparative global transcriptomic analyses of EMPROR cells, cMoPs, monocytes and mature plasmacytoid dendritic cells (pDC) isolated from DPE-GFP:RAG1^-/-^ BM using next-generation sequencing. The unique transcriptional program of EMPROR cells was highlighted by 2D-PCA analysis when compared with the known macrophage precursors **(Fig. 1F)**. Further bioinformatic comparison with transcriptomic datasets from previously described progenitor populations including HSC (*35*), LMPP (*35*) and CMP (*36*) underscored the unique transcriptional profile of EMPROR cells **(Fig. 1F** and **Supplementary Figure 6A-C)**. To compare gene profiles of EMPROR cells with known macrophage precursors, namely cMoPs and monocytes, we employed the recently described hexagonal and rose diagrams (*16*). These analyses revealed the distinct expression profiles of select gene sets in EMPROR cells (**Suppl. Fig. 6A**). Further transcriptomic analysis corroborated the low abundance of transcripts for *C-fms, Cx3cr1, Ccr2* and *Klf-4* in EMPROR cells when compared to cMoPs and monocytes (**Suppl. Fig. 5H, 6B**). We also found higher expression of *CD4* in EMPROR cells (**Suppl. Fig. 5H, 6B**), low expression levels of *CD34* (**Suppl. Fig. 5H**), and intermediate expression of PU.1 in EMPROR cells (**Suppl. Fig. 5H**). IRF-8 expression was negligible in all the cell types except pDCs (**Suppl. Fig. 5H**). Quantitative RT-PCR validated the differential expression of *CD4*, *Ccr2* and *Cx3cr1* in cMoPs, monocytes and EMPROR cells (**Suppl. Fig. 5I**). Unsupervised K-means clustering analysis of significantly differentially expressed genes present within EMPROR cells, HSC, LMPP, CMP, cMoPs, monocytes and pDC identified six unique clusters (**Suppl. Fig. 6C** and **Suppl. Data 1**). While clusters II, V and VI were shared between several populations, cluster I was specific for EMPROR cells (**Suppl. Fig. 6C** and **Suppl. Data 1)**. Gene ontogeny analysis of cluster I identified enrichment of several biological pathways, including those involed in haematopoiesis, myeloid cell and erythrocytic development (**Suppl. Fig. 6D** and **Suppl. Data 1**). Together, these data highlight that EMPROR cells are distinct from the known haematopoietic and myeloid precursors based on phenotypic and transcriptional analyses.

### Lineage potential of EMPROR cells

The potential of EMPROR cells to generate macrophages *in vitro* was tested by culturing them in macrophage differentiation media. cMoPs and monocytes were employed as experimental controls. All cultured cell populations generated macrophages (**Suppl. Fig. 7A and B**), with cMoPs and EMPROR cells showing robust expansion of CD45^+^CD11b^+^F4/80^+^ cells (**Suppl. Fig. 7B**). The *in vivo* potential to generate GFP^+^ macrophages was tested in the meningeal macrophage depletion model (**Fig. 1G**). FACS-sorted cMoPs, monocytes and EMPROR cells were administered i.t., 3-4 days post-mannosylated clodronate liposome treatment. Imaging was performed four weeks later (**Fig. 1G**). Only EMPROR cells generated GFP^+^ PVM within the brain (**Fig. 1G**), demonstrating their unique potential to give rise to this specialised macrophage subset.

To test their ability to give rise to T cells, EMPROR cells, cMoPs, monocytes, as well as lin^-^CD117^+^ BM cells (positive control) were cultured on OP9 and OP9-DL1 stromal cells as described previously (*37*). Other than lin^-^CD117^+^ BM cells, all cell populations failed to generate T cells (**Suppl. Fig. 7C** and **D**). To decipher their B cell differentiation potential, we employed the B cell MethoCult^TM^ as well as OP-9 co-culture methodology (*37, 38*) (**Suppl. Fig. 7E** and **F**). None of the tested cell populations, except the positive control, generated B cells (**Suppl. Fig. 7E** and **F**). When cultured in MethoCult^TM^ containing transferrin, SCF, IL-3, IL-6, EPO, TPO, GM-CSF and M-CSF, EMPROR cells gave rise to colonies comprising macrophage, neutrophil, megakaryocytic and erythroid lineages (**Fig. 1H, I** and **Suppl. Fig. 8A**), establishing that EMPROR cells possess an erythro-myeloid differentiation potential similar to that described for embryonic EMP (*39*). In contrast, monocytes failed to generate colonies but differentiated into macrophages (data not shown), while cMoP colonies were restricted to the monocyte-macrophage lineage (**Suppl. Fig. 8B**). Finally, the potential to generate dendritic cells (DCs) *in vitro* was assessed (*40*). Unlike cMoPs, EMPROR cells failed to generate CD45^+^CD11b^+^CD11c^+^MHCII^hi^ DCs in the presence of GM-CSF, but gave rise to CD45^+^CD11b^+^CD11c^-^ F4/80^+^ macrophages (**Fig. 1J** and **Suppl. Fig. 8C**). Monocytes did not undergo extensive expansion but differentiated into DCs and macrophages **(Suppl. Fig. 8C** and **D**). Thus, compared to cMoPs and monocytes, EMPROR cells show distinct lineage potential *in vitro*, with erythroid, granulocyte and macrophage progenitor activity whilst lacking DC or lymphoid potential.

Next, we assessed the lineage potential of EMPROR cells *in vivo* during tumour development. To faithfully track cells of donor origin arising from adoptively transferred precursors including erythrocytes, we utilised DPE-GFP:mT/mG mice, in which all cells express a membrane-bound fluorescent reporter (Tomato; mTom). We engrafted monocytes (0.5×10^6^), cMoPs and EMPROR cells (0.5-2×10^5^ each) sorted from the BM of adult DPE-GFP:mT/mG mice into sublethally irradiated congenic PTPRC^a^ mice. E0771 breast tumours were established 24h post-reconstitution and various tissues were harvested three weeks later (**Fig. 2A)**. Transfer of cMoPs and monocytes resulted in scarce mTom^+^ tumour-associated macrophages (TAMs) (**Fig. 2A** and **Suppl. Fig 9A**). In contrast, EMPROR cells gave rise to a robust, GFP^-^ TAM population (**Fig. 2A-C**), and a smaller population of GFP^+^ macrophages (**Fig. 2B** and **Suppl. Fig. 9A-B)**. In addition, EMPROR cells also yielded neutrophils within tumours (**Fig. 2A** and **C**), thereby corroborating their myeloid differentiation potential ascertained *in vitro*.

**Figure 2:**
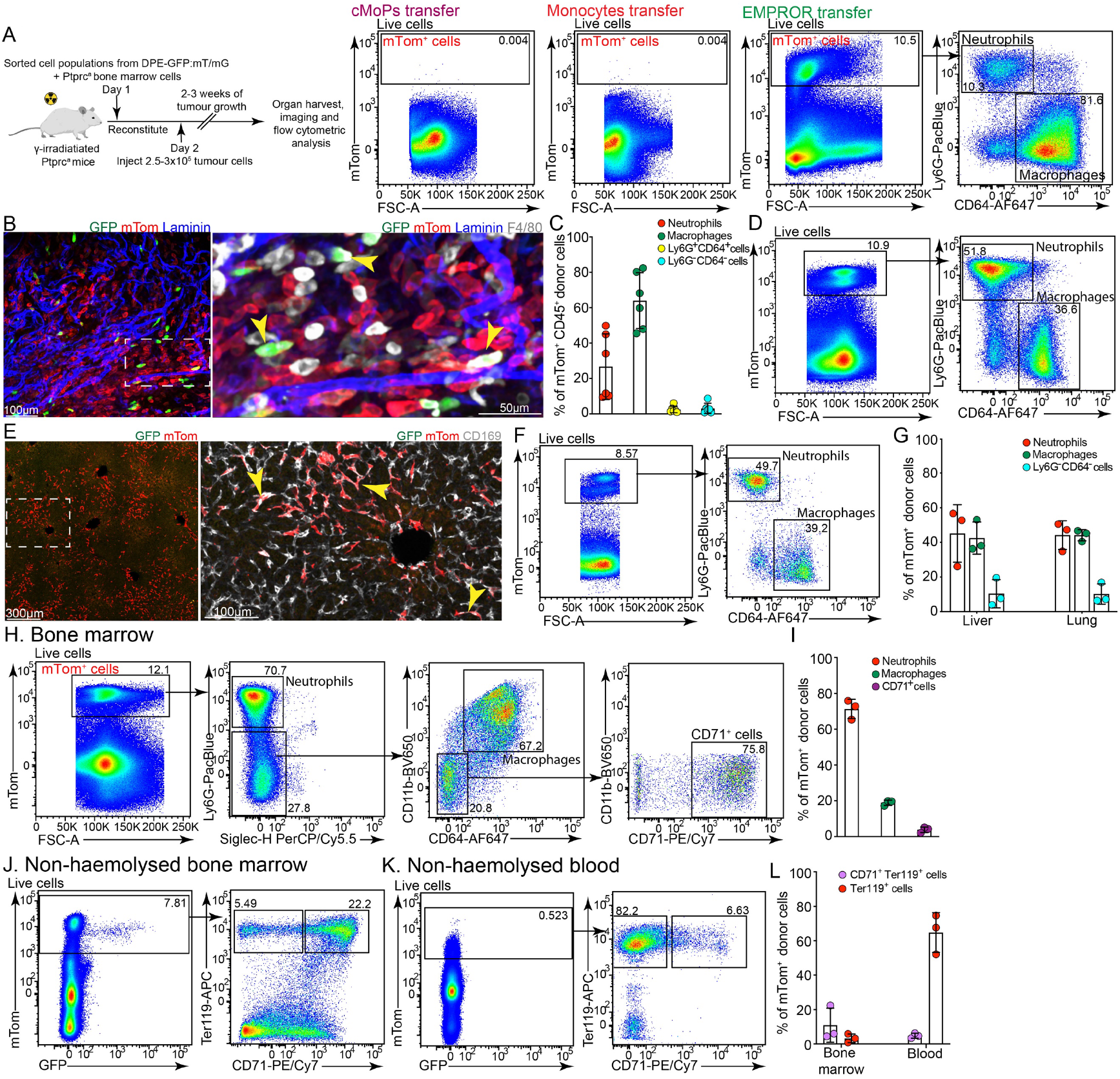
EMPROR cells give rise to erythroid and myeloid cells during tumour development *in vivo*. **A**, Flow cytometric analysis of E0771 tumours harvested from chimeric mice reconstituted with sorted cMoPs, monocytes or EMPROR cells from DPE-GFP:mT/mG mice. cMoPs and monocyte groups show minimal infiltration and expansion of mTom^+^ donor cells. A robust population consisting predominantly of neutrophils and macrophages is observed in tumours from mice reconstituted with EMPROR cells. Representative flow plots from one of six independent sort, adoptive transfer and tumour implantation experiments are shown; **B**, Confocal imaging of immunostained E0771 tumour section harvested from mice reconstituted with EMPROR cells confirms the presence of numerous mTom^+^, mTom^+^F4/80^+^ and mTom^+^F4/80^+^GFP^+^ donor-derived cells (GFP^+^ cells marked by yellow arrowheads), **C**, Quantification of donor-derived immune cell populations detected within the tumour milieu. Data are cumulative of six independent sort, adoptive transfer and tumour implantation experiments. Bars represent mean ± S.D.; **D**, Flow cytometric analysis of liver infiltrating donor cells in chimeric mice reconstituted with EMPROR cells show a robust population of neutrophils and macrophages in tumour-bearing mice; **E**, Confocal microscopy of liver sections obtained from mice reconstituted with EMPROR highlights the presence of numerous mTom^+^ CD169^+^GFP^-^ donor-derived Kupffer cells (marked by yellow arrowheads); **F**, Flow cytometric analysis of lungs from chimeric mice reconstituted with EMPROR cells delineates the presence of mTom^+^ neutrophils and macrophages in tumour-bearing mice; **G**, Quantification of donor-derived immune cell populations detected within the liver and lungs of tumour-bearing mice; **H**, Flow cytometric analysis of the chimeric bone marrow of mice reconstituted with EMPROR cells. Similar to *in vitro* studies, a significant fraction of neutrophils, macrophages and CD71^+^ cells can be observed in tumour-bearing mice; **I**, Quantification of various myeloid and erythroid cell populations in the haemolysed bone marrow of tumour-bearing chimeric mice reconstituted with EMPROR cells; **J**, Identification of erythrocytes and erythrocytic precursors within the non-haemolysed bone marrow of chimeric mice reconstituted with EMPROR cells; **K**, Identification of erythrocytes and erythrocytic precursors within the non-haemolysed blood of chimeric mice reconstituted with EMPROR cells; **L**, Quantification of erythrocytes and erythrocytic precursors within the non-haemolysed bone marrow and blood of tumour bearing chimeric mice reconstituted with EMPROR cells. Representative flow plots from one of three independent sort, adoptive transfer and tumour implantation experiments are depicted for panels D, F, H, J and K. Data presented for panel G, I and L are cumulative of three independent sort, adoptive transfer and tumour implantation experiments. Bars represent mean ± S.D. Images for panel B and E are representative of tissues harvested from tumour bearing chimeric mice. The data presented is representative of one of three independent sort, adoptive transfer and tumour implantation experiments. S.D.-Standard deviation.

Further analysis of other organs from tumour-bearing mice receiving reporter-tagged EMPROR cells revealed that the livers and lungs harboured a substantial number of mTom^+^ neutrophils and macrophages (**Fig. 2D-G** and **Suppl. Fig. 9C**). Confocal microscopic analysis of liver sections identified mTom^+^ macrophages as CD169^+^ Kupffer cells (**Fig. 2E** and **Suppl. Fig. 9C**). Also, the brain contained a small number of donor-derived macrophages (**Suppl. Fig. 9D**), while minimal mTom^+^ cells were found in the thymus reiterating the lack of T cell potential (**Suppl. Fig. 9E**). EMPROR cells also gave rise to haematopoietic cells in the BM including neutrophils, macrophages and CD71^+^Ter119^+^ erythroblasts (**Fig. 2H-J**). Circulating erythrocytes were also derived from EMPROR cells demonstrating the erythroid potential within the EMPROR population (**Fig. 2K** and **L**). Together, these experiments substantiated an erythro-myeloid differentiation potential of EMPROR cells *in vivo* in the context of a tumour-bearing host.

### Identification of MIXER tissue as the initial site for the development of EMPROR cells

EMP are known to arise within the yolk sac during early embryonic development (*39, 41, 42*) and are believed to be consumed pre-partum (*6, 39*). To assess whether EMPROR cells are also present during ontogeny, we employed a differential fluorescent tagged timed-mating strategy, wherein female DPE-GFP mice were mated with male DPE-GFP:mT/mG mice (**Fig. 3A**), allowing for the discrimination of maternal (mTom^-^) and embryonic (mTom^+^) cells. We initially utilised this strategy of differential fluorescent labelling to discern the embryonic tissue from maternal cells within the gravid uterus (**Fig. 3B** and **Suppl. Fig. 10A)**. The first observation of these experiments was the unexpected size of the extraembryonic tissue. Previous anatomical studies using histology have underappreciated the extent to which the embryonic cells invade the uterus as it was not possible to distinguish these tissues based on morphology alone. For simplicity, we refer to this tissue as <u>M</u>aternally <u>I</u>nterfaced e<u>X</u>tra-<u>E</u>mb<u>R</u>yonic (MIXER) tissue. MIXER tissue could be clearly separated from the mTom^+^ embryo proper and yolk sac **(Fig. 3B).** Secondly, we found that MIXER, but not the yolk sac, contained GFP^+^ cells which also expressed mTom **(Fig. 3B** and **Suppl. Fig. 10B).** To corroborate these data, we micro-dissected MIXER from embryo proper and yolk sac. Flow cytometry revealed the presence of mTom^+^GFP^+^CD115^-^ cells in MIXER but not within the yolk sac (**Fig. 3C-E**). These cells displayed a similar phenotype as adult EMPROR cells **(Fig. 3D).** We called these cells eEMPROR (embryonic EMPROR) in concordance with their embryonic origin **(Fig. 3B-D)**. Other than eEMPROR cells, MIXER tissue also harboured CD115+ cells (**Fig. 3D** and **F**), including classical CSF1-R^+^CD117^+^CD45^lo^CD93^+^ EMP (*6, 43*) (hereafter, referred to as CSF1-R^+^ EMP) **(Fig. 3G)**. Unlike eEMPROR cells, CSF1-R^+^ EMP did not show expression of CD4 within the MIXER tissue **(Fig. 3H)** or the developing yolk sac (**Suppl. Fig. 10C** and **D**). Other than CD115^+^ cells, we also found embryonically-derived CD135^+^ and CD117^+^ cells within the MIXER tissue similar to those observed within the yolk sac **(Fig. 3I-J)**.

**Figure 3:**
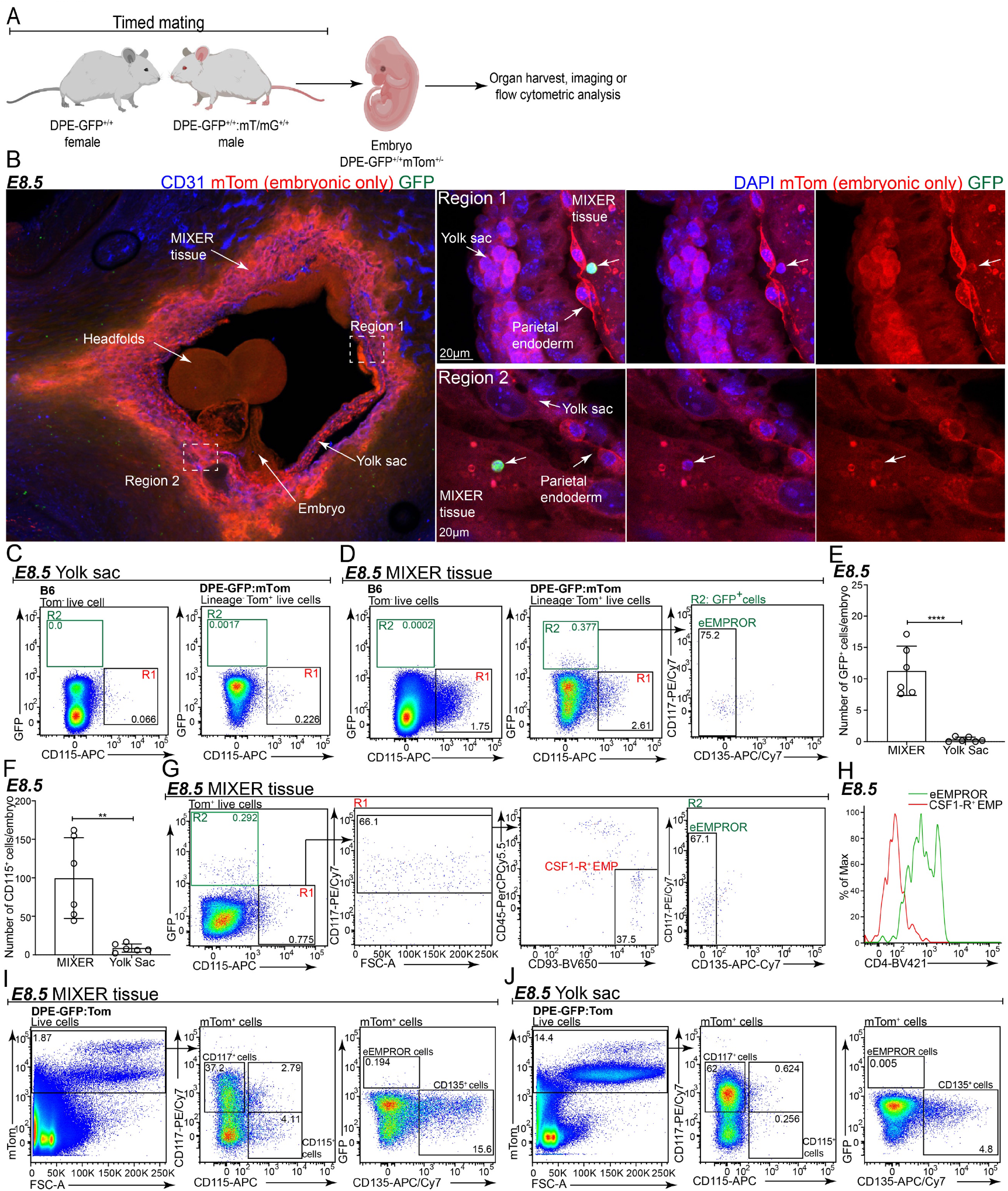
Embryonic EMPROR cells arise within MIXER tissue. **A**, Schematic representation of the timed mating strategy; **B**, Representative confocal image of a section of the E8.5 conceptus revealing the extent of extra-embryonic tissue (MIXER) within the gravid uterus. Higher magnification images highlight the presence of GFP^+^ cells within the MIXER tissue. The images are representative of n=4 embryos from 4 independent timed-mating pairs from three independent experiments; **C** and **D**, Flow cytometric analysis of yolk sac and MIXER tissue from E8.5 embryos identifying the absence or presence of GFP^+^ cells; respectively. Lineage markers consist of CD3e, Ly6G, NK1.1, CD19, Siglec-H, and CD64. Representative flow plots from n=5 independent litters (6-11 embryos/litter) from five separate timed mating pairs from three independent experiments are shown. The flow cytometry data presented is from 8 embryos staged as E8.5 with 6-9 somite pairs; **E,** Bar graph represents the number of GFP^+^ cells observed within the MIXER tissue and the yolk sac; **F,** Bar graph represents the number of CD115^+^ cells observed within the MIXER tissue and the yolk sac. The data presented in panel E and F is cumulative of three independent experiments, n=6 independent litters (46 embryos in total) and is normalised to the number of E8.5 embryos analysed per experiment. Bar represents mean ± S.D, with the symbols representing individual data points from each litter. All embryos were staged between 6-9 somite pairs (E8.5) except one litter staged as 2-6 somite pairs (E8.25-8.5); **G**, Flow cytometric analysis exhibiting the presence of CSF1-R^+^ EMP and eEMPROR cells within the E8.5 MIXER tissue; **H**, Histogram plots depicting CD4 expression on CSF1-R^+^ EMP and eEMPROR cells present within the E8.5 MIXER tissue. Panel G and H, data representative of n=4 independent litters (6-11 embryos/litter) from four separate timed-mating pairs from two independent experiments. The embryos were staged as E8.5 with 6-8 somite pairs; **I**, Flow cytometric analysis of various lineage markers expressed by mTom^+^ cells present within the MIXER tissue. Representative flow plots from n=6 independent litters (6-11 embryos/litter) from six separate timed-mating pairs from three independent experiments are shown. The embryos were staged as E8.5 with 6-9 somite pairs; **J**, Flow cytometric analysis of various lineage markers expressed by mTom^+^ cells present within the yolk sac. Representative flow plots from n=6 independent litters (6-11 embryos/litter) from six separate timed-mating pairs from three independent experiments are shown. The embryos were staged as E8.5 with 6-9 somite pairs. Unpaired t-test was employed; ****P ≤0.0001, **P ≤0.01, S.D. -Standard deviation.

The discovery of MIXER tissue as a site for CSF1-R^+^ EMP and eEMPROR cell residence raised the question as to whether, in general, this tissue displays haematopoietic activity. To test this, we sorted mTom^+^ cells from the MIXER and yolk sac and cultured them on OP-9 stromal cells in media supplemented with SCF, IL-3, IL-6, EPO, TPO, GM-CSF and M-CSF. Flow cytometric analyses highlighted the development of erythroid and myeloid lineages from both tissues **(Fig. 4A-D)**. Furthermore, we observed that similar to yolk sac, most of the leukocytes generated from the MIXER tissue were CD31^+^ (**Fig. 4E-F**). Together, these data suggest that eEMPROR cells develop within an alternative haematopoietic site, namely MIXER, while conventional CSF1-R^+^ EMP are present within the MIXER tissue as well as the yolk sac.

**Figure 4:**
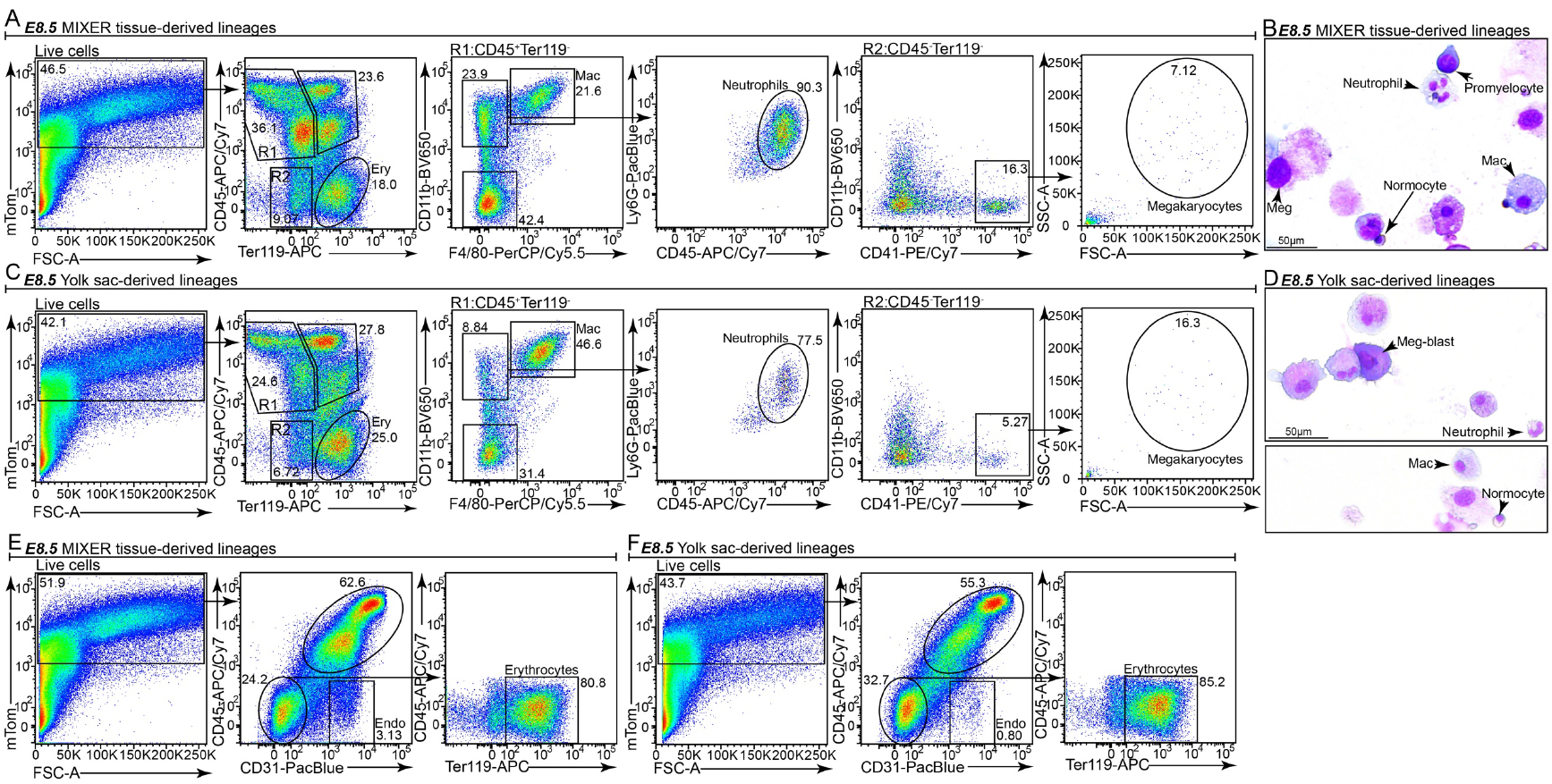
MIXER-derived cells exhibit haematopoietic potential. **A**, Flow cytometric analysis of lineages generated post-culture of mTom^+^ cells sorted from the MIXER tissue. The sorted cells were cultured on OP-9 stromal cells in the presence of SCF, IL-3, IL-6, EPO, TPO, GM-CSF and M-CSF. Representative flow plots from one of three independent sort and culture experiments are shown; **B**, Giemsa staining of colonies depicting erythroid as well as myeloid lineage potential generated post-culture of mTom^+^ cells sorted from the MIXER tissue and cultured on OP-9 stromal cells in the presence of SCF, IL-3, IL-6, EPO, TPO, GM-CSF and M-CSF. Representative image from one of three independent sort and culture experiments is shown; **C**, Flow cytometric analysis of lineages generated post-culture of mTom^+^ cells sorted from yolk sac. The sorted cells were cultured on OP-9 stromal cells in the presence of SCF, IL-3, IL-6, EPO, TPO, GM-CSF and M-CSF. Representative flow plots from one of three independent sort and culture experiments are shown; **D**, Giemsa staining of colonies demonstrating erythroid as well as myeloid lineage potential generated post-culture of mTom^+^ cells sorted from yolk sac and cultured on OP-9 stromal cells in the presence of SCF, IL-3, IL-6, EPO, TPO, GM-CSF and M-CSF. Representative images from one of three independent sort and culture experiments are shown; **E**, Flow cytometric analysis of cell lineages arising from sorted mTom^+^ MIXER derived cells Representative flow plots from one of three independent sort and culture experiments are shown; **F**, Flow cytometric analysis of cell lineages arising from sorted yolk sac-derived cells. Representative flow plots from one of three independent sort and culture experiments are shown. The co-expression of CD45 and CD31 in panel A and B highlights the potential recent emergence of these cells from hemogenic endothelium. Endo refers to CD45^-^CD31^+^ endothelial cells.

### EMPROR cells in later embryonic development

At the later stages of embryonic development, the dorsal aorta showed the highest number of eEMPROR cells (**Fig. 5A** and **5B**). As previously described (*7*), we found a robust population of lin^-^GFP^-^CD115^+^ cells within the foetal liver (**Fig. 5A** and **5B**), which included the embryonic cMoPs and monocytes (*7*) (**Suppl. Fig. 11A**). At E14.5, eEMPROR cells (**Suppl. Fig. 11B**) as well as *bona fide* CD45^+^CD64^+^F4/80^+^GFP^+^ macrophages were detected within the head of embryos (**Fig. 5C**). Confocal microscopy confirmed that many GFP^+^F4/80^+^ macrophages localised to the perivascular space (**Fig. 5D**). Similarly, CD45^+^CD11b^+^CD64^+^F4/80^+^GFP^+^ macrophages were also detected within the skin of E14.5 embryos (**Suppl. Fig. 11C**). This indicated that eEMPROR cells precede mature macrophages during development and may give rise to GFP^+^ PVM in various organs.

**Figure 5:**
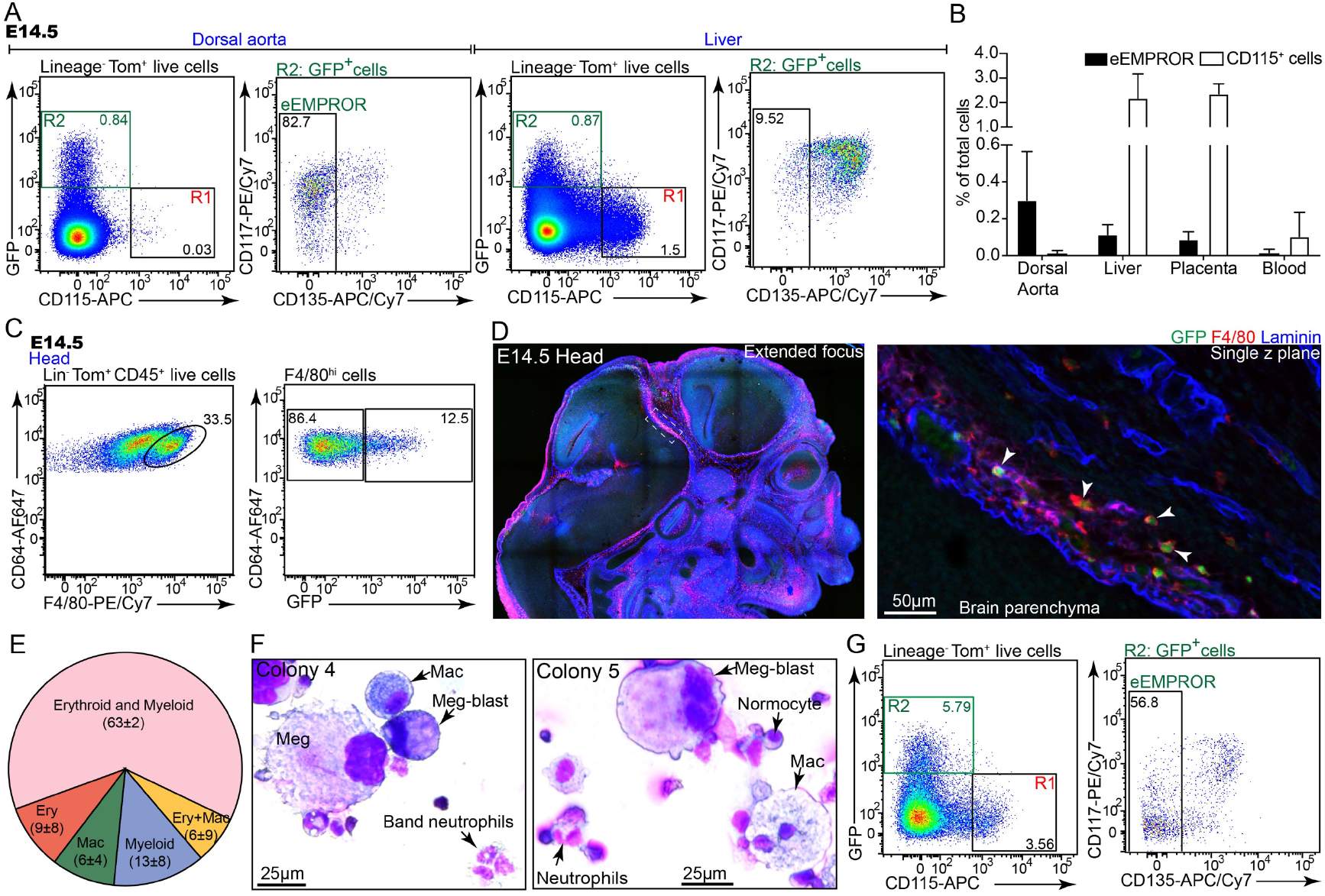
Prenatal development of EMPROR cells. **A,** Flow cytometric analysis of dorsal aorta and liver from E14.5 embryos. The data reveals the presence of eEMPROR cells and lin^-^GFP^-^CD115^+^ precursors within these tissues. Lineage markers consist of CD3e, Ly6G, NK1.1, CD19, Siglec-H, and CD64. Representative flow plots from one of four independent experiments are shown; **B**, Quantification of eEMPROR and CD115^+^ precursors within various haematopoietic organs of E14.5 embryos. Data cumulative of four independent experiments. Bars represent mean ± S.D.; **C**, Flow cytometric analysis of foetal head identifying the presence of GFP^+^ macrophages. Representative flow plots from one of three independent experiments are shown; **D**, Confocal imaging of immunostained E14.5 embryo head shows GFP^+^F4/80^+^ macrophages (white arrowheads) around the brain parenchyma. Representative images of n=3, E14.5 embryos harvested from three independent timed mating pairs from three independent experiments; **E**, Pie chart depicting the proportion of macrophage (Mac), myeloid (macrophage and neutrophil), erythroid and macrophage (Ery+Mac), erythroid (Ery) and erythro-myeloid (erythroid+myeloid, erythroid with macrophage and neutrophil lineages) colonies generated post-culture of sorted eEMPROR cells from foetal liver at E14.5 in MethoCult™ GF M3434 containing transferrin, SCF, IL-3, IL-6, EPO, and supplemented with TPO, GM-CSF and M-CSF. Data is cumulative of 62 colonies from three independent sort and culture experiments. The values represent % of colonies ± S.D. for each population; **F**, Giemsa staining of colonies depicting erythroid as well as myeloid lineage potential generated post-culture of sorted eEMPROR cells in MethoCult™ GF M3434 containing transferrin, SCF, IL-3, IL-6, EPO, and supplemented with TPO, GM-CSF and M-CSF. Data represents images from three of 21 colonies from three independent sort and culture experiments; **G**, Flow cytometric analysis of developing bone marrow from E19.5 embryos. Lineage markers consist of CD3e, Ly6G, NK1.1, CD19, Siglec-H, and CD64. Representative flow plots from one of three independent experiments are shown. Flow plots presented in panel A, C and G are from embryos harvested from independent litters from separate timed-mating pairs from independent experiments. S.D.-Standard deviation.

eEMPROR cells, as well as CD115^+^ cMoPs and monocytes were found within the circulation at E14.5 (**Suppl. Fig. 11D**). Notably, however, the placental labyrinth, which is one of the major sites of haematopoiesis at this stage (*44*), harboured very few GFP^+^ cells (**Suppl. Fig. 11E**). Vitelline arteries and veins were devoid of eEMPROR cells (**Suppl. Fig. 11F**). Similarly, E14.5 yolk sacs were also devoid of eEMPROR cells (**Suppl. Fig. 11G**) but harboured GFP^+^ macrophages (**Suppl. Fig. 11H**).

To formally test their capability of generating macrophages, we sorted cMoPs and eEMPROR cells from E14.5 embryonic livers and cultured them in the presence of M-CSF. Similar to their adult counterparts, both cell populations produced macrophages (**Suppl. Fig. 11 I** and **J**). In colony forming assays, eEMPROR cells gave rise to colonies consisting of macrophage, neutrophil, megakaryocytic as well as erythroid lineages (**Fig. 5E, F** and **Suppl. Fig. 12A-B**), highlighting their erythro-myeloid potential *in vitro*. As expected, monocytes did not generate colonies and cMoP colonies were restricted to the monocyte-macrophage lineage (**Suppl. Fig. 12C – G**).

Finally, flow cytometric analysis of the embryonic BM at E19.5 revealed the emergence of eEMPROR cells pre-partum within the embryonic bone (**Fig. 5G)**. At this stage, embryonic liver also demonstrated a robust population of eEMPROR cells along with embryonic cMoPs and monocytes (**Suppl. Fig. 11K).** Taken together, these experiments highlight the presence of EMPROR cells within the developing haematopoietic organs with their subsequent residence in the prenatal BM.

### Transcriptional profiling establishes the close relationship between embryonic and adult EMPROR cells

Using transcriptional profiling, we further tested the relationship between embryonic and adult EMPROR cells in comparison to cells of the monocytic lineage. To this end, eEMPROR cells and cMoPs were isolated from foetal livers at E14.5 and compared to monocytes, cMoPs and EMPROR cells harvested from adult BM. Uniform manifold approximation and projection (UMAP) analysis combining the first ten principal components revealed that EMPROR cells clustered separately from conventional monocyte lineage cells (**Fig. 6A**). This was further corroborated by hierarchical clustering analysis, which revealed that EMPROR cells and monocytic cells formed separate branches in the dendrogram (**Fig. 6B** and **Suppl. Fig 13A**).

**Figure 6:**
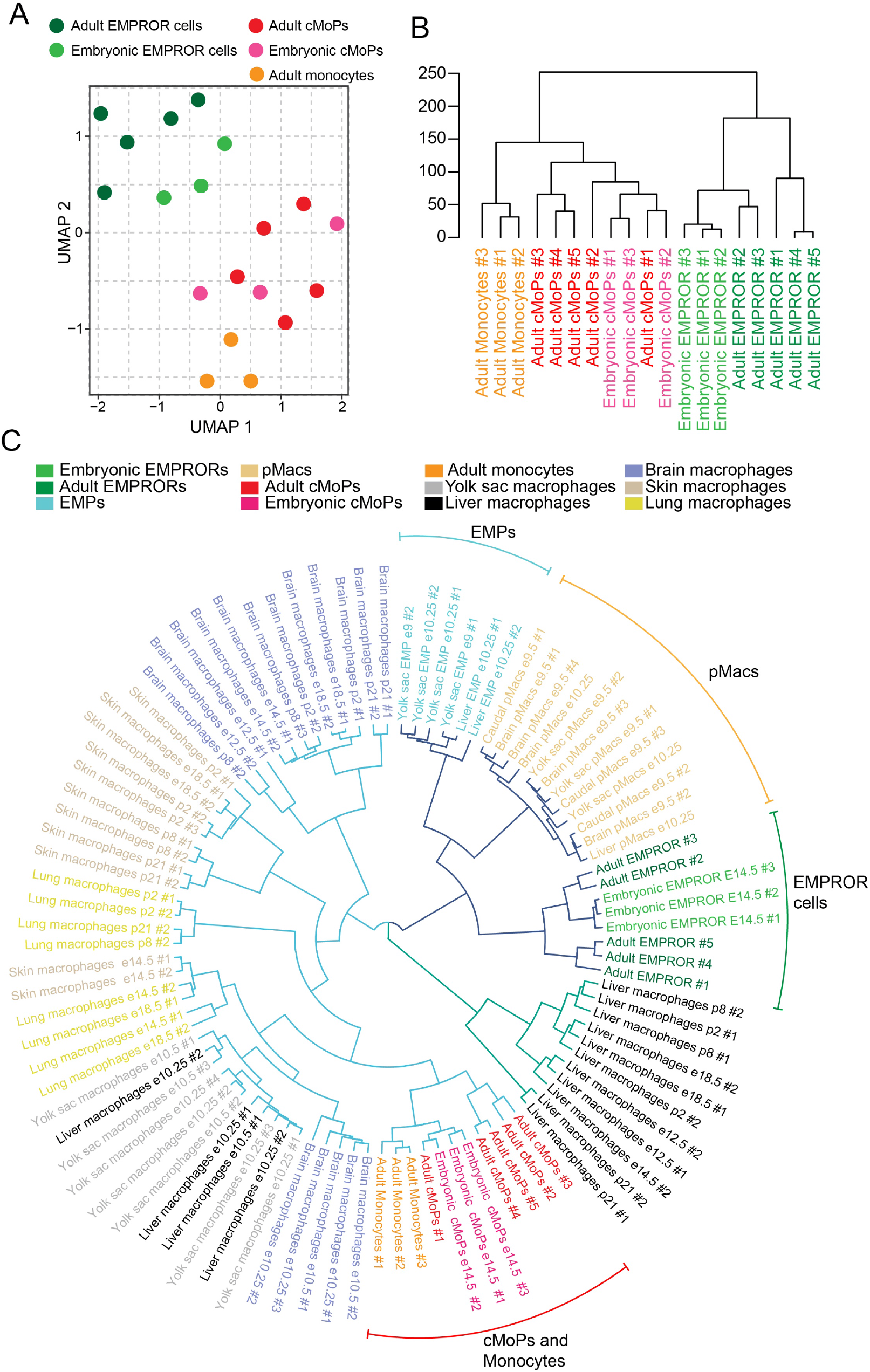
Transcriptional profiling of embryonic and adult EMPROR cells. **A**, Embryonic EMPROR cells and cMoPs were sorted by flow cytometry from pooled foetal livers of DPE-GFP:mT/mG mice at E14.5 (n=6-9 embryos/sort). Monocytes, cMoPs and EMPROR cells were sorted from adult bone marrow. Each symbol is representative of an independent sample. UMAP plot of bulk RNA-Seq of the individual cell populations shows the close association between eEMPROR and adult EMPROR cells. **B**, Hierarchical clustering of bulk RNA-Seq of cell populations described in A; **C,** Hierarchical clustering of RNA-Seq data based on top 1000 genes as determined by cell-type based variance between cells populations described in (A) and classical EMP, pMacs and, pre- and post-natal tissue macrophages. Data for classical EMP, pMacs, pre- and post-natal tissue macrophages were obtained from a previous study^3^. DeSeq2 normalised counts from both studies were combined, scaled and centered. Batch effects were removed using ComBat batch removal procedure^4^. The data shows the clustering of adult and embryonic EMPROR cells, in proximity to classical EMP and pMacs^3^. Also note that monocytes and cMoPs cluster with tissue macrophages.

In order to determine the relationship of EMPROR cells to other known embryonic macrophage precursors, such as EMP and pMacs, as well as mature macrophage populations in the embryo and adult organism, we performed unbiased comparisons of transcriptomes available in the literature (*43*). Strikingly, in this large dataset of distinct cell populations, adult EMPROR and eEMPROR cells clustered adjacent to each other and in proximity to other embryonic precursors (**Fig. 6C**). Embryonic and adult cMoPs, on the other hand, were distinct from EMPROR cells and rather clustered with other macrophage populations (**Fig. 6C**). We next identified the top 100 genes most uniformly expressed in adult and embryonic EMPROR cell populations. This set of genes also hightlighted certain similarities in the gene expression profiles of EMPROR cells, EMP and pMacs (**Suppl. Fig. 13B**). Gene ontogeny analysis of these top 100 genes depicted enrichment of several biological pathways, including those involed in haematopoiesis and leukocyte development (**Suppl. Fig. 13C**). Furthermore, comparative analysis of adult and embryonic EMPROR cells with the “pseudo-bulk”sc-RNA-seq dataset of mouse haematopoietic stem cell and progenitor cell differentiation (*45*), highlighted the presence of three independent cell population (**Suppl. Fig. 14A-C**). Notably, embryonic and adult EMPROR cells clustered together to form an independent branch within the dendrogram (**Suppl. Figure 14A**). Haematopoietic stem cell populations and multipotent progenitors formed an independent branch, while cMoPs and monocytes alongside megakaryocyte-erythrocyte progenitors (MEP), common myeloid progenitors (CMP) and granulocyte monocyte progenitors (GMP) (**Suppl. Figure 14A**) formed the third branch. UMAP analysis corroborated a similar clustering pattern of the cell populations, with EMPROR cells forming a distinct cluster (**Suppl. Figure 14B**). Similar to previous analysis (**Suppl. Fig. 13A-C**), EMPROR cells demonstrated the presence of a gene signature distinct from the known mouse hematopoietic stem and progenitor cell lineages (**Suppl. Figure 14C**). We therefore conclude that embryonic and adult EMPROR cells exhibit closely related transcriptional programs distinct from the previously described monocytic precursors.

## Concluding Remarks

One of the central tenets of macrophage development is the monocytic origin of macrophages involved in inflammation (*46*) and cancer (*47–49*) in the adult. However, the infiltrating monocytic origin of TAMs has recently been challenged (*50–52*). Here, we identify and characterise a potent, distinct tissue-resident macrophage progenitor - the EMPROR cell. In contrast to other recently described macrophage progenitor populations (*6, 7, 39*), EMPROR cells do not arise within the yolk sac but rather in a distinct extraembryonic site (MIXER) which has not previously been demonstrated to possess embryo-derived hematopoietic activity. Unlike cMoPs and monocytes, the EMPROR cells do not harbour dendritic cell but rather erythroid, neutrophil, megakaryocytic and macrophage potential. The fact that these cells appear to be distinct from the classical monocytic lineage, as determined by phenotypic marker expression, transcriptome analysis and lineage potential *in vitro* and *in vivo*, indicates that at least two parallel systems with macrophage differentiation potential exist within the adult BM. This could represent a failsafe mechanism ensuring the production of essential myeloid immune cells in the case of impairment of one lineage, for example, during infections or under conditions of limited growth factors. Alternatively, monocytes and EMPROR cells may generate distinct macrophage populations, which is supported by the finding that GFP^+^ PVM detected in DPE-GFP transgenic mice only derive from EMPROR cells. Future studies will define the precise transcriptional and functional profile, and subset-specific contribution of monocyte- and EMPROR-derived macrophages to various tissues as well as under different disease settings.

The presence of an alternative site of haematopoiesis at the maternal-foetal interface posits the question as to how haematopoiesis begins at this site. We currently do not know whether blood cell development in MIXER and yolk sac results in qualitative differences between the developing cell lineages. Nonetheless, the presence of haematopoietic progenitors within MIXER tissue at an early stage suggests that this tissue might be more intimately involved in shaping the intraembryonic environment than previously assumed. The cells and factors involved in the generation of haematopoietic cells in the MIXER await further characterisation.

Lastly, our demonstration of a distinct transcriptional program that is shared between eEMPROR cells and adult EMPROR cells strongly argues for their close relationship and may indicate that these cells are maintained from the earliest time of haematopoietic development through to adulthood, albeit in different body sites. Nevertheless, we cannot exclude that HSC in the BM partially or exclusively generate EMPROR cells after birth. EMPROR cell-specific lineage tracing models would be needed to address this question. However, currently, no suitable lineage tracing mouse lines exist in this regard; available lines, such as Csf1r^Mer-iCre-Mer^Rosa^LSL-YFP^ (*53*) and Cx3cr1^CreER^Rosa26^YFP^ (*19, 54*) utilise markers that are highly expressed in classical EMPs when compared to EMPROR cells. Further, existing CD4-cre lines (*55*) use different constructs compared to the DPE-GFP line resulting in transgene expression in different cell types. Therefore, new lineage tracing mouse lines will have to be constructed in the future to address the ontogeny of adult EMPROR cells and their progeny. Regardless, based on their clear existence during development and in the adult, we postulate that EMPROR cells hold a separate, functionally unique place within the macrophage realm and complement the monocyte lineage in the generation of tissue macrophages in the adult.

## Acknowledgments

We thank M. Rizk, J. Qin, A. Cooray, K. Bremner, L. Shaw, Q. Lee and R. Kwan for animal husbandry and genotyping. We thank the Core Flow Cytometry and Imaging Facility at the Centenary Institute especially F. Kao, C. Royle, A. Aspland, S. Allen, T.M. Ashhurst, K. Jahn, D. Liu and A. Smith for their outstanding technical support during the course of this project. We would also like to thank the Australian Genome Research Facility (AGRF), Melbourne, Victoria and Ramaciotti Centre for Genomics (UNSW), Sydney for performing Next Generation Sequencing. We thank L. Cavanagh for administrative assistance. We thank the outstanding support of our animal facility staff especially R. Barugahare, C. Stuart, N. Littlejohn, D. Moyes, K. Hickey, C. Juaton, S. Ilufi, B. Etheridge, M. Murarotto and H. Golan for assistance with timed-mating experiments. We thank B. Roediger, C. Jolly, P. Bertolino, and F. Sierro for helpful discussions. We thank S. Naik for providing OP9 and OP9-DL1 cells and Prof. R. Anderson for providing E0771 breast adenocarcinoma cells. We thank M. Guilliams (VIB) for kindly providing the scripts for the generation of hexagonal and rose diagrams. We thank N. Pearce, N. Keilar, J. Crosbie, A. Campbell and N. Yung for administrative assistance. We acknowledge the assistance of Cure Cancer (Cancer Australia) PdCCRS funding, University of Sydney ECR support and Centenary Future Fellowship support to ST. S.J. was supported by the European Research Council (Adv ERC 340345). WW and RJ were supported by grant 1126403 from National Health and Medical Research Council, Australia. WW was supported by a grant from the Vienna Science and Technology Fund.

## Conflict of Interest

Authors declare no conflict of interest.

## Data availability

Data generated in this study are available from the corresponding authors upon request. All RNA sequencing datasets will be made publicly available as GEO datasets.

## Author Contributions

ST, RJ and WW designed and supervised the study. ST and RJ performed experiments and analysed data. ST, RJ and BM carried out the flow cytometry and sorts, lineage analysis, bone-marrow chimera studies, RNA preparation and RT PCR. ST and RJ carried out confocal and intravital microscopy experiments. METP assisted with tissue preparation for confocal microscopy. LS assisted with bone-marrow studies. SR, FL, RS, JJLW, MW and MF performed bioinformatic analyses. STF and RJ performed embryo dissections. STF, RJ, BM and ST carried out embryo analysis. SJ, JJLW and STF provided valuable technical and intellectual input. AJM provided important technical input. ST, RJ and WW wrote the manuscript. SJ and STF provided assistance with editing the manuscript. All authors have read and approved the manuscript.

---

**Supplementary Figure 1:**
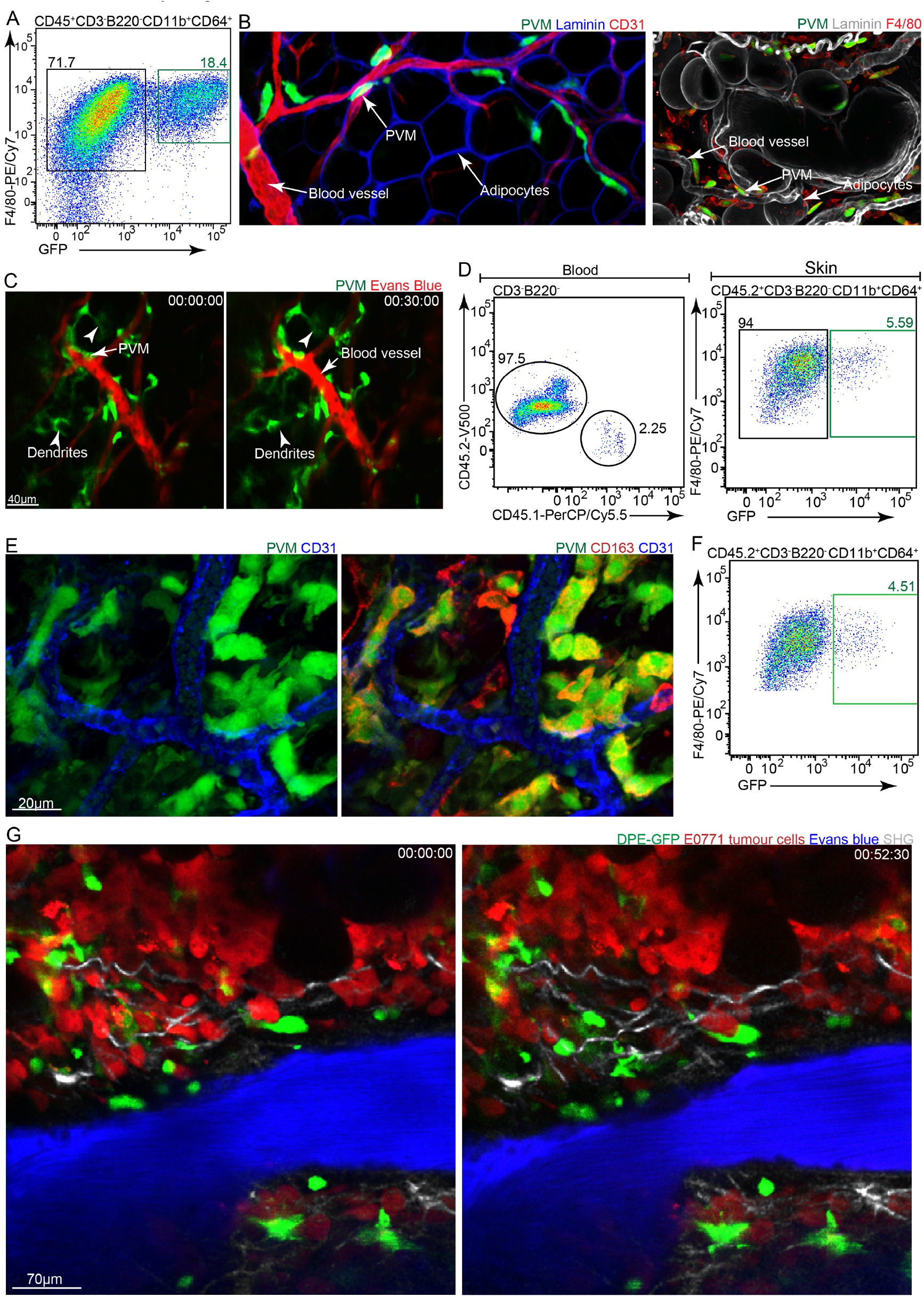
Identification of GFP^+^ perivascular macrophages in murine mammary fat pads and mammary tumours. **A**, Flow cytometric analysis identifying CD45^+^CD3e^-^B220^-^CD11b^+^CD64^+^F4/80^+^GFP^-^ and GFP^+^ macrophages within mammary fat pads of naïve DPE-GFP:RAG-1^-/-^ mice. Representative flow plot from one of three independent experiments is shown; **B**, Confocal imaging of immunostained mammary fat pad identifies GFP^+^ perivascular macrophages. Co-staining with F4/80 is depicted in the right panel. Image data is representative of tissues harvested from n=3 mice from three independent experiments; **C**, Intravital imaging of sessile GFP^+^ perivascular macrophages displaying actively scanning dendrites. Data represents images from three independent experiments; **D**, Flow cytometric analysis of the ear skin from DPE-GFP:RAG-1^-/-^→PTPRCA chimeras demonstrating partial replacement of CD45.2^+^ CD3e^-^B220^-^CD11b^+^CD64^+^F4/80^+^ GFP^+^ macrophages within the tissue over 15 months, n=3; **E**, Confocal imaging of immunostained mammary tumours shows the presence of numerous GFP^+^ perivascular macrophages along neoangiogenic tumour vasculature. Data represents images from three independent experiments; **F**, Flow cytometric analysis identifying CD45^+^CD3e^-^B220^-^ CD11b^+^CD64^+^F4/80^+^GFP^-^ and GFP^+^ macrophages within mammary tumours. Representative flow plot from one of three independent experiments is shown; **G,** Intravital imaging of E0771 mammary tumours transplanted into DPE-GFP:RAG-1^-/-^ mice demonstrates the presence of GFP^+^ PVM around the tumour vasculature. Image data representative of intravital studies with n=3 mice from three independent experiments.

**Supplementary Figure 2:**
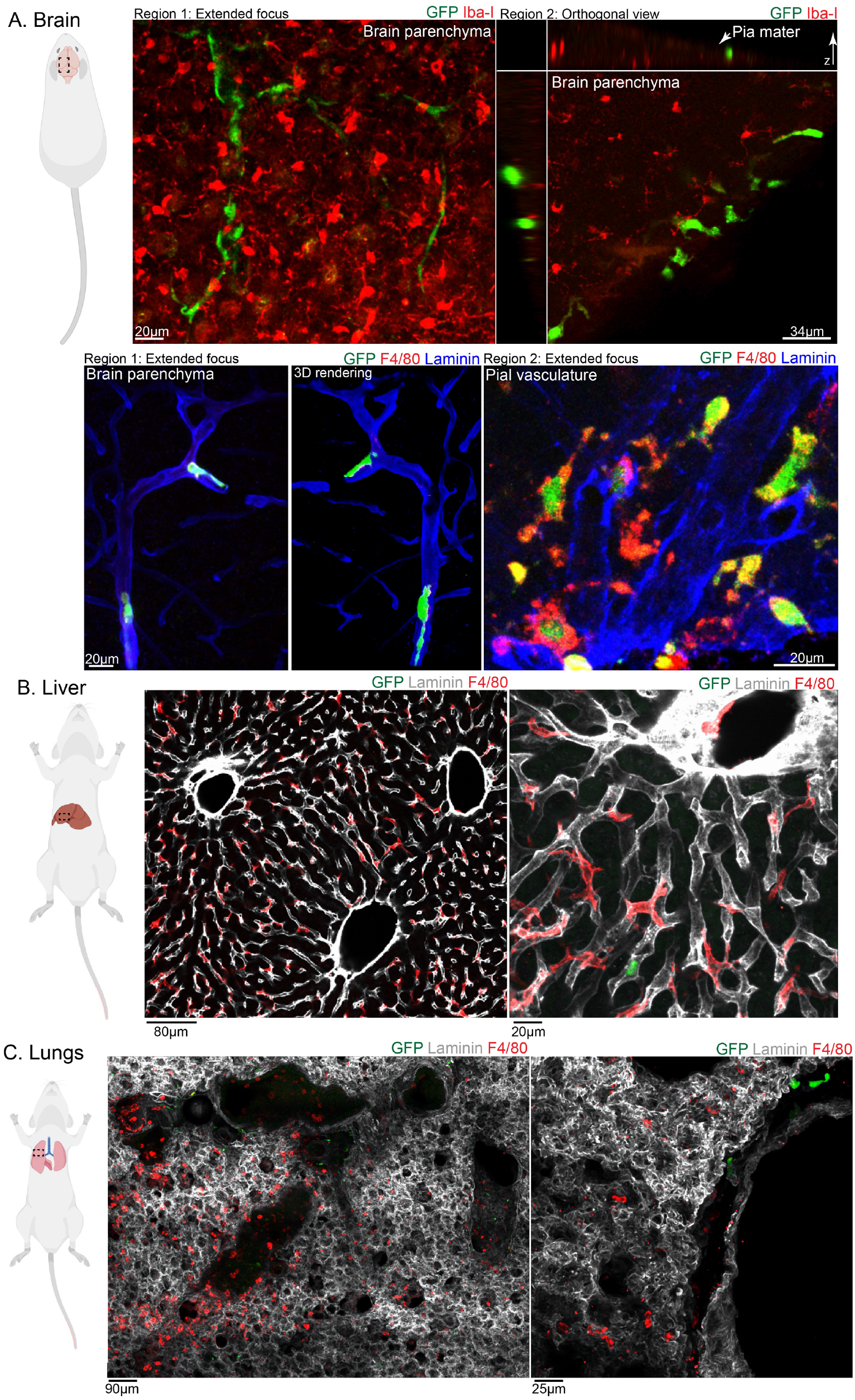
GFP^+^ perivascular macrophage distribution in various murine tissues. **A**, Confocal imaging of immunostained brain sections from DPE-GFP:RAG-1^-/-^ mice showing the absence of GFP expression in microglial cells (marked by Iba-I expression) and distinct expression of GFP in perivascular macrophages along the pial vasculature; **B**, Confocal imaging of immunostained liver sections from DPE-GFP:RAG-1^-/-^ mice reveals the absence of GFP expression in Kupffer cells; **C**, Confocal imaging of immunostained lung sections from DPE-GFP:RAG-1^-/-^ mice highlighting the absence of GFP expression in lung macrophages. Pane A-C, Image data is representative of tissue harvested from n=3 mice from two independent experiments.

**Supplementary Figure 3:**
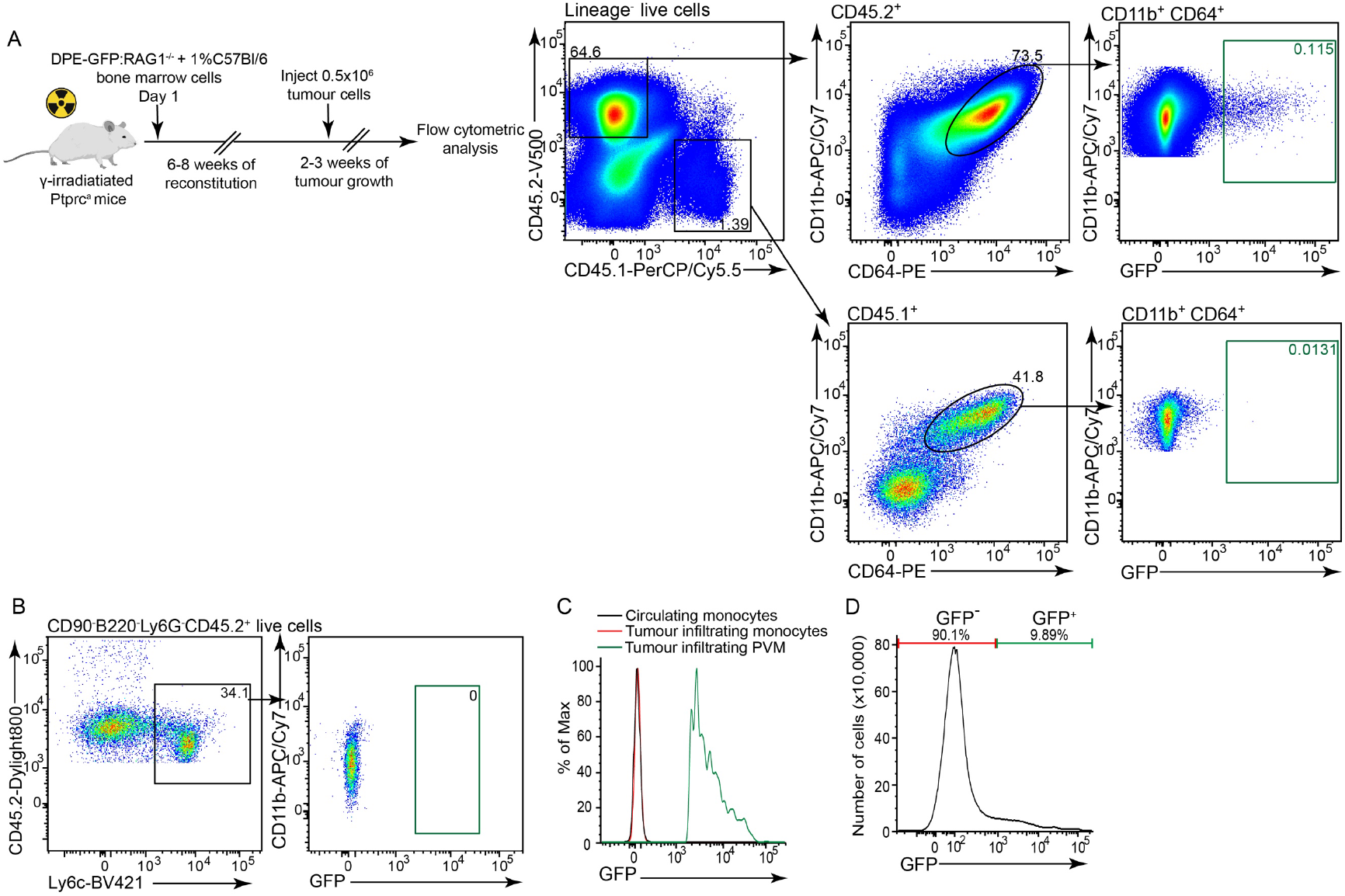
Origin of GFP^+^ macrophages post-inflammation. **A**, Flow cytometric analysis of tumours from DPE-GFP:RAG-1^-/-^→PTPRCA chimeric mice identifying the presence of donor-derived GFP^+^ and GFP^-^ macrophages; n=4 mice. Data is representative of one of two independent experiments. Lineage markers consist of CD90 and B220. CD45.1 and CD45.2 negative cells represent tumour cells; **B**, Flow cytometric analysis of blood from DPE-GFP:RAG-1^-/-^→PTPRCA chimeric mice highlighting the absence of GFP in donor-derived monocytes. n=4 mice. Data is representative of one of two independent experiments; **C**, Histogram plot delineating the expression of GFP in circulating and tumour infiltrating monocytes, tumour infiltrating GFP^+^ PVM were used as an internal positive control for GFP expression; n=4 mice. Data is representative of one of two independent experiments. **D**, Histogram plot displaying the presence of GFP^+^ and GFP^-^ cells within the DPE-GFP bone marrow. Histogram plot is representative of several independent experiments (n > 5).

**Supplementary Figure 4:**
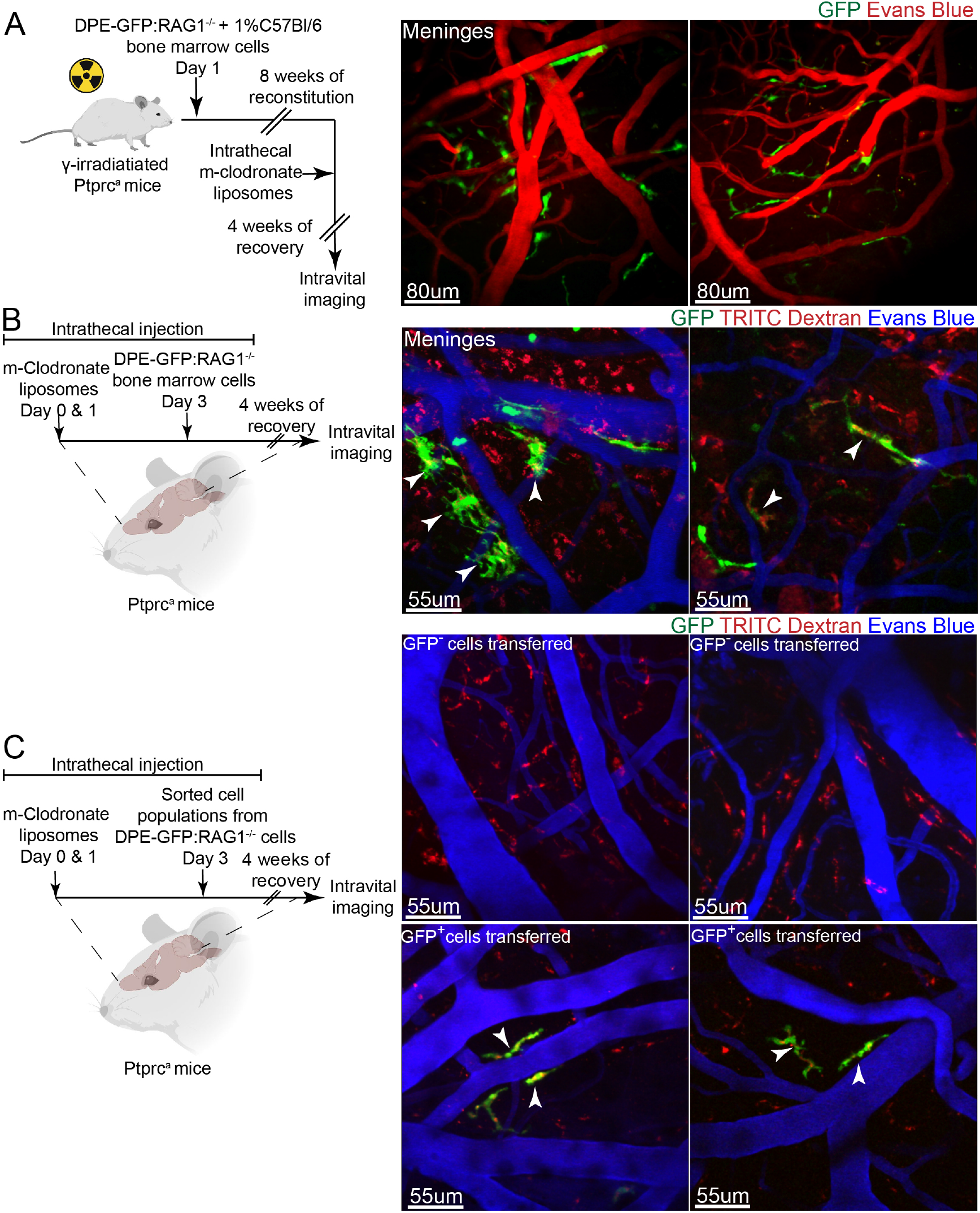
Reconstitution of perivascular macrophages in the CNS. **A**, Intravital imaging of DPE-GFP:RAG-1^-/-^→PTPRCA chimeric mice 4 weeks post-depletion of macrophages by mannosylated clodronate liposomes identifies the presence of numerous donor-derived GFP^+^ perivascular macrophages in the CNS. Image data is representative of n=4 mice from four independent chimera experiments; **B**, Intravital imaging of brain from mice injected with unfractionated DPE-GFP:RAG-1^-/-^ bone marrow cells post-meningeal macrophage depletion identifies the presence of several GFP^+^TRITC dextran^+^ PVM. This experiment establishes the intrathecal precursor transfer methodology used in subsequent experiments. Image data is representative of n=3 mice from two independent experiments; **C**, Intravital imaging of brain from mice injected with sorted GFP^-^ and GFP^+^ bone marrow cells from DPE-GFP:RAG-1^-/-^ mice. GFP^+^ perivascular macrophages were only detected in mice injected with GFP^+^ bone marrow cells. Image data is representative of three independent experiments wherein the CNS was repleted with GFP^+^ (n=4) and GFP^-^ (n=3) sorted bone marrow cells from DPE-GFP:RAG-1^-/-^ mice.

**Supplementary Figure 5:**
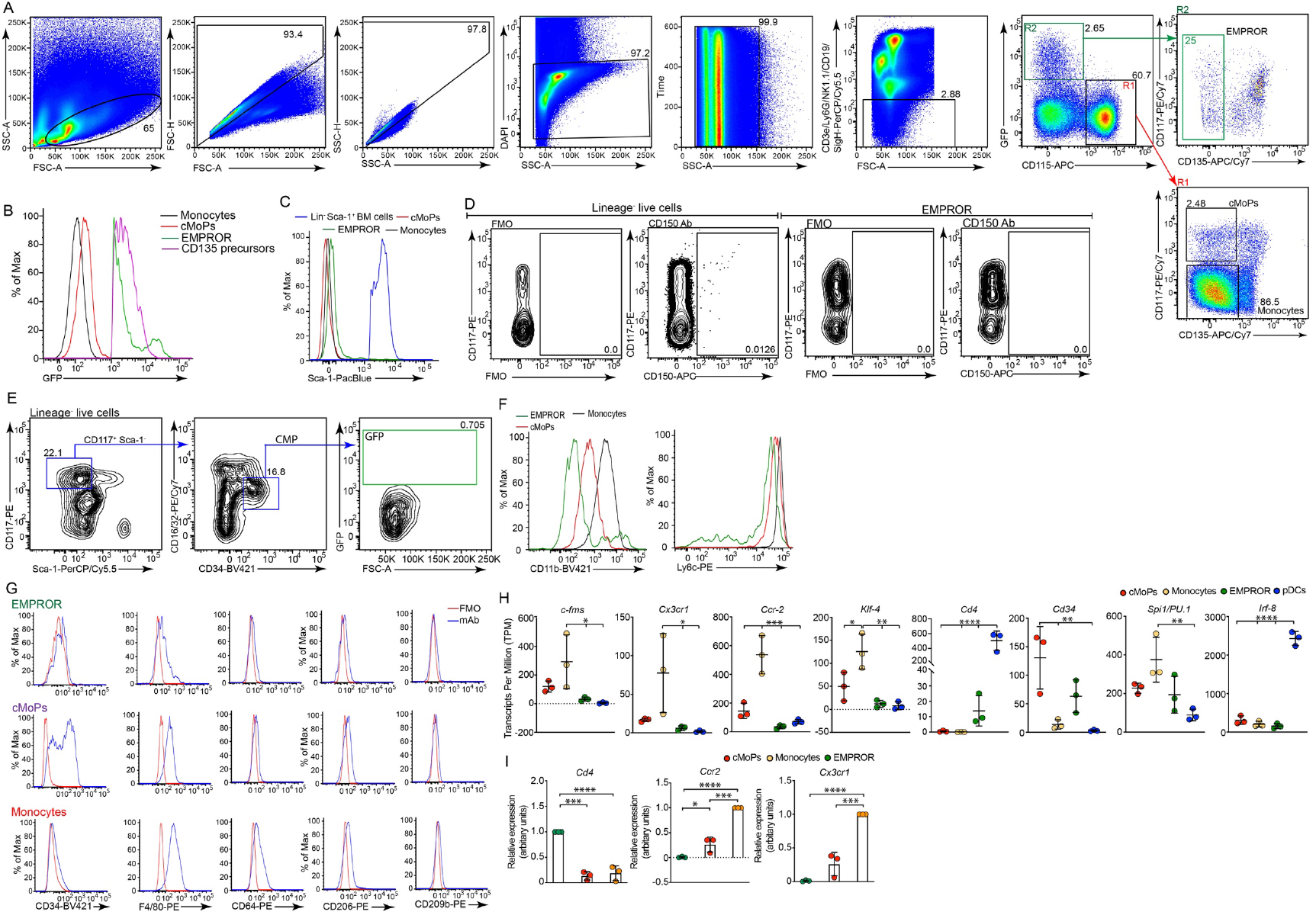
Characterisation of EMPROR cells. **A,** Flow cytometric analysis of bone marrow from DPE-GFP mice identifies EMPROR cells as SSC^lo^DAPI^-^ CD3e^-^Ly6G^-^NK1.1^-^CD19^-^SiglecH^-^CD115^-^CD135^-^GFP^+^CD117^+^ cells. Representative flow plots from one of several independent experiments (n>5) are shown; **B**, Histogram plots depicting the expression of GFP in cMoPs, monocytes, CD135 precursor and EMPROR cells (n>5); **C**, Histogram plots showing the surface expression of Sca-1 on cMoPs, monocytes, and EMPROR. Bone marrow cells not expressing lineage markers but exhibiting surface expression of Sca-1 have been overlayed as an internal positive control. Lineage markers consist of CD3e, Ly6G, NK1.1, CD19 and Siglec-H; **D**, Dot plots representing the lack of surface expression of CD150 on EMPROR cells. Bone marrow cells not expressing lineage markers but exhibiting surface expression of CD150 have been displayed as an internal positive control. Lineage markers consist of CD3e, Ly6G, NK1.1, CD19 and Siglec-H; **E**, Flow cytometric analysis of DPE-GFP bone marrow revealing the absence of GFP expression in common myeloid progenitors^5^. Lineage markers consist of CD3e, CD4, CD8, B220, IgM, Ter119, CD127, CD19 and Gr-1^5^; **F**, Histogram plots demonstrating the expression of CD11b and Ly6C by cMoPs, monocytes and EMPROR within the adult bone marrow; **G**, Histogram plots displaying the expression of CD34, F4/80, CD64, CD206 and CD209b on cMoPs, monocytes and EMPROR cells. Representative flow plots from one of three independent experiments are shown in panel C-G. Note that for panel A, F and G the lineage cocktail comprised of CD3e, Ly6G, NK1.1, CD19 and Siglec-H antibodies; **H**, Transcriptomic analysis of various genes associated with myeloid cell development and function emphasising the unique transcriptional signature of EMPROR cells when compared to cMoPs and monocytes. The data was analysed from the sequencing of sorted precursors from three independent sort experiments (n=5-7 mice/sort); **I**, qRT-PCR analysis confirming the differential expression of *Cd4*, *Ccr2* and *Cx3cr1* mRNA in EMPROR cells compared to cMoPs and monocytes. The data was generated from qRT-PCR of sorted precursors from three independent sort experiments (n=4-5 mice/sort). One-way ANOVA was employed; ****P ≤0.0001, ***P ≤0.001, **P ≤0.01, *P ≤0.1. Bars represent mean ± S.D. S.D.-Standard deviation.

**Supplementary Figure 6:**
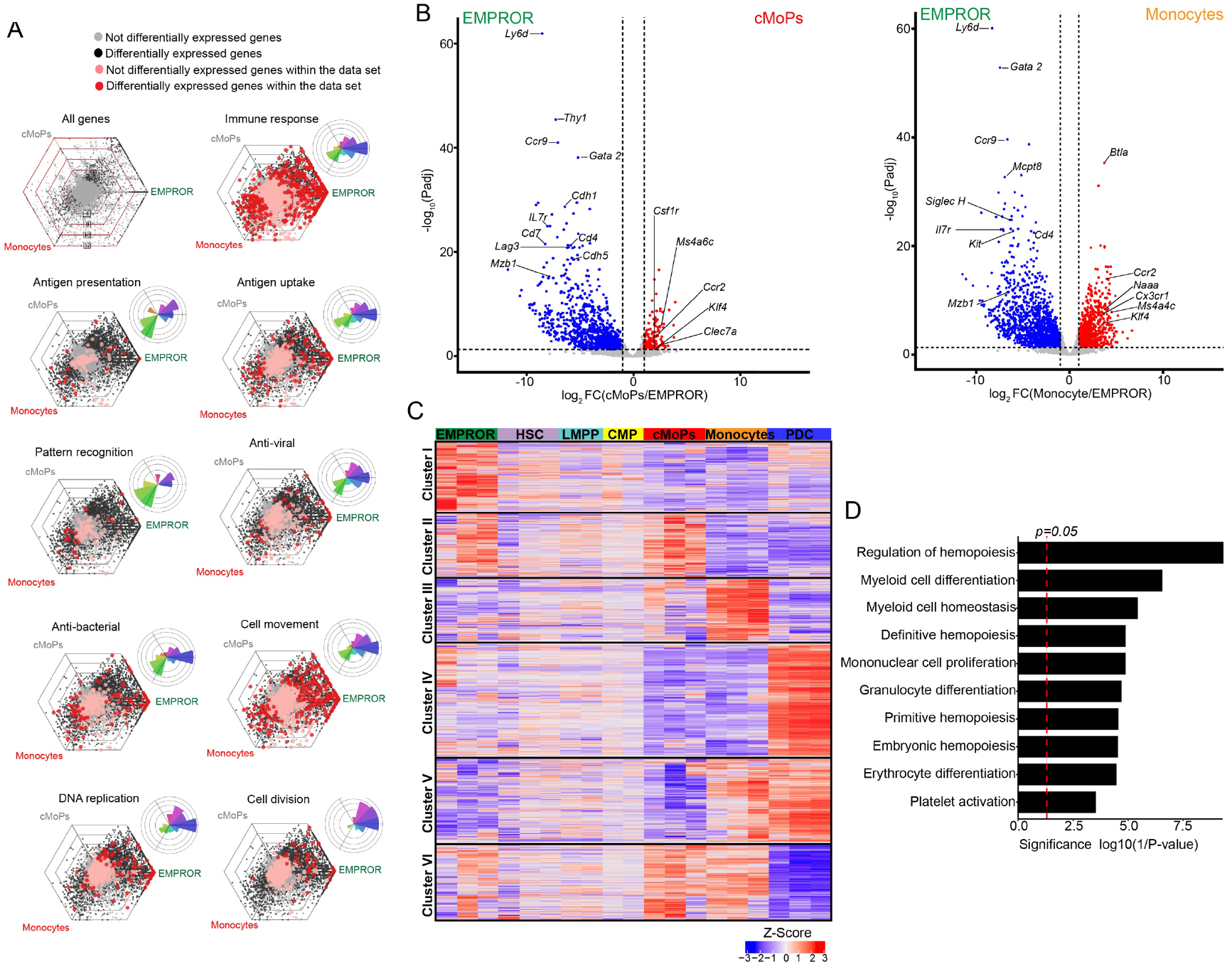
Transcriptional analysis of EMPROR cells. **A**, Hexagonal and rose diagrams highlighting the unique gene expression profile of EMPROR cells when compared to cMoPs and monocytes; **B**, Volcano plots depicting select differentially expressed genes between EMPROR cells and cMoPs and monocytes; **C**, K-means clustering analysis of differentially expressed genes (*P* <0.05 after Benjamini-Hochberg correction, fold-change > 2) in HSC, LMPP, CMP, cMoPs, monocytes, pDC and EMPROR cells identifies 6 unique clusters, with cluster I being specific for EMPROR cells. Expression values for each differentially expressed gene are normalised across all samples by Z-score transformation method; **D**, Gene ontology analysis of cluster I depicts the significant representation of functions involved in erythroid and myeloid haematopoiesis.

**Supplementary Figure 7:**
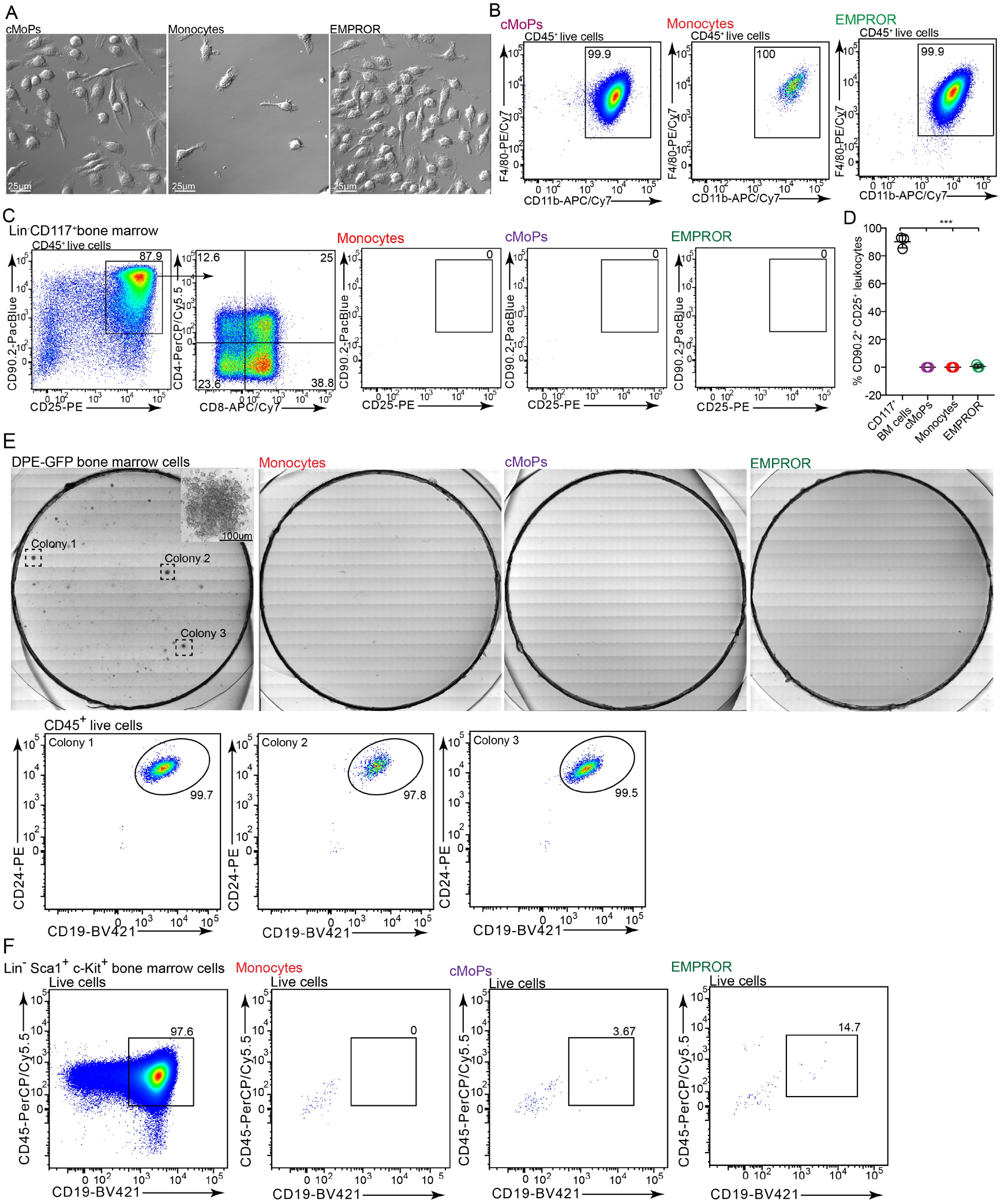
*In vitro* differentiation potential of adult EMPROR cells. **A**, Confocal differential interference contrast (DIC) imaging of cells following *in vitro* culture of sorted cMoPs, monocytes and EMPROR cells in RPMI containing M-CSF. Representative images from one of two independent experiments; **B**, Flow-cytometric analysis of cultured cells identifying the presence of CD45^+^CD11b^+^F4/80^+^ macrophages. Representative flow plot from one of three independent experiments; **C**, Flow cytometric analysis of sorted cell populations from the bone marrow of DPE-GFP mice post-culture on OP-9 and OP-9 DL1 stromal cells corroborates the limited potential of the precursors to differentiate into T cells and their precursors. Lin^-^CD117^+^ bone marrow cells were used as a positive control. Representative flow plots from one of three independent experiments; **D**, Graph representing the lack of T cell potential of sorted cMoPs, monocytes and EMPROR cells from DPE-GFP bone marrow when cultured employing OP-9 and OP-9/DL-1 system. Sorted lin^-^CD117^+^ bone marrow cells were used as a positive control. Data are cumulative of three independent experiments. Bars represent mean ± S.D.; **E**, Confocal imaging and flow cytometry of sorted cell populations from the bone marrow of DPE-GFP mice post-culture on B cell MethoCult™ depicts the limited potential of the precursors to differentiate into B cells. Unfractionated bone marrow cells were utilised as a positive control. Representative flow plot of 3 colonies from one of three independent experiments; **F**, Flow cytometric analysis of sorted cell populations from the bone marrow of DPE-GFP mice post-culture on OP-9 stromal cells illustrates the limited potential of the precursors to differentiate into B cells or their precursors. Lin^-^Sca-1^+^c-Kit^+^ bone marrow cells were employed as a positive control.

**Supplementary Figure 8:**
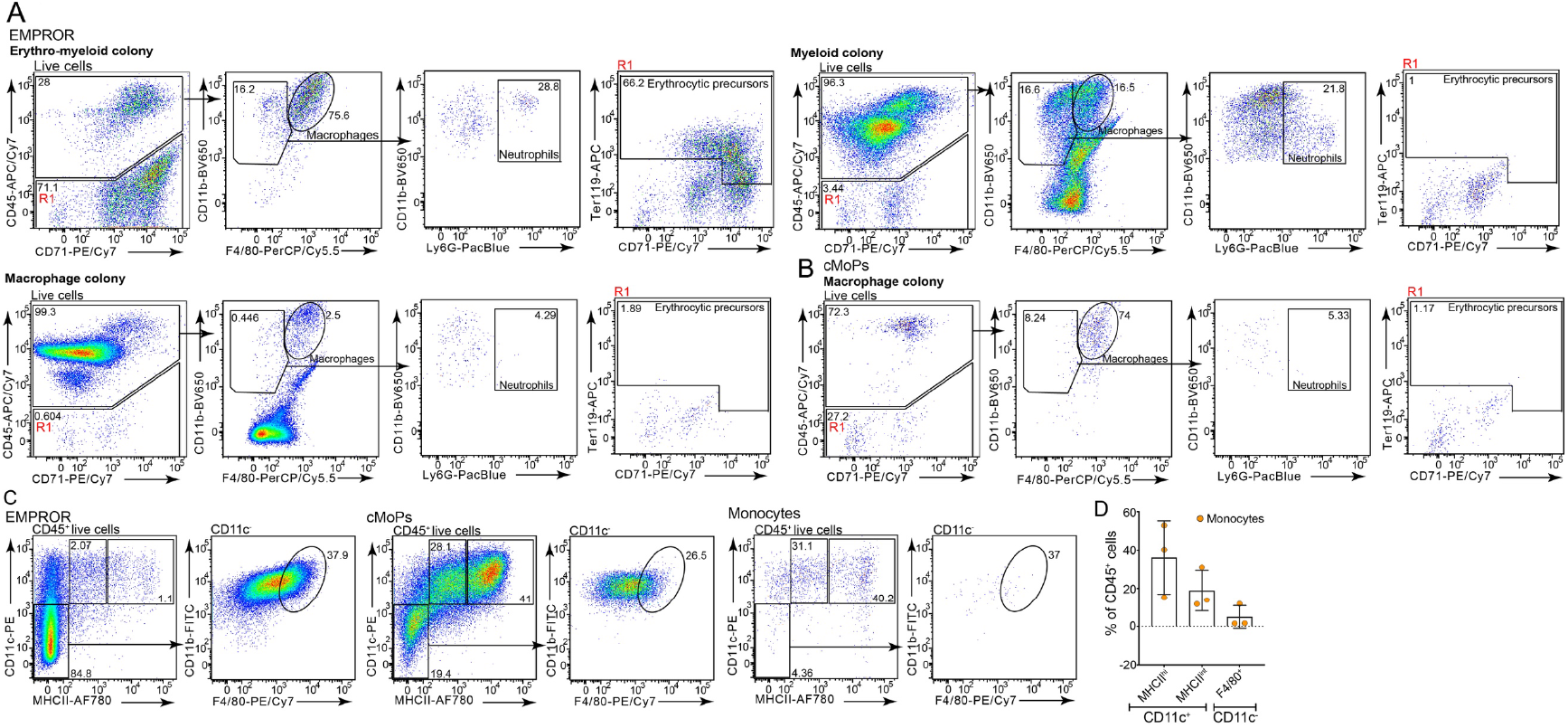
*In vitro* erythro-myeloid and dendritic cell differentiation potential of EMPROR cells. **A**, Flow-cytometric analysis of colonies obtained post-culture of sorted cell populations in MethoCult™ GF M3434 containing transferrin, SCF, IL-3, IL-6, EPO, and supplemented with TPO, GM-CSF and M-CSF. Flow plots highlight colonies derived from EMPROR cells with erythro-myeloid, myeloid and macrophage potential. Erythroid cells were defined as CD45^-^ Ter119^+^CD71^+/hi^, neutrophils as CD45^+^CD11b^+/hi^F4/80^-^Ly6G^+^ and macrophages as CD45^+^CD11b^+^F4/80^+^ cells. Flow plots of three colonies from one of three independent sort and culture experiments are shown. A total of 34 independent colonies were analysed; **B**, cMoP colonies were restricted to macrophage potential only. Representative flow plots from one of three independent sort and culture experiments are shown; **C**, Flow-cytometric analysis depicting the development of CD11c^+^MHCII^+^ and CD11c^+^MHCII^int^ dendritic cells, and CD11c^-^MHCII^-^CD11b^+^F4/80^+^ macrophages obtained from *in vitro* differentiation of sorted EMPROR cells, cMoPs and monocytes in the presence of 20 ng/ml GM-CSF. Flow-plots are representative of one of three independent sort and culture experiments; **D**, Quantification of % of CD11c^+^MHCII^+^ and CD11c^+^MHCII^int^ dendritic cells, and CD11c^-^MHCII^-^CD11b^+^F4/80^+^ macrophages obtained from *in vitro* differentiation of sorted monocytes. Data are cumulative of three independent sort and culture experiments. Bars represent mean ± S.D. S.D.-Standard deviation.

**Supplementary Figure 9:**
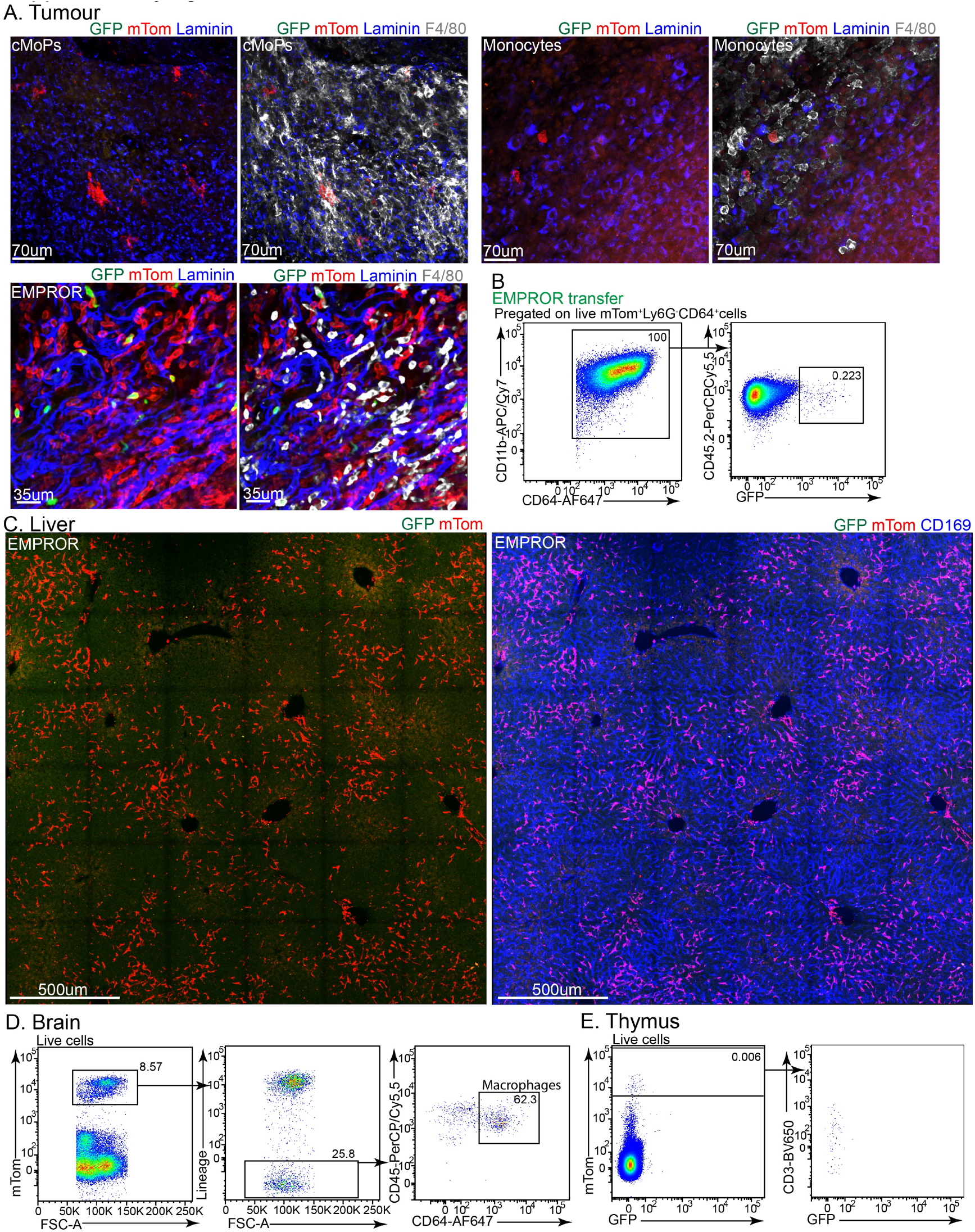
*In vivo* lineage potential of EMPROR cells. **A**, Confocal imaging of immunostained E0771 tumour section obtained from mice reconstituted with cMoPs, monocytes or EMPROR cells shows the presence of mTom^+^F4/80^+^ cells. Images are representative of one of three independent sort, adoptive transfer and tumour implantation experiments; **B**, Flow cytometric analysis of tumour obtained from chimeric mice reconstituted with EMPROR cells identifies GFP^+^ macrophages within the tumour. Representative flow plots from one of six independent sort, adoptive transfer and tumour implantation experiments are shown; **C**, Confocal imaging of immunostained liver section obtained from mice reconstituted with EMPROR cells demonstrates the presence of numerous mTom^+^CD169^+^ cells (large field of view). Images are representative of one of three independent sort, adoptive transfer and tumour implantation experiments; **D**, Flow cytometric analysis of brain from chimeric mice reconstituted with EMPROR cells identifies mTom^+^ macrophages within the tissue. Representative flow plots from one of three independent sort, adoptive transfer and tumour implantation experiments are shown; **E**, Flow cytometric analysis of thymus from chimeric mice reconstituted with EMPROR cells depicts the absence of mTom^+^ T cells within the tissue. Representative flow plots from one of six independent sort, adoptive transfer and tumour implantation experiments are shown.

**Supplementary Figure 10:**
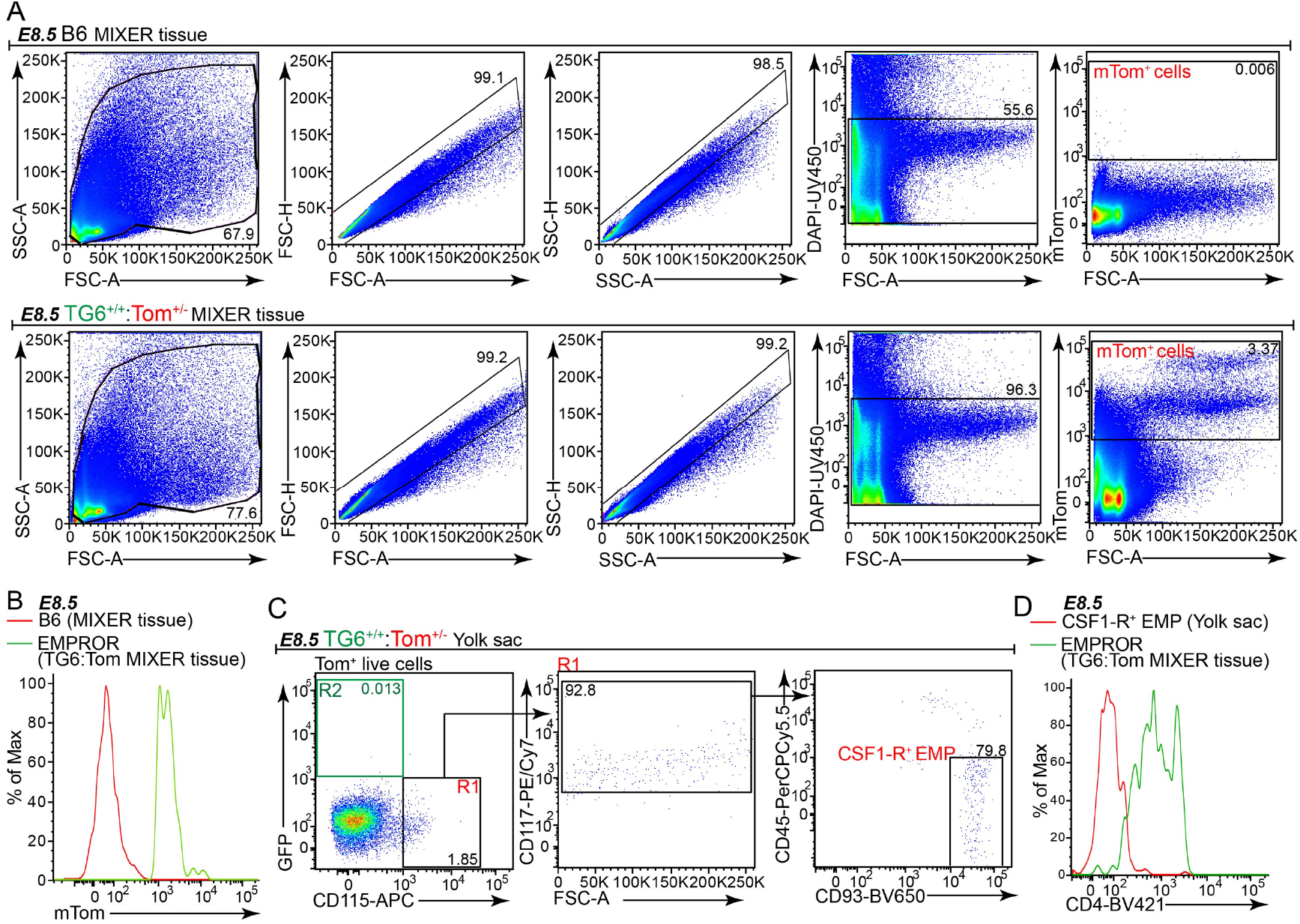
Identification of embryo-derived cells at the maternal-embryo interface. **A**, Flow-cytometric analysis of MIXER tissue isolated from either C57BL/6 or DPE-GFP^+/+^:mTom^+/-^ concepti depicting the gating strategy for mTom^+^ cells. Representative flow plots from n=5 independent litters (6-11 embryos/litter) from five separate timed-mating pairs from three independent experiments are shown. The embryos were staged as E8.5 with 6-9 somite pairs; **B**, Histogram plots indicating the expression of mTom by EMPROR cells identified in the MIXER tissue of DPE-GFP^+/+^:mTom^+/-^ concepti. Representative flow plots from n=5 independent litters (6-11 embryos/litter) from five separate timed-mating pairs from three independent experiments are shown. The embryos were staged as E8.5 with 6-9 somite pairs; **C**, Flow cytometric analysis highlighting the presence of CSF1-R^+^ EMP but not EMPROR cells within the yolk sac at E8.5; **D**, Histogram plots showing CD4 expression on CSF1-R^+^ EMP within the yolk sac as compared to MIXER tissue-derived EMPROR cells. Panel C and D, data representative of n=4 independent litters (6-11 embryos/litter) from four separate timed-mating pairs from two independent experiments. The embryos were staged as E8.5 with 6-8 somite pairs.

**Supplementary Figure 11:**
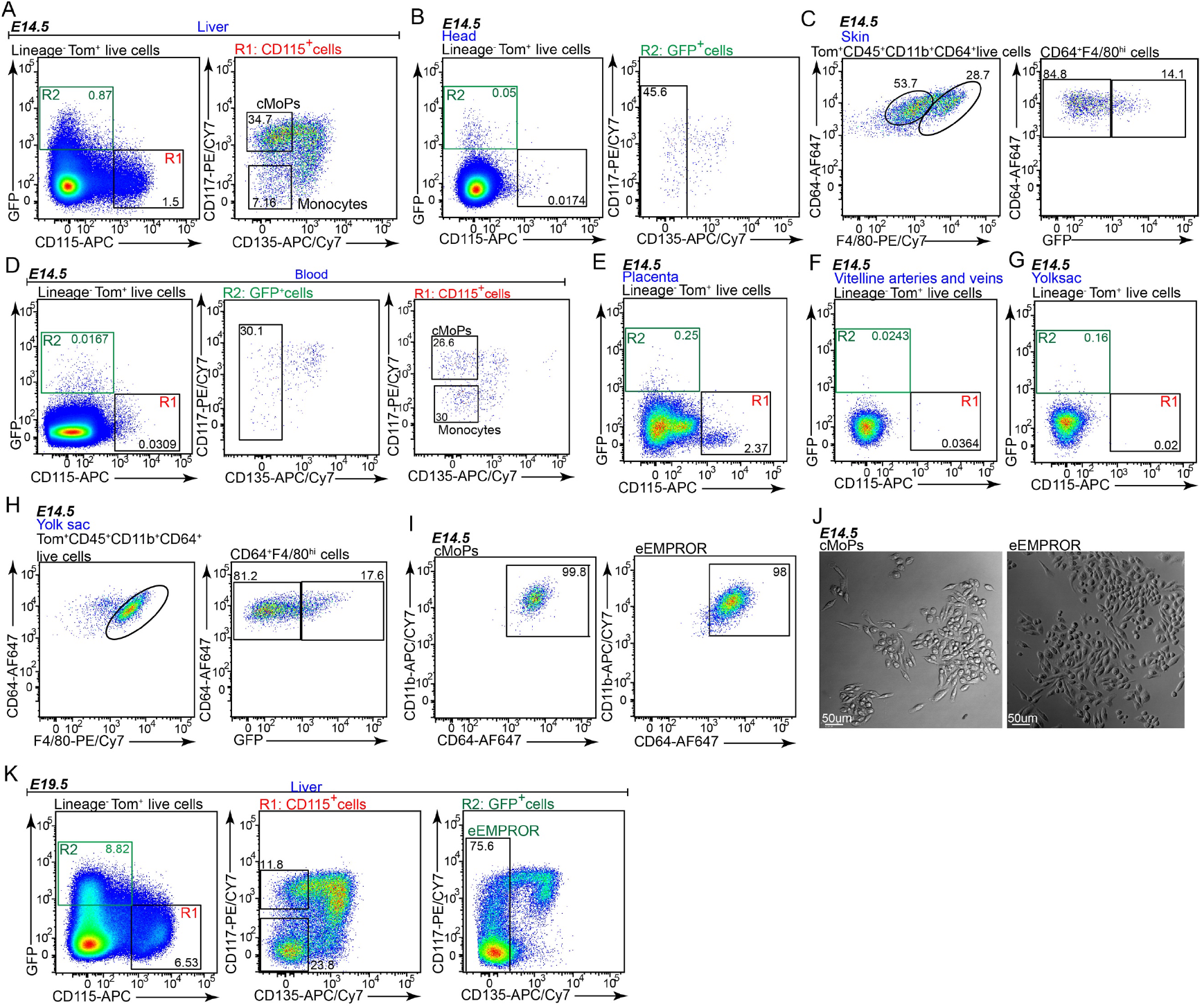
Development of embryonic EMPROR cells and macrophage precursors. **A**, Flow cytometric analysis of E14.5 foetal liver showing the presence of CD115^+^ cMoPs and monocytes. Representative flow plots from one of four independent experiments are shown; **B**, Flow cytometric analysis of E14.5 embryo head revealing the presence of eEMPROR cells. Representative flow plots from one of four independent experiments are shown; **C**, Flow cytometric analysis of foetal skin demonstrating the presence of GFP^+^ macrophages. Representative flow plots from one of two independent experiments are shown; **D**, Flow cytometric analysis of E14.5 embryonic blood highlighting the presence of eEMPROR cells, CD115^+^ cMoPs and monocytes. Representative flow plots from one of two independent experiments are shown; **E**, Flow cytometric analysis of E14.5 foetal placenta depicting the presence of few GFP^+^ precursors. Representative flow plots from one of four independent experiments are shown; **F**, Flow cytometric analysis of E14.5 vitelline arteries and veins displaying the presence of few GFP^+^ precursors. Representative flow plots from one of four independent experiments are shown; **G**, Flow cytometric analysis of E14.5 yolk sac indicating the absence of GFP^+^ precursors. Representative flow plots from one of four independent experiments are shown. Lineage markers for panel B-G consist of CD3e, Ly6G, NK1.1, CD19, Siglec-H and CD64. **H**, Flow cytometric analysis of E14.5 yolk sac detailing the presence of both GFP^-^ and GFP^+^ macrophages. Representative flow plots from one of three independent experiments are shown; **I**, Flow cytometric analysis illustrating the development of CD11b^+^CD64^+^ macrophages from sorted cMoPs and eEMPROR cells when cultured with 50 ng/ml M-CSF. Representative flow plots from one of three independent sort and culture experiments are shown; **J**, Confocal imaging of macrophages obtained from *in vitro* culture of sorted embryonic cMoPs and eEMPROR cells in RPMI medium containing M-CSF. Representative flow plots from one of two independent sort and culture experiments are shown; **K**, Flow cytometric analysis of foetal livers from E19.5 embryos exhibiting the presence of embryonic EMPROR (eEMPROR) cells. The presence of lin^-^GFP^+^CD115^-^ CD117^+^CD135^+^ and lin^-^GFP^-^CD115^+^ precursors can also be detected within the tissue. Lineage markers consist of CD3e, Ly6G, NK1.1, CD19, Siglec-H, and CD64. Representative flow plots from one of three independent experiments are shown. Flow plots presented in Panel A-H and K are from embryos harvested from independent litters from separate timed-mating pairs from independent experiments.

**Supplementary Figure 12:**
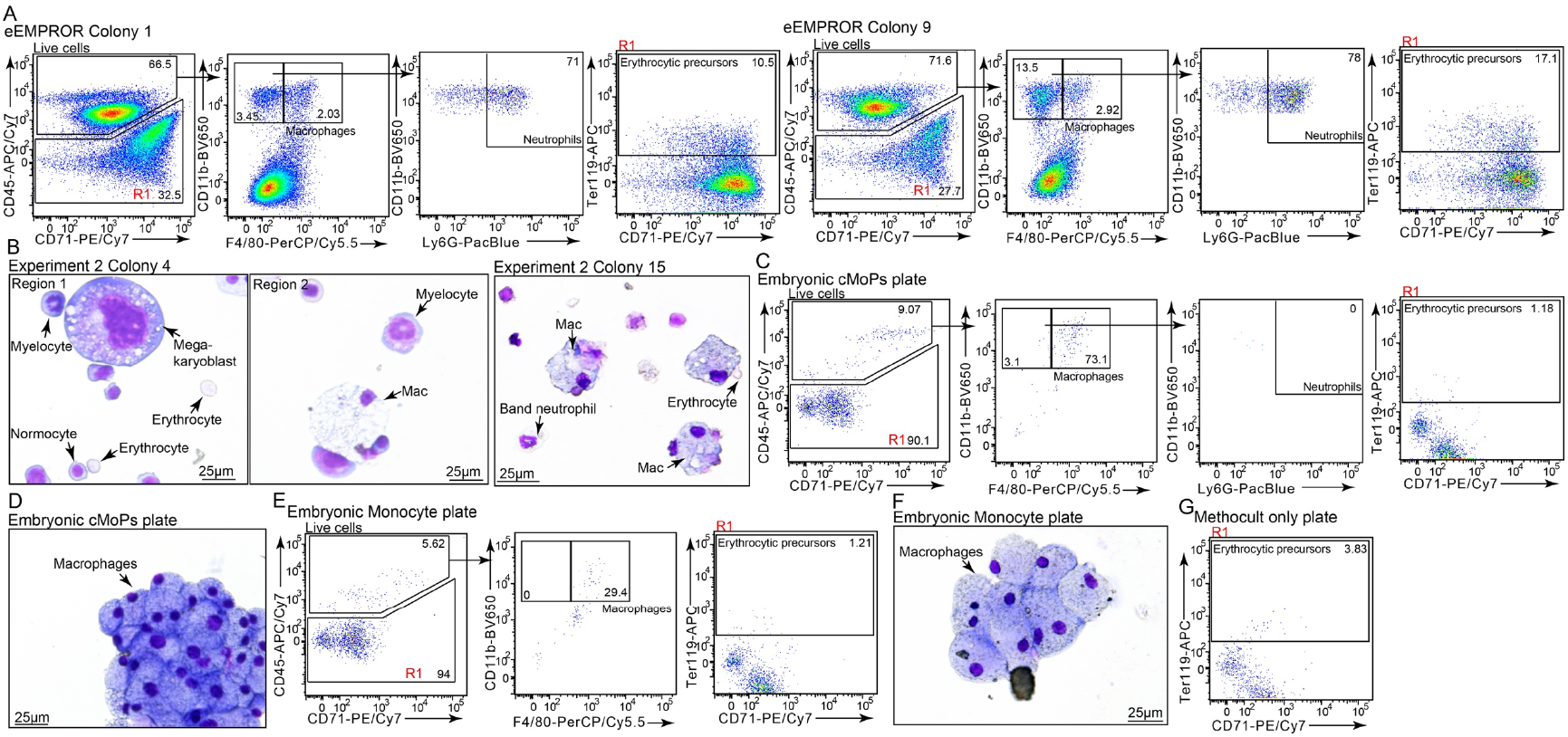
Erythro-myeloid potential of embryonic EMPROR cells *in vitro.* **A**, Flow cytometric analysis of two representative colonies depicting erythroid and myeloid (macrophage and neutrophil) lineage potential of sorted eEMPROR cells in MethoCult™ GF M3434 containing transferrin, SCF, IL-3, IL-6, EPO, TPO, GM-CSF and M-CSF (see methods for details). Data representative of 62 colonies from three independent sort and culture experiments; **B**, Giemsa staining of three independent colonies highlighting erythroid and myeloid lineage potential of sorted eEMPROR cells in MethoCult™ GF M3434 containing transferrin, SCF, IL-3, IL-6, EPO, and supplemented with TPO, GM-CSF and M-CSF. Data represents images from three of 21 independent colonies from one of the three independent sort and culture experiments. **C**, Flow cytometric analysis of cells demonstrating the macrophage differentiation potential of sorted embryonic cMoPs in MethoCult™ GF M3434 containing SCF, IL-3, IL-6, EPO, and supplemented with TPO, GM-CSF and M-CSF. Data representative of colonies from three independent sort and culture experiments; **D**, Giemsa staining of cells obtained from cMoPs culture displaying macrophage lineage potential. Data representative of one of three independent sort and culture experiments; **E**, Flow cytometric analysis of colonies revealing the restricted macrophage potential of embryonic monocytes post-culturing in MethoCult™ GF M3434 containing transferrin, SCF, IL-3, IL-6, EPO, and supplemented with TPO, GM-CSF and M-CSF. Data representative of one of three independent sort and culture experiments; **F**, Giemsa staining of colonies delineates the macrophage lineage potential generated post-culture of sorted embryonic monocytes in MethoCult™ GF M3434 containing transferrin, SCF, IL-3, IL-6, EPO, and supplemented with TPO, GM-CSF and M-CSF. Data representative of one of three independent sort and culture experiments. Both embryonic cMoPs and monocytes lack erythroid and neutrophil lineage potential under these conditions; **G**, Flow cytometric analysis of MethoCult™ GF M3434 without any cells highlighting the background signal present in the medium. Data is from a single experiment.

**Supplementary Figure 13:**
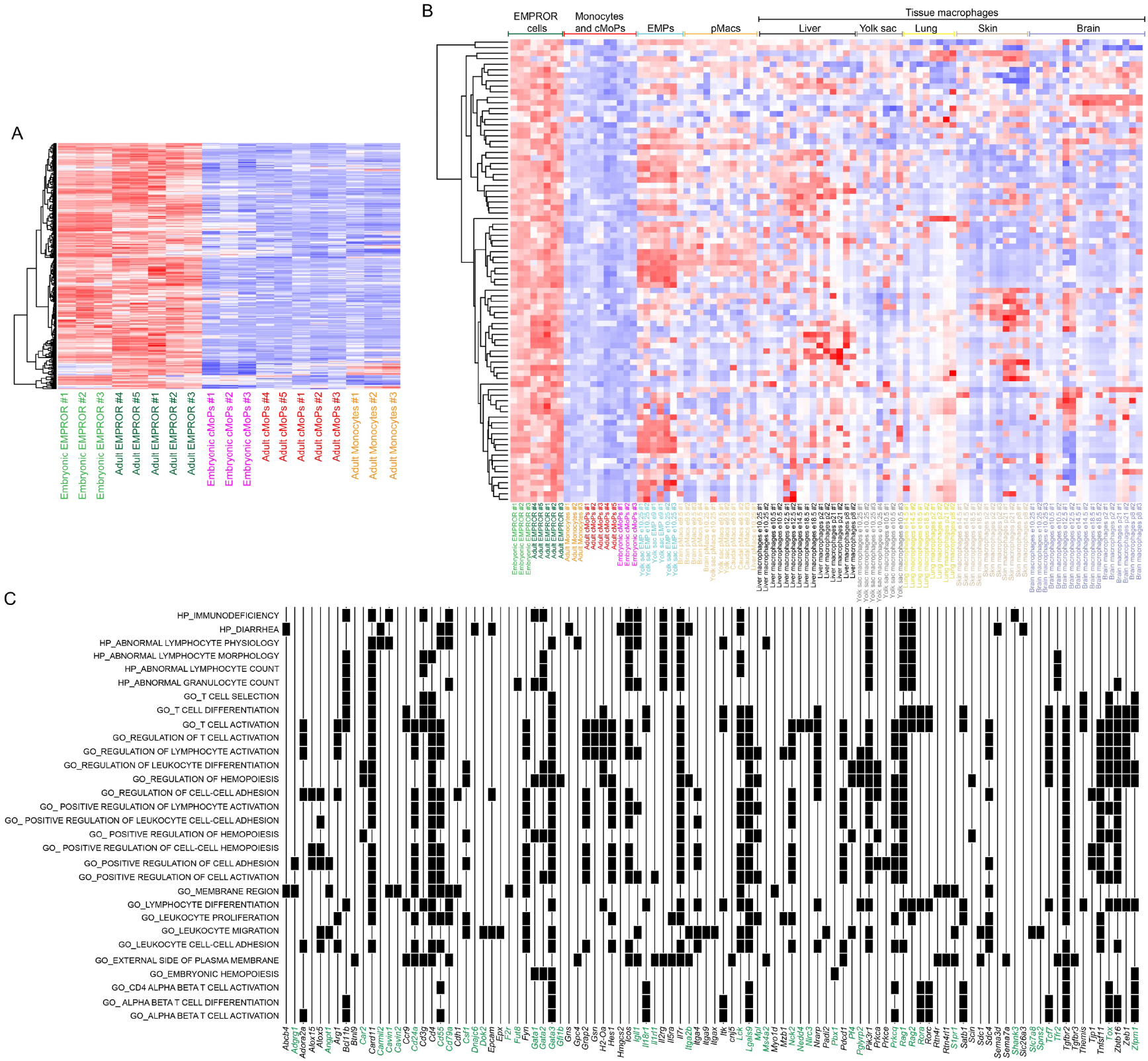
Transcriptional profiling of embryonic and adult EMPROR cells compared to other macrophage precursors and tissue-resident macrophage populations. **A**, Heat map displaying the expression profile of the top 100 genes differentially expressed within EMPROR cells when compared to cMoPs and monocytes. To determine the highly expressed genes within these samples, we calculated intra-class correlation coefficients. The top 100 genes displaying the highest intraclass correlation coefficient and higher median gene expression when compared to cMoPs and monocytes were selected for representation; **B**, Heat map depicting the expression of top 100 genes (as described in panel A) in context to the previously described EMP, pMacs, pre- and post-natal macrophages^3^. Red colour indicates high gene expression, blue indicates low gene expression. Columns are colour coded according to the cell type; **C**, Heatmap highlighting the relationship between top100 genes (as described in panel A) and GO terms. X-axis shows gene names present in at least one gene set. Genes coloured in green were also detected in the previously described Cluster 1 (Supplementary Figure 6C). Y-axis shows gene sets (GO terms). Displayed GO terms have a multiple testing corrected P-value < 0.05. Black squares indicate if a gene is present within gene sets. Note, *Cd4* was present in Cluster IV shared between pDC and EMPROR cells, therefore it is not highlighted as common signature between Cluster I (Supplementary Figure 6C and Supplemental Data 1) and panel C.

**Supplementary Figure 14:**
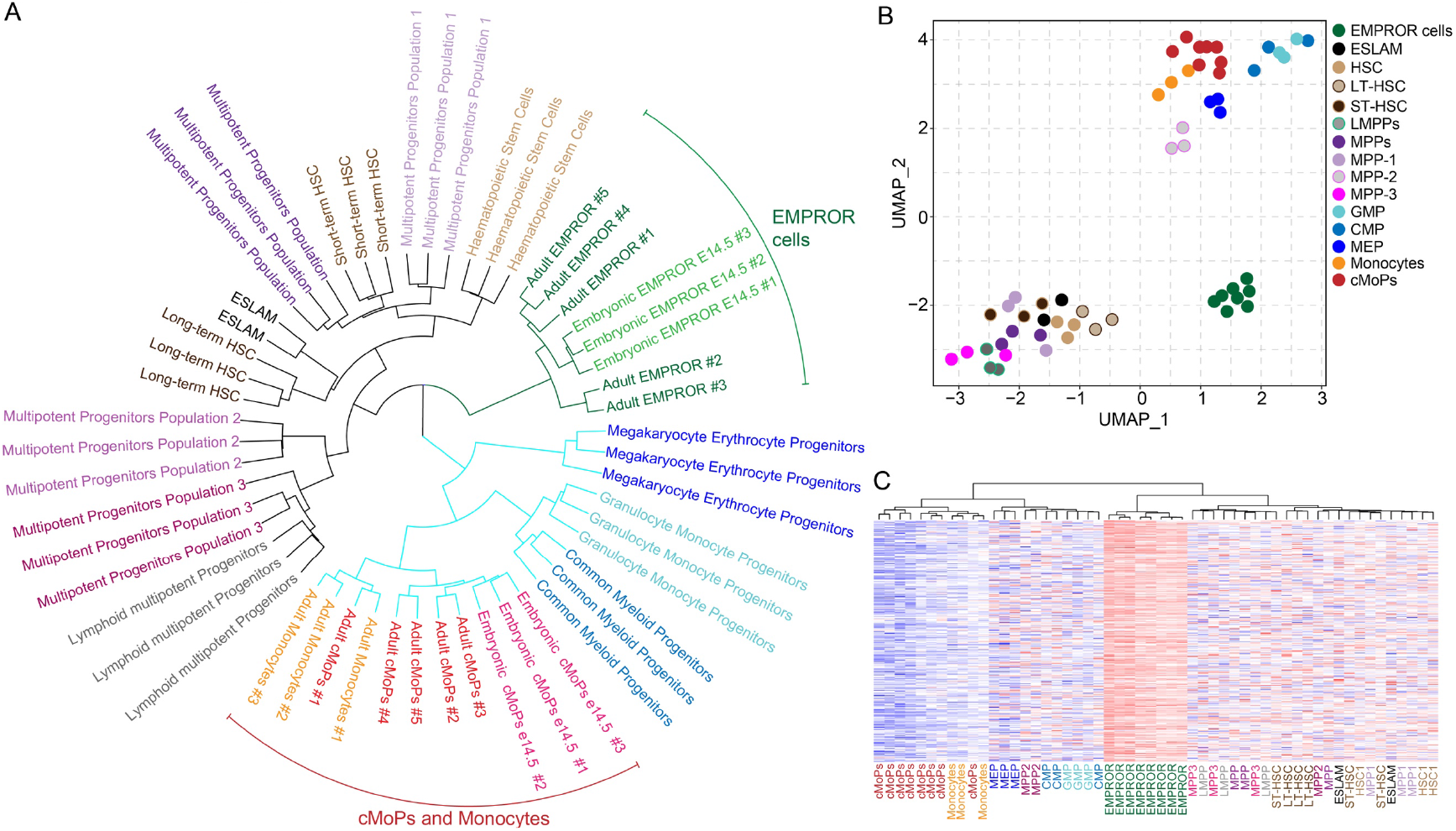
Comparative analysis with scRNA-seq dataset of mouse hematopoietic stem and progenitor cell differentiation. **A**, Dendrogram representation of clustering analysis highlighting the presence of three independent branches consisting of adult and embryonic EMPROR cells, HSCs and multipotent progenitors, and cMoPs, monocytes and CMP, GMP and MEP; **B**, Uniform Manifold Approximation and Projection (UMAP) of the data depicting the presence of three independent cell clusters; **C**, Heatmap analysis of the populations highlighting the distinct gene expression within EMPROR cell population. *Abbreviations:* HSC- Haematopoietic stem cells, LT-HSC- Long-term HSC, ST-HSC- Short-term HSC, LMPP- Lymphoid primed multipotent progenitors, MPP and MPP1- 3- Multipotent progenitors (4 cell populations in total), MEP- Megakaryocyte erythrocyte progenitors, CMP- Common myeloid progenitors, GMP- Granulocyte monocyte progenitors.

## Experimental methods

### Mice

All transgenic mice strains were bred and maintained within the Animal Facility at the Centenary Institute of Cancer Medicine and Cell Biology. Mice were maintained under specific-pathogen-free (SPF) conditions, at 22-26°C with a 12 hours light/dark cycle. C57BL/6 and PTPRC^a^ mice were obtained from commercial vendors, namely Australian BioResources facility (ABR), Moss Vale, New South Wales, Australia and Animal Resource Centre (ARC), Canning Vale, Western Australia, Australia. Mice obtained from other facilities were rested at the Centenary Institute Animal Facility for a week before experiments were performed. *mT/mG* mice^1^ were purchased from The Jackson Laboratory. *DPE-GFP* mice have been described previously^2^. Unless specified, mice between the ages of 6 to 18 weeks of either sex were used for experimentation. All experiments were performed as per the ethics protocols approved by the animal ethics committee, Sydney Local Health District (SLHD) and Animal ethics committee, University of Sydney (USyd). All experiments were performed in accordance with the approved Standard Operating Procedures (SOPs) and flow cytometry biosafety protocols at the Centenary Institute. Adult mice were euthanised by CO_2_ asphyxiation. For embryo collection, pregnant female mice were euthanised and the embryos were surgically excised and exsanguinated by decapitation. Individual strains employed in this study have been listed throughout the manuscript.

#### DPE-GFP Mouse Strain

The DPE-GFP mouse strain was developed by microinjecting a construct comprising of the distal and proximal CD4 enhancer elements based on the pTG construct described by Gerald Siu^3^. In DPE-GFP mice, all T cells (including naïve and effector cells) express GFP^2, 4^. Serendipitously, we subsequently discovered that, in addition to T cells, plasmacytoid dendritic cells (pDC)^5^ and perivascular macrophages (PVM)^6^ show robust GFP expression in DPE-GFP mice.

### Bone marrow chimera generation

6-8-week-old female PTPRC^a^ mice were irradiated (∼10Gy) prior to reconstitution. Bone marrow was harvested from the femurs and tibiae of female donor mice by flushing cells from the medullary cavity employing a 26-gauge needle connected to a 5 ml syringe containing either Hank’s Balanced Salt Solution, 1X HBSS (Gibco, Catalogue no.-14175-095), 1X HBSS +10% FCS or RPMI+10%FCS. The resultant cell suspension was centrifuged at 524g for 5-10 minutes at room temperature. Bone marrow cells were treated with ACK lysing buffer (Gibco, Catalogue no.-A10492-01) for 10 minutes at room temperature to lyse red blood cells. Post-incubation, the ACK lysing buffer was neutralised with sort buffer or RPMI+10%FCS, filtered through 70 µm cell strainer, and centrifuged at 524g for 5-10 minutes at room temperature. The supernatant was discarded and the cell pellet was washed twice with 1X HBSS. Cells were counted and 10 million cells were adoptively transferred into previously irradiated mice. The chimeric mice were supplemented with drinking water containing antibiotics (Sulfamethoxalone 400mg + Trimethoprim 80mg solution at 0.1ml/100ml of drinking water) for 2 weeks to prevent opportunistic infections. In experiments where transgenic RAG-1 deficient mice were utilised for adoptive transfers, 1% of total i.e. 100,000 of either C57Bl/6 or PTPRCA-derived bone marrow cells were co-injected with the 10 million transgenic RAG1 deficient donor cells to repopulate the adaptive immune cell lineages. Chimeric mice were utilised as per the experimental design.

### Flow cytometric analysis of bone marrow precursors

Bone marrow cells were harvested from the femurs and tibiae of 6 – 12-week-old DPE-GFP or DPE-GFP:RAG-1^-/-^ mice. Bone marrow cells were flushed from the bone marrow utilising FACS wash (1XPBS + 2% fetal bovine serum [FBS, HyClone, Catalogue no.- sh30084.03] + 0.2% EDTA + 0.2% Sodium azide), filtered through a 70 µm cell strainer (BD Pharmingen), and centrifuged at 524g for 5-10 minutes at 4°C. Bone marrow cells were treated with ACK lysing buffer (Gibco, Catalogue no.- A10492-01) for 5-10 minutes at room temperature to lyse red blood cells. Post-incubation, the ACK lysing buffer was neutralised with FACS wash, filtered through 70 µm cell strainer, and centrifuged at 524g for 5-10 minutes at 4°C. The supernatant was discarded and the cell pellet was distributed into multiple wells to phenotype precursors cells employing fluorochrome-conjugated antibodies against specific cell surface markers of interest. To identify EMPROR cells, the following antibodies were used: Lineage (CD3e, Ly6G, NK1.1, CD19, Siglec-H) on PerCP/Cy5.5, CD115 on APC, CD117 on PE/Cy7, and CD135 on APC/Cy7. To delineate the expression of various cell surface markers of interest, the above-mentioned panel was utilised with the marker of choice being rotated on Phycoerythrin fluorophore (PE), Brilliant Violet 421 (BV421) or Pacific Blue (PacBlue). In experiments wherein, the surface expression of various proteins was evaluated utilising the Allophycocyanin fluorophore (APC), CD115 staining was performed utilising PE fluorophore. In experiments wherein, surface expression of various proteins was evaluated utilising Phycoerythrin-Cy7 (PE/Cy7), CD117 staining was performed utilising a PE conjugated antibody. FMO and isotype controls were also used as negative controls for marker expression. Please refer to Table 1 detailed below for the specific list of the antibodies and their clones.

**Table 1:** Flow cytometry antibodies.

| **Antigen** | **Clone(s)** | **Fluorophores** |
| --- | --- | --- |
| B220 | RA3-6B2 | APC, PerCP/Cy5.5, BV650 |
| CCR2 | 475301 | PE |
| CD3e | 145-2C11 | PerCP/Cy5.5, APC, BV650, PE/Cy7 |
| CD4 | RM4-5 | APC, PerCP/Cy5.5, BV421 |
| CD8 | 53-6.7 | APC/Cy7 |
| CD11b | M1/70 | APC/Cy7, BV421, BV650 |
| CD11c | HL3 | PE, PE/Cy7 |
| CD19 | 1D3 | PerCP/Cy5.5, APC/Cy7, BV421 |
| CD24 | M1/69 | PE |
| CD25 | PC61 | PE |
| CD31 | MEC 13.3 | APC, Biotin |
| CD34 | MEC14.7 | BV421 |
| CD41 | eBioMWRReg30 | PE/Cy7 |
| CD43 | 1B11 | APC |
| CD44 | IM7 | APC |
| CD45 | 30-F11 | PerCP/Cy5.5, APC/Cy7 |
| CD45.1 | A20 | PerCP/Cy5.5 |
| CD45.2 | 104 | V500, PerCP/Cy5.5, DyLight800 |
| CD64 | X54-5/7.1.1 | AF647, BV421, PE |
| CD69 | H1.2F3 | PE |
| CD71 | RI7217 | PE/Cy7 |
| CD90.2 | 53-2.1 | PacBlue, PE, V500 |
| CD115 | AFS98 | APC, PerCP/Cy5.5 |
| CD117 | 2B8 | PE, PE/Cy7 |
| CD127 | SB/199 | APC |
| CD150 | 9D1 | APC |
| CD206 | C068C2 | PE |
| CD209b | ER-TR9 | PE |
| CD326 | G8.8 | APC, PE |
| CD335 | 29A1.4 | V450 |
| CX3CR1 | 149006 | PE |
| CXCR3 | CXCR3-173 | PE |
| CXCR5 | 2G8 | PE |
| CD135 Biotin | A2F10 | Not applicable |
| CD16/32 | 93 | PE Cy7 |
| Fc Block | 2.4G2 | Not applicable |
| F4/80 | BM8 | PE, PE/Cy7, PerCP/Cy5.5 |
| Gr-1 | RB6-8C5 | PE, APC/Cy7 |
| IgM | II/41 | APC |
| Ly6C | HK1.4 | PE, BV421 |
| Ly6G | 1A8 | PerCP/Cy5.5, PacBlue |
| Mac-3 | M3/84 | PE |
| MHC II | M5/114.15.2 | PE, V500, AF780 |
| NK1.1 | PK136 | PerCP/Cy5.5, APC |
| Sca-1 | D7 | PerCP/Cy5.5, PE |
| Siglec F | E50-2440 | PE |
| Siglec H | 551 | PerCP/Cy5.5, AF647 |
| Streptavidin APCCy7 | Not applicable | APC/Cy7 |
| Streptavidin PECy7 | Not applicable | PE/Cy7 |
| Streptavidin PacBlue | Not applicable | Pacific Blue (PacBlue) |
| Ter119 | TER-119 | PE, APC |

### Sorting of bone marrow precursor populations

Bone marrow cells were harvested from the femurs, tibiae, and spines of 4-7 × 6 – 12-week-old female DPE-GFP, DPE-GFP:RAG-1^-/-^ or DPE-GFP:mT/mG mice. Bones were crushed using a mortar and pestle with sort buffer (1XHBSS [Gibco, Catalogue no.-14175-095] + 2% fetal bovine serum [FBS, HyClone, Catalogue no.-sh30084.03] + 0.2% EDTA) and filtered through a 70 µm cell strainer. Cells were centrifuged at 524g for 5-10 minutes at 4°C, resuspended in 15 ml of ACK lysing buffer to lyse red blood cells, and incubated for 5-10 minutes at 37°C. ACK lysis was neutralised by the addition of sort buffer, filtered again through a 70 µm strainer, and centrifuged at 524g for 5-10 minutes at 4°C. The supernatant was discarded and the cell pellet was resuspended in sort buffer containing Fc block and biotinylated CD135 antibody, stained for 45 minutes to 1 hour on ice. The cell suspension was washed three times with sort buffer and the bone marrow cells were stained with a secondary antibody cocktail containing the following antibodies: lineage (CD3e, Ly6G, NK1.1, CD19, Siglec-H and CD64) on PerCP/Cy5.5, CD115 on APC, CD117 on PE/Cy7, and Streptavidin-APC/Cy7 (to mark CD135 expression). Cells were sorted on the BD FACS Aria-II. Sorted cells were collected in 5 ml polypropylene tubes containing 100-200µl of 100% FBS (double filtered using 0.45-micron filters). Post-sort purities were evaluated for the sorted populations. Sort populations displaying >95% purity were employed for experiments.

### RNA extraction

Bone marrow was harvested from the femurs, tibiae, and spines of 4-7 × 6 – 12-week-old female DPE-GFP mice. Bone marrow was processed for sorting, as described above. Populations of interest were directly sorted into the RLT buffer to minimise any changes in the transcriptional profile. RNA was extracted from the sorted cell populations using the Qiagen RNeasy^®^ Plus Micro Kit (Qiagen, Catalogue no.-74034) in accordance with the manufacturer’s instructions.

### RNA sequencing for adult bone marrow precursors

Post-precursor sort and RNA extraction, quality of the isolated RNA was assessed using the electrophoresis feature of Agilent 2100 Bioanalyser (Perkin Elmer). High-quality RNA from the sorted cells from three independent experiments were used as an input for HiSeq library preparation. Libraries were run on the Illumina HiSeq 2500 platform employing 100bp single read sequencing, with a minimum of 10 million reads per sample achieved. Data handling, sequence alignment and assembly was carried out in collaboration with the AGRF’s bioinformatics facility.

### Hexagonal Plots and Rose Diagrams

The analysis for *Suppl. Fig 6A* was performed as described below. In brief, QC of RNA-seq data was performed using FastQC^7^. FASTQ files were aligned to *Mus musculus* reference genome grc38 using RSEM^8^ using default parameters. Hexagonal plots were generated with Triwise package in R^9, 10^, utilising recommended parameter settings in the package vignette. Gene expression read counts were normalised using the R package edgeR^11^. Differentially expressed genes were obtained applying the same pipeline utilised in the Triwise vignette which is based on the Limma R package^12, 13^. Black dots represent differentially expressed genes, gray dots represent other genes in the dataset. Each of the 120-degree angle is associated to one of the 3 populations and the distance from the centre is proportional to the fold change difference. For each gene module, differentially expressed genes are shown as red dots in the hexagonal plots (differentially expressed associated to the gene set).

### 2D PCA, volcano plots, K-means clustering and GO terms

The analysis for *Fig. 1F and Suppl. Fig 6B-D* was performed as described below. For gene expression analysis of RNA-seq data (HSCs and LMPPs from GSE76140^14^, CMPs from GSE107495^15^ and the manuscript dataset), the adaptors and low-quality bases assessed using FastQC^7^ 0.11.8 were trimmed by Trimmomatic^16^ 0.38 with the default settings. Trimmed reads were then aligned to the ensembl 86 (GRCm38.p4) reference genome using STAR^17^ 2.5.2a. FeatureCounts^18^ 1.5.1 was subsequently employed to convert aligned short reads into read counts for each sample. The data were then analyzed using R^19^ 3.1.2 and DEseq2^20^ 1.28.1. The ComBat^21, 22^ based on the empirical Bayes method, was applied to the values normalised by variance stabilising transformation from DESeq2 for the batch effect adjustment between samples. Differentially expressed genes of various groups were identified using Wald test^23^, with fold-change > 2, P <0.05 after Benjamini-Hochberg^24^ correction and base mean >1. Unsupervised K-means clustering analysis^25^ was performed using the significantly differentially expressed genes. Gene Ontology analysis was performed using the *enrichGO* function in clusterProfiler R package^26^. P values for enrichment were calculated using a hypergeometric test with Benjamini-Hochberg correction. Principal component analysis (PCA)^27^ plot and heatmap were performed in R.

### Integration with publicly available datasets

The analysis for *Fig. 6A-C and Suppl. Fig 13* was performed as described below. The dataset GSE81686 comprises of EMP, pMacs and macrophages taken from different tissues and at different timepoints during embryonic and early development^28^. We downloaded DeSeq2 normalized count matrix from GSE81686 dataset to serve as input for this analysis. For Fig.6A-B we analyzed RNAseq data as described in pervious section. Briefly, data were aligned with STAR, features counted with feature count, counts were normalised with DeSEQ2. We created a combined dataset consisting of 13190 EntrezIDs present in both studies across 95 samples. The combined dataset was scaled and centered. Batch effect introduced by two independent RNAseq analysis in the present study combined with published dataset was removed using ComBat batch removal procedure^29^. Using linear mixed models^30^, we decomposed the variance for every gene in the dataset into variance introduced by batch, experiment (i.e. GSE81686 and the manuscript datasets), age (i.e. embryonic, post-natal, adult) and cell type. For principal component analysis we used the top 1000 genes explaining the most variance accounted to cell type (Supplementary table 2). These genes were used as input for Principal component analysis. We used the first 10 principal components for hierarchical clustering and UMAP visualization of the integrated dataset.

### Genes uniformly expressed in adult and embryonic EMPROR cells

To define a set of genes highly expressed in both embryonic and adult EMPROR cells, we calculated single score intra-class correlation coefficients (ICC) for each gene. For this we grouped the data into EMPROR cells (consisting of adult and embryonic), cMOPs (consisting of adult and embryonic) and monocytes. We used the CRAN package “irr” and defined a two-way random model (model 2) evaluating the consistency within each group. The top 100 genes with the highest intra-class correlation coefficients as well as a higher median gene expression than cMoPs and monocytes were selected for heatmap representation (Supplementary table 2). Over representation analysis (Supplementary Figure 13) was performed in previous section.

### RT-PCR analysis

RNA was isolated from the sorted cell populations as described above. For reverse transcription/cDNA synthesis, 1µl of random hexamer primers (Promega, Catalogue no.-C118A) were added to the RNA. The RNA/primer mix was heated at 70°C for 5 minutes, cooled on ice, and spun down. A reverse transcription cocktail was subsequently prepared in the following manner (per reaction): 5µl of M-MLV 5X reverse transcriptase buffer (Promega, Catalogue no.-M531A), 5µl of 10 mM dNTP mix (Bioline, Catalogue no.-BIO-39044), and 1µl of RNaseOUT^TM^ Recombinant Ribonuclease Inhibitor (Invitrogen, Catalogue no.-10777019). 11µl of the cocktail was added to the RNA/primer mix, then 1µl M-MLV reverse transcriptase enzyme was (New England BioLabs, Catalogue no.-M0368) added to each sample. Samples were gently mixed and incubated at 37°C for 1 hour, followed by a heat inactivation step at 92°C for 5 minutes. Newly synthesised cDNA samples were topped up with 50µl nuclease-free water. RT-PCR was carried out using the Bioline SensiFAST^TM^ Probe Lo-ROX Kit (Bioline, Catalogue no.-84005) and Stratagene Mx3005P (Agilent Technologies). Relative gene expressions were normalised by comparison with the expressions of GAPDH and Actin, and analysed using the 2^−ΔΔCT^ method. The specific genes tested are detailed in Table 2. RT-PCR for each gene was carried out with three biological repeats. The analysed data was plotted utilising Prism software (GraphPad Software Inc., CA) to evaluate subset-specific gene expression. Statistical analysis was performed using one-way ANOVA with Tukey’s multiple comparison test.

**Table 2:**
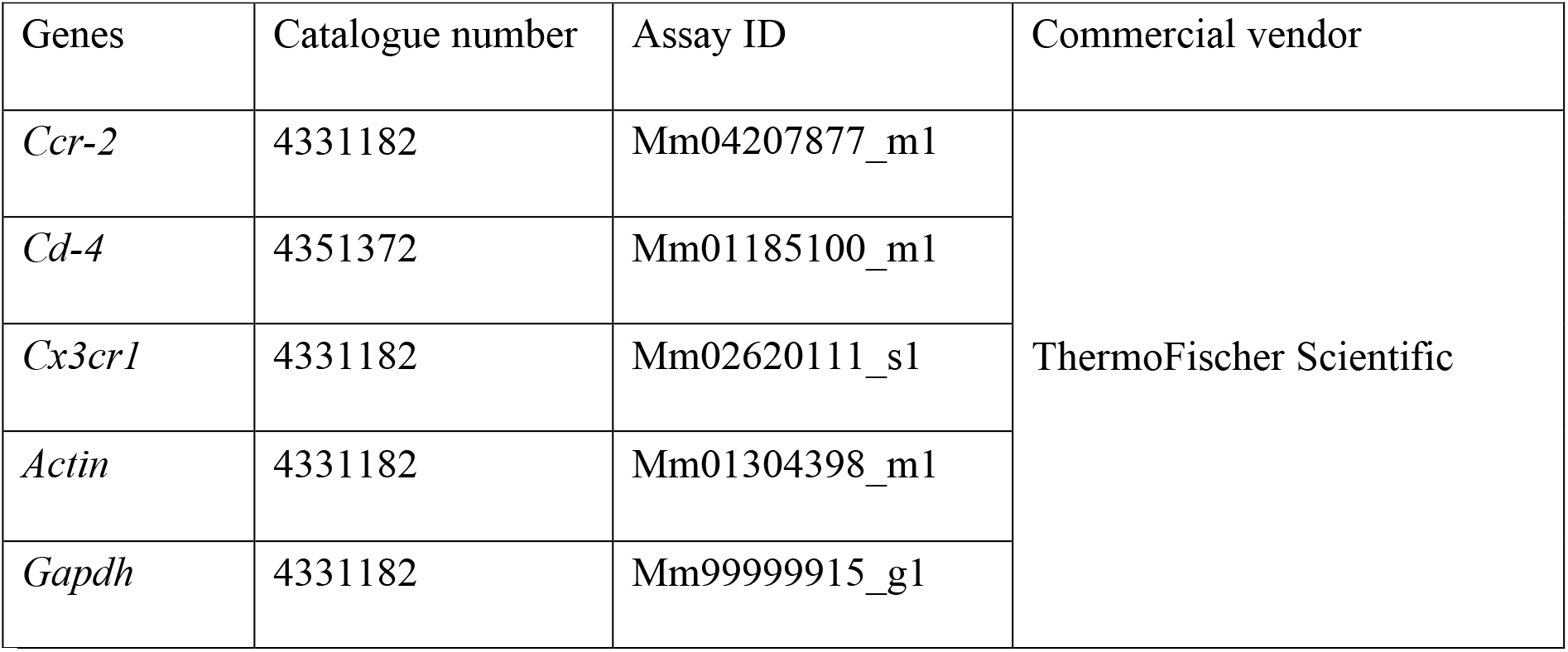
RT-PCR gene sets.

### Meningeal macrophage depletion and adoptive transfer of sorted precursors

Meningeal macrophages were depleted by the administration of mannosylated clodronate liposomes (Clodrosome, Encapsula Nanoscience LLC) into the subarachnoid space. In brief, mice were anaesthetised using Ketamine/Xylazine (100/10 mg/kg of body weight). Post-anaesthesia, 10µl of mannosylated clodronate liposome was injected intrathecally (i.t.) utilising 0.3ml insulin syringe (BD Pharmingen, Bevel length-1mm). The syringe containing the mannosylated clodronate liposome was used to puncture through the parietal bone of the cranium and the liposome solution was injected. A second dose of 10µl of mannosylated clodronate liposome was administered 24 hours later. The treated mice were allowed to rest for 3-4 days, after which sorted immune cell precursors were injected i.t. in 10µl of HBSS as described previously. The mice were allowed to recover for 4-6 weeks and intravital imaging was performed as detailed later.

### Surgical preparation for intravital imaging

a. *Mammary fat pad and tumour imaging*-For imaging mammary fat pad or tumour, a custom-built stage was employed. In brief, mice were anaesthetised utilising Ketamine/Xylazine (100/10 mg/kg of body weight) with repeated half-dose injections as required. Adequate depth of anaesthesia was monitored regularly by checking the hindleg withdrawal and palpebral reflex. The surgical region was dehaired using a commercial hair removal cream, Nair (Church & Dwight). The skin was thoroughly cleaned to remove residual hair removal cream. The mouse was then placed on a temperature controlled surgical board. A lateral incision was made from the lower abdomen to the neck area. Using blunt dissection, the skin was separated from the underlying tissue and laid onto a temperature-controlled imaging platform. The skin flap was held in place using sutures and then superfused with a buffered saline solution (pH 7.4; at 37°C) in a specialised chamber. A cotton-gauze sheet was used to cover the exposed peritoneal wall and the surrounding tissue, which was kept warm with heated (37°C) saline buffer. The exposed mouse mammary fat pad or the developing tumour was then imaged via intravital multiphoton microscopy. Throughout the experiment, the core body temperature was controlled by a rectal thermistor and an electronically regulated siliconised heating pad. The tissue temperature was maintained independently using a regulated heated stage. Non-toxic dyes such as Evans blue conjugated to serum albumin^31^ were injected intravenously to highlight blood vessels.
b. *Brain imaging*-Brain imaging was performed as described previously^32, 33^ with minor modifications. For intravital imaging experiments, adult mice were anaesthetised using Ketamine/Xylazine (100/10 mg/kg of body weight) to achieve surgical plane anaesthesia. Adequate depth of anaesthesia was monitored regularly by checking the hindleg withdrawal and palpebral reflex. Buprenorphine (0.05-0.1 mg/kg of body weight) was administered following induction of anaesthesia to provide multimodal analgesia. Throughout the experiment, the core body temperature was controlled by a rectal thermistor and an electronically regulated siliconised heating pad. Post-anaesthesia, the hair in the submandibular area of the neck and on the skullcap was removed using a commercial hair removal cream, Nair (Church & Dwight). The skin was then cleaned carefully with 1XHBSS to remove residual traces of the cream and hair. The head was immobilised on a using a custom-built stage and care was taken to avoid any respiratory distress. An incision was made in the scalp and the frontoparietal skull was exposed for surgical preparation.

Following immobilisation of the head, a dental drill (Henry Schein Inc.) equipped with a grinding bit (dental bit) was employed to grind a small region of the bone (4-5mm in diameter) to nearly 8-15µm thickness. Saline was added regularly during the bone grinding procedure to avoid generation of heat and to limit any damage. Care was taken to avoid damaging the meninges (Dura mater and Pia mater). The mouse post-cranial window preparation was transferred onto the stage of the intravital microscope and the core body temperature was regularly monitored. Mice were injected intravenously with Evans Blue conjugated to BSA^31^ to visualise the vasculature. In some experiments, 10µl of TRITC-dextran was injected into the subarachnoid space 4-6 hours before cranial window preparation.

### Intravital imaging

Multiphoton microscopy was performed on a LaVision BioTec TriMScope with 20X (NA 0.95) water immersion objective. The set up includes six external non-descanned dual channel reflection/fluorescence detectors, a diode pumped, wideband modelocked Ti-Sapphire femtosecond laser (Mai Tai HP; SpectraPhysics, 720nm-1050nm, <140fs, 90MHz), and an APE optical parametric Oscillator (OPO) system (tuning range 1050-1400nm). Imaging was performed for 270×270µm region at 500×x500 pixel resolution with 6-8 z steps for various time durations as per experimental requirement.

### Confocal imaging

Confocal laser scanning microscopy was performed on the tissue sections or whole mounts as per previously established protocols^6^. Briefly, tissues of interest were harvested and fixed overnight with 4% paraformaldehyde at 4°C. Post-fixation, all tissue samples were washed with immunostaining buffer (1X PBS, 2% FCS, 500mM EDTA, 0.2% Azide, 0.3% Triton X). In case of whole mount staining; for example, mammary fat pad, the intact tissues were directly stained with antibodies, washed and processed for confocal imaging. Other tissues such as tumours, adult liver, gravid uterus and E14.5 embryos were embedded in 5% low-melting agarose and sectioned to 100-200 µm thickness serial sections utilising 1000Plus Sectioning system Vibratome. Tissues were stained with primary antibodies (Table 3) in immunostaining buffer for 48-72 hrs on a rocker shaker at 4°C. Post-staining, the samples were washed thrice with immunostaining buffer at room temperature for minimum 30 min per wash. Tissues were then stained with secondary antibodies (Table 3) in immunostaining buffer overnight on a rocker shaker at 4°C. Post-staining, the samples were washed thrice with immunostaining buffer at room temperature for a minimum of 30min per wash. Tissues were mounted on a cleaned slide using DABCO and imaged using Leica TCS SP5 equipped with an acousto-optical beam splitter and hybrid detectors for sensitive signal detections. Image analysis was carried out using Volocity software (Perkin Elmer, MA).

**Table 3:** Antibodies for immunostaining.

| Antibody | Clone name/Isotype | Dilution | Commercial vendor |
| --- | --- | --- | --- |
| Anti-CD31 | MEC 13.3<br>/Rat IgG 2a | 1:200 to<br>1:400 | BD Pharmingen |
| Anti-Laminin | Not applicable/ Rabbit<br>polyclonal IgG | 1:400 | Sigma Aldrich |
| Anti-F4/80 | Rat hybridoma<sup>34</sup> | 1:3 | Not applicable |
| Anti-Iba1 | Not applicable/ Rabbit<br>polyclonal IgG | 1:200 | Wako |
| Anti-CD163 | Not applicable/ Rabbit<br>polyclonal IgG | 1:200 | SantaCruz<br>Biotechnology |
| Anti-CD169 | ED3/ Rat IgG 2a | 1:400 | AbD Serotech |
| Goat anti-Rat IgG Alexa<br>Fluor 647 | Not applicable /Goat<br>polyclonal Ig G | 1:750 | ThermoFisher<br>Scientific |
| Goat anti-Rat IgG Alexa<br>Fluor 594 |  |  |  |
| Goat anti-Rabbit IgG Alexa<br>Fluor 647 |  |  |  |
| Goat anti-Rabbit IgG Alexa<br>Fluor 405 |  |  |  |

### Lineage potential

**1) B and T cell potential**

**a) OP9/DL-1 System- OP-9 for testing B cell and OP-9/DL1 for testing T cell potential.**

To determine the potential of bone marrow precursors in generating adaptive immune cell lineages, we utilised previously published methodologies with minor modifications^35^. Briefly, two-three days before the initiation of the OP9/OP9-DL-1 co-culture system, OP-9 cells were plated into 10 cm tissue culture dishes (Corning, Catalogue no.-430167) such that the cells were 80 – 90% confluent on the day of co-culture initiation. OP-9 and OP-9-DL-1 cells were cultured in the following media: Minimum Essential Medium Eagle (MEM) (Sigma, Catalogue no.-M8042) + 20% FBS + 2% PenStrep (Gibco, Catalogue no.-15140-122) + 1% L-Glutamine (Gibco, Catalogue no.-25030-081), + 1X tissue-culture grade β-mercaptoethanol (Gibco, Catalogue no.-21985-023). On the day of co-culture, bone marrow was harvested from the femurs, tibiae, and spines of 4 × 6 – 12-week-old female DPE-GFP mice. Bone marrow cells were harvested, processed, and sorted as described above. 25% of the bone marrow cell suspension was separated to sort for CD117^+^Sca-1^+^ lineage^-^(CD3e, CD4, CD11b, CD19, Siglec-H, Ly6G, NK1.1) cells to serve as a positive control for T- and B-cell development. Sorted cells were seeded in 10 cm tissue-culture dishes of 80 – 90% confluent OP9 cells containing 5 ng/ml Flt-3L (R&D, Catalogue no.-427-FL) and 1 ng/ml IL-7 (PeproTech, Catalogue no.- 217-17-10). For T-cell development assay, EMPROR cells, cMoPs, and monocytes were non-trypsin passaged on days 5, 8, and 12 of co-culture, with 25% of the total cell resuspension transferred onto new 10 cm tissue-culture dishes containing 80 – 90% confluent OP9-DL-1 cells with 5 ng/ml Flt-3L and 1 ng/ml IL-7 for each non-trypsin passage. For positive control, only 10% of the total cell resuspension was transferred as the rate of proliferation was much higher in this population. On day 16 of co-culture, all cell suspensions were non-trypsin passaged, washed, stained, and analysed for T-cell development using flow cytometry (BD LSR Fortessa^TM^).

For B-cell development assay, cells were non-trypsin passaged on days 5, 8, 12, and 16 of co-culture, with 25% of the total cell resuspension transferred onto new 10 cm tissue-culture dishes containing 80 – 90% confluent OP9 cells with 5 ng/ml Flt-3L and 1 ng/ml IL-7 for each non-trypsin passage of EMPROR cells, cMoPs, and monocytes. Similar to T-cell development, only 10% of the total cell resuspension was transferred for the positive control to account for the rapid proliferation rate. On day 20 of co-culture, all cell suspensions were non-trypsin passaged, washed, stained, and analysed for B-cell development using flow cytometry.

**b) B cell development-Methocult**

Bone marrow was harvested from the femurs, tibiae, and spines of 3 × 6 – 12-week-old female DPE-GFP mice. For B-cell development, 10,000 sorted cells were plated in 1 ml MethoCult^TM^ M3630 (STEMCELL Technologies, Catalogue no.-03630) containing 2% PenStrep. One week later, colonies were picked and washed twice in FACS wash before being transferred to a 96 well U Bottom plate to be stained for flow cytometric determination of B-cell development. For this experiment, 2x10^5^ total BM cells were plated in the same conditions described above to serve as a positive control for B-cell development.

2) Macrophage potential-Methocult and liquid culture

Bone marrow was harvested, processed, and sorted as described above. For macrophage development, 1000 sorted cells were plated in 1-2ml MethoCult^TM^ GF M3434 (Stemcell Technologies, Catalogue no.-03434) with 50-100 ng/ml M-CSF (PeproTech, Catalogue no.-0315-02-10). One week later, colonies were picked and washed twice in FACS wash (1× PBS, 2% FBS, 2 mM EDTA, 0.01% Azide) before being transferred to a 96 well U Bottom plate to be stained for flow cytometric determination of macrophage development.

Alternatively, sorted cell populations were cultured in macrophage differentiation medium (DMEM Glutamax + Pen Strep + Sodium Pyruvate + 10% FCS) in the presence of 50 ng/ml M-CSF. Post-differentiation, adherent cells were either imaged *in-situ* or harvested for flow cytometric analysis.

**3) Erythro-myeloid potential-Methocult**

Bone marrow was harvested from the femurs, tibiae, and spines of 3 × 6 – 12-week-old female DPE-GFP mice. Bone marrow cells were isolated, processed, and sorted as described above. For assessing the erytho-myeloid potential, duplicate samples containing 800-1000 sorted cells per 1.5 ml MethoCult^TM^ GF M3434 (STEMCELL Technologies, Catalogue no.- 03434) which already contains transferrin, SCF, IL-3, IL-6 and EPO and was further supplemented with 5ng/ml TPO (PROSPEC Biotech, Catalogue no.- CYT-346), 100ng/ml EPO (PROSPEC Biotech, Catalogue no.- CYT- 201), 3ng/ml GM-CSF (PROSPEC Biotech, Catalogue no.- CYT-222) and 5ng/ml M-CSF (PeproTech, Catalogue no.- 0315-02-10). The cell suspension was plated in 2x 35mm petri dishes and incubated at 37°C. Two weeks later, individual colonies were picked up in an unbiased manner. We obtained 20-30 colonies distinct colonies for EMPROR cells and cMoPs. Approximately 10-20 colonies per group/experiment were analysed using flow cytometry, while 5-10 colonies per group/experiment were used for Giemsa staining and cytological evaluation. For monocytes that form 3-4 cell clusters, the complete plate containing Methocult was analysed by flow cytometry, Giemsa staining and cytological evaluation. All samples were washed twice in FACS wash (1× PBS, 2% FBS, 2 mM EDTA, 0.01% Azide) before transferring to a 96 well U Bottom plate to be stained for flow cytometric determination of erythromyeloid potential.

**4) Dendritic cell potential**

Sorted cell populations were cultured as described previously^36^. Briefly, sorted cells were suspended in cell culture medium (RPMI 1640 supplemented with glutamine, penicillin, streptomycin, 2-mercaptoethanol and 10% FCS) in the presence of 20ng/ml GM-CSF (ProSpec) and plated in individual wells of a 6 well plate and placed in cell culture incubator at 37°C maintained at 5% CO_2_. Post-differentiation, cells were harvested for flow cytometric analysis.

**5) Erythro-myeloid potential of embryonic precursors-OP-9 system**

Two to three days before the initiation of the OP9 co-culture system, OP-9 cells were plated into 6 well plates (Corning, Catalogue no.-430167) such that the cells were 80 – 90% confluent on the day of co-culture initiation. OP-9 cells were cultured in the following media: Minimum Essential Medium Eagle (MEM) (Sigma, Catalogue no.- M8042) + 20% FBS + 2% PenStrep (Gibco, Catalogue no.-15140-122) + 1% L-Glutamine (Gibco, Catalogue no.-25030-081), + 1X tissue-culture grade β- mercaptoethanol (Gibco, Catalogue no.-21985-023). On the day of co-culture, MIXER and yolk sac tissues were harvested from the embryos. The tissues were harvested, processed, and sorted as described above. Sorted cells were seeded on top of 80 – 90% confluent OP9 cells containing SCF, IL-3, IL-6, EPO, TPO, GMCSF and MCSF. Plates were routinely monitored and after haematopoietic colonies were visualised, they were flushed with 1XHBSS+10% FCS, washed, stained and analysed for EMP potential using flow cytometry (BD LSR Fortessa^TM^) and cytology.

### Giemsa staining and cell cytology

Giemsa staining was performed as per the previously established protocol^37^. Briefly, colonies from MethoCult cultures were harvested by aspiration and subsequent resuspension in 200µl of FACS buffer in a 96 well U Bottom plate. The plate was centrifuged at 931g for 5 min at room temperature. The cell pellet was washed twice to remove traces of MethoCult M3434. After the final wash, the cell pellet was resuspended in 100µl of 1XHBSS and sorted cells were centrifuged onto glass slides for 5 minutes at 500 RPM (28.23g) (Cytospin3 cytocentrifuge; Shandon). The slides were airdried overnight, fixed with pure methanol for 5 min, air dried and stained with Giemsa stain (Sigma Aldrich, Catalogue no.- GS1L-1L) for 30 min as per the manufacturer’s recommendation. The glass slides were gently washed with tap water to remove excess stain, air dried and imaged. Imaging was performed utilising Leica DM6000B microscope employing a 20X objective. The tiled mosaic for the complete region containing stained cells was acquired and analysed for the presence of various cell lineages.

### Assessment of *in vivo* lineage potential

A day prior to sorting of the precursors, 3 × 6 – 8-week-old female PTPRCA mice were irradiated (∼10Gy). On the day of the sort, bone marrow was harvested from the femurs, tibiae, and spines of 4-6 × 6 – 12-week-old female DPE-GFP:mT/mG mice. Bone marrow cells were harvested, processed, and sorted as described above. Sorted cells were combined with 2-5 million PTPRC^a^-derived bone marrow cells and adoptively transferred into previously irradiated mice. The next day, mice were challenged with tumours by injecting 2.5-3.0 × 10^5^ E0771 murine breast adenocarcinoma cells into the inguinal mammary fat pad of the reconstituted mice. Mice were regularly monitored, and when tumour volumes approached 0.8 cm^3^, mice were euthanised as per the ethics and institutional guidelines and various organs were harvested. Donor-derived immune cells were tracked in various organs utilising the procedures detailed below;

#### Bone marrow

Bone marrow was harvested from the femurs and tibiae. Long-bones were flushed into a 70 µm strainer with FACS wash, and centrifuged at 524g for 5 minutes at 4°C. Bone marrow cells were treated with 5 ml ACK lysing buffer for 10 minutes at room temperature to lyse red blood cells, neutralised with FACS wash, and centrifuged at 524g for 5 minutes at 4°C. The supernatant was discarded and cells were resuspended in FACS wash and transferred to a 96 well U Bottom plate to be stained for flow cytometry. However, for the purposes of analysing the contribution of sorted precursors towards erythrocytic lineage, the ACK lysing step was not performed.

#### Brain

Immune cells from the brain were harvested as per previously established protocols^38^ with modifications. Briefly, whole brains were chopped into small pieces in 1× PBS containing 10% FBS, 1 mg/ml collagenase type IV, and 2% v/v DNase, and then passed through a 19G needle to obtain a homogeneous cell suspension. Brain tissue was incubated in collagenase for 40 minutes at 37°C. Digested brain tissue was filtered through a 100 µm strainer, neutralised with FACS wash, and centrifuged at 524g for 5 minutes at 4°C. The supernatant was discarded, cells were resuspended in 7 ml of 30% Percoll (GE Healthcare, Catalogue no.-17-0891-01), and centrifuged at 931g for 8 minutes at 4°C. The top layer, as well as supernatant, were carefully removed using a transfer pipette. The leftover pellet was resuspended in 2 ml of ACK lysing buffer to lyse red blood cells and incubated for 2 – 3 minutes at room temperature. ACK lysis was neutralised by addition of FACS wash and centrifuged at 524g for 5 minutes at 4°C. Cells were washed an additional two times in FACS wash before being transferred to a 96-well U Bottom plate to be stained for flow cytometry.

#### Liver

Immune cells from the liver were harvested as per previously established protocols^39^. Briefly, liver was perfused *in situ* with 1× PBS containing 10% FBS and 1 mg/ml collagenase type IV before being harvested and cut into small pieces in 1× PBS containing 10% FBS, 1 mg/ml collagenase type IV, and 2% v/v DNase, and then passed through a 19G needle to obtain a homogeneous cell suspension. Liver tissue was incubated for 40 minutes at 37°C. Digested liver tissue was filtered through a 100 µm strainer, neutralised with FACS wash, and centrifuged at 524g for 5 minutes at 4°C. The supernatant was discarded, cells were resuspended in 15 ml of 40% Percoll, and centrifuged at 931g for 8 minutes at 4°C. The top layer, as well as supernatant, were carefully removed using a transfer pipette. The leftover pellet was resuspended in 5 ml ACK lysing buffer to lyse red blood cells and incubated for 5 minutes at room temperature. ACK lysis was stopped by the addition of FACS wash and centrifuged at 524g for 5 minutes at 4°C. Cells were washed an additional two times in FACS wash before being transferred to a 96 well U Bottom plate to be stained for flow cytometry.

#### Skin

Cutaneous tissue was processed as described previously^6^; briefly, ears were split into dorsal and ventral halves before cutting into small pieces in 1× PBS containing 10% FBS, 5 mg/ml collagenase type IV, and 2% v/v DNase, and incubated for 1 hour at 37°C. Digested ear skin tissue was filtered through a 100 µm cell strainer with FACS wash and centrifuged at 524g for 5 minutes at 4°C. RBC lysis was performed in 5 ml ACK lysing buffer for 10 minutes at room temperature, neutralised with FACS wash, and centrifuged at 524g for 5 minutes at 4°C. The supernatant was discarded and cells were resuspended in FACS wash and transferred to a 96 well U Bottom plate to be stained for flow cytometry.

#### Whole Blood

Whole blood was harvested by cardiac puncture (post-CO_2_ asphyxiation) and transferred to a tube containing 500µl Alsever’s solution. Whole blood suspension was topped up with FACS wash and centrifuged at 524g for 5 minutes at 4°C. Supernatant was discarded and cell pellet resuspended in 1 ml FACS wash. 100µl of the cell suspension was transferred to a 96 well U Bottom plate to be stained for flow cytometry.

#### Blood leukocytes

The remaining blood cell suspension was red blood cell-lysed in 10 ml of ACK lysing buffer for 5 minutes at 37°C, neutralised with FACS wash and centrifuged at 524g for 5 minutes at 4°C. The supernatant was discarded and the cell pellet resuspended in 1 ml FACS wash. 200-300µl of the cell suspension was transferred to a 96 well U Bottom plate to be stained for flow cytometry.

#### Thymus and Lung

Whole thymus and lung tissues were chopped with fine scissors, filtered through a 70 µm strainer, topped up with FACS wash, and centrifuged at 524g for 5 minutes at 4°C. Cell suspensions were optionally RBC lysed in ACK lysing buffer for 5 minutes at room temperature depending upon the visual presence of red blood cells, followed by neutralisation with FACS wash and centrifugation at 524g for 5 minutes at 4°C. Supernatants were discarded and cell pellets resuspended in FACS wash and transferred to a 96 well U Bottom plate to be stained for flow cytometry.

#### Tumour

Whole tumours were excised from the mice in 1× PBS. Tumours were washed twice with 1× PBS before being chopped into small pieces and incubated in PBS containing 10% FBS, 1 mg/ml collagenase type IV, and 2% v/v DNase for 1 hour at 37°C. Digested tumour tissue was filtered through a 100 µm cell strainer, topped up with FACS wash, and centrifuged at 524g for 5 minutes at 4°C. Red blood cells were lysed with 5 ml of ACK lysing buffer for 5 minutes at room temperature, followed by neutralisation with FACS wash and centrifugation at 524g for 5 minutes at 4°C. Cells were resuspended in 10 ml FACS wash and filtered through a nylon filter before being overlayed onto 3 ml Ficoll (GE Healthcare, 17-1440-03). Cell/Ficoll mixture was centrifuged at 931g for 1 hour at room temperature with brakes off. Buffy layer was subsequently removed and washed several times with FACS wash before being transferred to a 96 well U Bottom plate to be stained for flow cytometry.

### Timed mating, embryonic tissue harvest and processing for flow cytometric analysis

For timed mating experiments, the following strategy was employed: DPE-GFP^+/+^ female mice (homozygous) were mated with DPE-GFP^+/+^: mT/mG^+/+^ male mice (homozygous). This mating strategy results in DPE-GFP^+/+^:mT/mG^+/-^ embryos. In these mice, the expression of mTomato marks cells of the embryonic origin (**Fig. 3A**). Two to three days prior to the initiation of timed mating, soiled bedding from the cages harbouring DPE-GFP:TOM male mice was transferred to the cages of female DPE-GFP mice (up to 20 weeks of age) to promote synchronisation of estrous cycles (Whitten effect). On the day of timed matings, female DPE-GFP mice were pooled with their respective male DPE-GFP:TOM mice and allowed to mate overnight. After this, every morning, the female mice were checked for vaginal plugs. Female mice once plugged were immediately separated and used as per experimental requirements. Female mice were weighed regularly post-coitus to check for physical signs of weight gain and pregnancy. Pregnant mice were selected at specific time-points, euthanised, and embryos harvested for processing and flow cytometry. Embryos were staged based on the presence of developmental landmarks. E8.5 embryos were staged based on the fusion of headfold, number of somite pairs, turning of the embryo, and allantois development, localisation and absence of fusion with the chorion.

1) At E8.5, yolk sac, embryo proper and MIXER tissues were harvested in 1X PBS containing 10% FBS. The samples were subsequently collagenased using 0.2 mg/ml collagenase type IV (Sigma, C5138-1G) for 20-30 minutes at 37°C with intermittent mixing. They were then very gently passed through a 18G/19G needle to obtain a homogenous sample. The cells were then centrifuged, washed in FACS wash, and transferred to a 96 well U Bottom plate prior to staining for flow cytometry.

For E14.5 embryos, organs were harvested and processed as follows:

***Liver*** – Livers were harvested in 1X PBS, chopped and then passed through a 19G needle to obtain a homogeneous cell suspension. Cells were filtered through a 100µm cell strainer and centrifuged at 754g for 10 minutes at 4°C. Samples were RBC-lysed with ACK lysing buffer for 5-10 minutes at 37°C, neutralised with FACS wash, centrifuged, and resuspended in FACS wash to be transferred to a 96 well U Bottom plate for staining and subsequent flow cytometry-based analysis.

***Blood*** – Blood was harvested from bleeding embryos in Alsever’s solution (0.4% w/v sodium chloride [Sigma, Catalogue no.-310166], 0.8% w/v sodium citrate [Sigma, Catalogue no.-S1804], 0.05% w/v citric acid monohydrate [Chem-Supply, Catalogue no.- CA014], 2% w/v d-Glucose [Sigma, Catalogue no.- G8270] in deionised water, pH 7.3). The blood cell suspension was washed in FACS wash, centrifuged at 754g for 10 minutes at 4°C, red blood cell-lysed in ACK lysing buffer for 10 minutes at 37°C and subsequently neutralised with FACS wash and centrifuged at 524g for 5 minutes at 4°C. The supernatant was discarded and the cell pellet was resuspended in FACS wash and transferred to a 96 well U Bottom plate for staining and subsequent flow cytometry.

***All other organs*** – All other organs were chopped into small pieces in 1× PBS containing 10% FBS, 0.2 mg/ml collagenase type IV, and 2% v/v DNase [Sigma, Catalogue no.- D5025-150KU], and then passed through a 19G needle to obtain a homogeneous cell suspension. Cells were incubated for 30 minutes at 37°C. Cells were then filtered through a 100 µm cell strainer and centrifuged at 754g for 10 minutes at 4°C. Cells were RBC-lysed with ACK lysing buffer for 5-10 minutes at 37°C, neutralised with FACS wash, centrifuged at 754g for 10 minutes at 4°C and resuspended in FACS wash to be transferred to a 96 well U Bottom plate for staining and subsequent flow cytometry.

### Sorting of embryonic precursor populations

Different embryonic tissues such as MIXER, yolk sac, and foetal liver were micro- dissected and processed as below.

1. E8.5 embryo proper, yolk sac and MIXER tissues were harvested in 1X PBS containing 10% FBS. Afterwards, they were collagenased in 0.2 mg/ml collagenase type IV (Sigma, C5138-1G) for 20-30 minutes at 37°C with intermittent mixing. They were then very gently passed through an 18/19G needle to obtain a homogenous sample. Cells were subsequently centrifuged, washed in sort buffer (1XHBSS [Gibco, Catalogue no.-14175-095] + 2% fetal bovine serum [FBS, HyClone, Catalogue no.- sh30084.03] + 0.2% EDTA) and filtered through a 70 µm cell strainer), and transferred to a 96 well U Bottom plate prior to staining for sorting. Cells were sorted on the BD FACS Aria-II. Sorted cells were collected in 5 ml polypropylene tubes containing 100- 200µl of 100% FBS (double filtered using 0.45-micron filters). Post-sort purities were evaluated for the sorted populations. Sort populations displaying >95% purity were employed for experiments.
2. E14.5 foetal livers were harvested in 1X PBS, chopped and then passed through a 19G needle to obtain a homogeneous cell suspension. Cells were then filtered through a 100µm cell strainer and centrifuged at 754g for 10 minutes at 4°C. Cells were RBC-lysed with ACK lysing buffer for 5-10 minutes at 37°C, neutralised with sort buffer (1XHBSS [Gibco, Catalogue no.-14175-095] + 2% foetal bovine serum [FBS, HyClone, Catalogue no.-sh30084.03] + 0.2% EDTA) and filtered through a 70 µm cell strainer), centrifuged, and resuspended in sort buffer to be transferred to a 96 well U Bottom plate for staining and subsequent sorting. Cells were sorted on the BD FACS Aria-II. Sorted cells were collected in 5 ml polypropylene tubes containing 100-200µl of 100% FBS (double filtered using 0.45-micron filters). Post-sort purities were evaluated for the sorted populations. Sort populations displaying >95% purity were employed for experiments.

### RNA extraction

Foetal livers were harvested from E14.5 time-mated mice and processed for sorting, as described above. Populations of interest were directly sorted into the RLT buffer to minimise any changes in the transcriptional profile. RNA was extracted from the sorted cell populations using the Qiagen RNeasy^®^ Plus Micro Kit (Qiagen, Catalogue no.- 74034) in accordance with the manufacturer’s instructions. Note, to minimise any artefacts arising from the analysis of data generated using two different sequencing methodologies, especially when comparing adult precursors cells to their embryonic counterpart, we also sorted adult EMPROR cells and adult cMoPs as described above. These sorted precursors were processed similar to embryonic precursors and were provided for sequencing along with their embryonic counterparts. Two replicates for adult EMPROR cells and cMoPs were processed and have been included in Fig. 6 and Suppl. Fig 13 as replicate sample #4 and #5 for each group.

### RNA sequencing for embryonic precursors

RNA sequencing for embryonic precursors was carried out at the Ramaciotti Centre for Genomics, UNSW, Australia. Post-precursor sort and RNA extraction, quality of the isolated RNA was assessed using the electrophoresis feature of Agilent 2100 Bioanalyser (Perkin Elmer). High-quality, 2ng RNA from three independent experiments were used as an input for SMARTer Total stranded mammalian pico v2 library preparation. Libraries were run using the Illumina NextSeq 500 2×75bp HO flowcell platform employing 100bp paired-end sequencing.

### Erythro-myeloid potential of embryonic precursors

Foetal liver cells were isolated, processed, and sorted as described above. For determining erythro-myeloid potential, duplicate samples containing 500 sorted cells were plated in 1.5 ml MethoCult^TM^ GF M3434 (STEMCELL Technologies, Catalogue no.- 03434) which already contains SCF, IL-3, IL-6 and EPO and was further supplemented with 5ng/ml TPO (PROSPEC Biotech, Catalogue no.- CYT-346), 100ng/ml EPO (PROSPEC Biotech, Catalogue no.- CYT-201), 3ng/ml GM-CSF (PROSPEC Biotech, Catalogue no.- CYT-222) and 5ng/ml M-CSF (PeproTech, Catalogue no.- 0315-02-10). The cell suspension was plated in 2x 35mm petri dishes and incubated at 37°C. Two weeks later, individual colonies were picked in an unbiased manner. We obtained 20-30 colonies distinct colonies for embryonic EMPROR cells and cMoPs. Approximately 20-25 colonies per group/experiment were analysed using flow cytometry, while 5-10 colonies per group/experiment were used for Giemsa staining and cytological evaluation. For monocytes that form 3-4 cell cluster, the complete plate containing Methocult was analysed by flow cytometry, Giemsa staining and cytological evaluation. All samples were washed twice in FACS wash (1× PBS, 2% FBS, 2 mM EDTA, 0.01% Azide) before transferring to a 96 well U Bottom plate to be stained for flow cytometric determination of erythromyeloid potential.

### Statistical analysis

Statistical analyses were performed using Prism software (GraphPad Software Inc). One-way or two-way analysis of variance (ANOVA) followed by either a Bonferroni or a Tukey’s multiple-comparison test (as described in figure legends and individual method sections) was used to determine statistical significance. Statistical difference was assumed if *P* < 0.05.

### Data availability

Data generated in this study is available from the corresponding authors upon reasonable request. All RNA sequencing datasets will be made publicly available as GEO datasets.

### Schematics

Schematics were generated using BioRender (Toronto, Canada)

## Notes

### Competing Interest Statement

The authors have declared no competing interest.

## Manuscript References

1. J. W. Pollard, Trophic macrophages in development and disease. Nat Rev Immunol 9, 259–270 (2009).

2. C. Varol, A. Mildner, S. Jung, Macrophages: development and tissue specialization. Annu Rev Immunol 33, 643–675 (2015).

3. T. A. Wynn, A. Chawla, J. W. Pollard, Macrophage biology in development, homeostasis and disease. Nature 496, 445–455 (2013).

4. T. A. Wynn, K. M. Vannella, Macrophages in Tissue Repair, Regeneration, and Fibrosis. Immunity 44, 450–462 (2016).

5. E. G. Perdiguero, F. Geissmann, The development and maintenance of resident macrophages. Nat Immunol 17, 2–8 (2016).

6. E. Gomez Perdiguero et al., Tissue-resident macrophages originate from yolk-sac-derived erythro-myeloid progenitors. Nature 518, 547–551 (2015).

7. G. Hoeffel et al., C-Myb(+) erythro-myeloid progenitor-derived fetal monocytes give rise to adult tissue-resident macrophages. Immunity 42, 665–678 (2015).

8. K. E. McGrath, J. M. Frame, J. Palis, Early hematopoiesis and macrophage development. Semin Immunol 27, 379–387 (2015).

9. S. Epelman, K. J. Lavine, G. J. Randolph, Origin and functions of tissue macrophages. Immunity 41, 21–35 (2014).

10. K. Kierdorf, M. Prinz, F. Geissmann, E. Gomez Perdiguero, Development and function of tissue resident macrophages in mice. Semin Immunol 27, 369–378 (2015).

11. G. Hoeffel, F. Ginhoux, Ontogeny of Tissue-Resident Macrophages. Front Immunol 6, 486 (2015).

12. F. Ginhoux, M. Guilliams, Tissue-Resident Macrophage Ontogeny and Homeostasis. Immunity 44, 439–449 (2016).

13. M. Guilliams, A. Mildner, S. Yona, Developmental and Functional Heterogeneity of Monocytes. Immunity 49, 595–613 (2018).

14. M. Laviron, A. Boissonnas, Ontogeny of Tumor-Associated Macrophages. Front Immunol 10, 1799 (2019).

15. C. E. Olingy, H. Q. Dinh, C. C. Hedrick, Monocyte heterogeneity and functions in cancer. J Leukoc Biol 106, 309–322 (2019).

16. L. van de Laar et al., Yolk Sac Macrophages, Fetal Liver, and Adult Monocytes Can Colonize an Empty Niche and Develop into Functional Tissue-Resident Macrophages. Immunity 44, 755–768 (2016).

17. C. L. Scott et al., Bone marrow-derived monocytes give rise to self-renewing and fully differentiated Kupffer cells. Nat Commun 7, 10321 (2016).

18. S. Chakarov et al., Two distinct interstitial macrophage populations coexist across tissues in specific subtissular niches. Science 363, (2019).

19. T. Goldmann et al., Origin, fate and dynamics of macrophages at central nervous system interfaces. Nat Immunol 17, 797–805 (2016).

20. M. Prinz, D. Erny, N. Hagemeyer, Ontogeny and homeostasis of CNS myeloid cells. Nat Immunol 18, 385–392 (2017).

21. H. M. Silva et al., Vasculature-associated fat macrophages readily adapt to inflammatory and metabolic challenges. J Exp Med 216, 786–806 (2019).

22. A. Abtin et al., Perivascular macrophages mediate neutrophil recruitment during bacterial skin infection. Nat Immunol 15, 45–53 (2014).

23. W. Weninger, M. Biro, R. Jain, Leukocyte migration in the interstitial space of non-lymphoid organs. Nat Rev Immunol 14, 232–246 (2014).

24. T. R. Mempel et al., Regulatory T cells reversibly suppress cytotoxic T cell function independent of effector differentiation. Immunity 25, 129–141 (2006).

25. R. Jain, S. Tikoo, W. Weninger, Recent advances in microscopic techniques for visualizing leukocytes in vivo. F1000Res 5, (2016).

26. C. E. Lewis, A. S. Harney, J. W. Pollard, The Multifaceted Role of Perivascular Macrophages in Tumors. Cancer Cell 30, 365 (2016).

27. A. E. Casey, W. R. Laster, G. L. Ross, Sustained enhanced growth of carcinoma EO771 in C57 black mice. Proc Soc Exp Biol Med 77, 358–362 (1951).

28. C. N. Johnstone et al., Functional and molecular characterisation of EO771.LMB tumours, a new C57BL/6-mouse-derived model of spontaneously metastatic mammary cancer. Dis Model Mech 8, 237–251 (2015).

29. D. K. Fogg et al., A clonogenic bone marrow progenitor specific for macrophages and dendritic cells. Science 311, 83–87 (2006).

30. C. Auffray et al., CX3CR1+ CD115+ CD135+ common macrophage/DC precursors and the role of CX3CR1 in their response to inflammation. J Exp Med 206, 595–606 (2009).

31. J. Hettinger et al., Origin of monocytes and macrophages in a committed progenitor. Nat Immunol 14, 821–830 (2013).

32. P. Papathanasiou et al., Evaluation of the long-term reconstituting subset of hematopoietic stem cells with CD150. Stem Cells 27, 2498–2508 (2009).

33. G. A. Challen, N. Boles, K. K. Lin, M. A. Goodell, Mouse hematopoietic stem cell identification and analysis. Cytometry A 75, 14–24 (2009).

34. K. Akashi, D. Traver, T. Miyamoto, I. L. Weissman, A clonogenic common myeloid progenitor that gives rise to all myeloid lineages. Nature 404, 193–197 (2000).

35. T. C. Luis et al., Initial seeding of the embryonic thymus by immune-restricted lympho-myeloid progenitors. Nat Immunol 17, 1424–1435 (2016).

36. J. E. Bolden et al., Identification of a Siglec-F+ granulocyte-macrophage progenitor. J Leukoc Biol 104, 123–133 (2018).

37. R. Holmes, J. C. Zuniga-Pflucker, The OP9-DL1 system: generation of T-lymphocytes from embryonic or hematopoietic stem cells in vitro. Cold Spring Harb Protoc 2009, pdb prot5156 (2009).

38. W. C. Chou, D. E. Levy, C. K. Lee, STAT3 positively regulates an early step in B-cell development. Blood 108, 3005–3011 (2006).

39. K. E. McGrath et al., Distinct Sources of Hematopoietic Progenitors Emerge before HSCs and Provide Functional Blood Cells in the Mammalian Embryo. Cell reports 11, 1892–1904 (2015).

40. J. Helft et al., GM-CSF Mouse Bone Marrow Cultures Comprise a Heterogeneous Population of CD11c(+)MHCII(+) Macrophages and Dendritic Cells. Immunity 42, 1197–1211 (2015).

41. J. Y. Bertrand et al., Three pathways to mature macrophages in the early mouse yolk sac. Blood 106, 3004–3011 (2005).

42. J. Palis, Hematopoietic stem cell-independent hematopoiesis: emergence of erythroid, megakaryocyte, and myeloid potential in the mammalian embryo. FEBS Lett 590, 3965–3974 (2016).

43. E. Mass et al., Specification of tissue-resident macrophages during organogenesis. Science 353, (2016).

44. C. Gekas et al., Hematopoietic stem cell development in the placenta. The International journal of developmental biology 54, 1089–1098 (2010).

45. S. Nestorowa et al., A single-cell resolution map of mouse hematopoietic stem and progenitor cell differentiation. Blood 128, e20–31 (2016).

46. D. A. Hume, K. M. Irvine, C. Pridans, The Mononuclear Phagocyte System: The Relationship between Monocytes and Macrophages. Trends Immunol 40, 98–112 (2019).

47. T. Kitamura et al., Monocytes Differentiate to Immune Suppressive Precursors of Metastasis-Associated Macrophages in Mouse Models of Metastatic Breast Cancer. Front Immunol 8, 2004 (2017).

48. R. A. Franklin et al., The cellular and molecular origin of tumor-associated macrophages. Science 344, 921–925 (2014).

49. D. M. Richards, J. Hettinger, M. Feuerer, Monocytes and macrophages in cancer: development and functions. Cancer Microenviron 6, 179–191 (2013).

50. Y. Zhu et al., Tissue-Resident Macrophages in Pancreatic Ductal Adenocarcinoma Originate from Embryonic Hematopoiesis and Promote Tumor Progression. Immunity 47, 323–338.e326 (2017).

51. R. L. Bowman et al., Macrophage Ontogeny Underlies Differences in Tumor-Specific Education in Brain Malignancies. Cell Rep 17, 2445–2459 (2016).

52. P. L. Loyher et al., Macrophages of distinct origins contribute to tumor development in the lung. J Exp Med 215, 2536–2553 (2018).

53. C. Schulz et al., A lineage of myeloid cells independent of Myb and hematopoietic stem cells. Science 336, 86–90 (2012).

54. S. Yona et al., Fate mapping reveals origins and dynamics of monocytes and tissue macrophages under homeostasis. Immunity 38, 79–91 (2013).

55. K. Aghajani, S. Keerthivasan, Y. Yu, F. Gounari, Generation of CD4CreER(T²) transgenic mice to study development of peripheral CD4-T-cells. Genesis 50, 908–913 (2012).

## Figure legend References

1 Johnson, W. E., Li, C. & Rabinovic, A. Adjusting batch effects in microarray expression data using empirical Bayes methods. Biostatistics 8, 118–127, doi:10.1093/biostatistics/kxj037 (2007).

2 Walker, W. L. et al. Empirical Bayes accomodation of batch-effects in microarray data using identical replicate reference samples: application to RNA expression profiling of blood from Duchenne muscular dystrophy patients. BMC Genomics 9, 494, doi:10.1186/1471-2164-9-494 (2008).

3 Mass, E. et al. Specification of tissue-resident macrophages during organogenesis. Science 353, doi:10.1126/science.aaf4238 (2016).

4 Leek, J. T., Johnson, W. E., Parker, H. S., Jaffe, A. E. & Storey, J. D. The sva package for removing batch effects and other unwanted variation in high-throughput experiments. Bioinformatics 28, 882–883, doi:10.1093/bioinformatics/bts034 (2012).

5 Akashi, K., Traver, D., Miyamoto, T. & Weissman, I. L. A clonogenic common myeloid progenitor that gives rise to all myeloid lineages. Nature 404, 193–197, doi:10.1038/35004599 (2000).

## M&M References

1 Muzumdar, M. D., Tasic, B., Miyamichi, K., Li, L. & Luo, L. A global double-fluorescent Cre reporter mouse. Genesis 45, 593–605, doi:10.1002/dvg.20335 (2007).

2 Mempel, T. R. et al. Regulatory T cells reversibly suppress cytotoxic T cell function independent of effector differentiation. Immunity 25, 129–141, doi:10.1016/j.immuni.2006.04.015 (2006).

3 Adlam, M., Duncan, D. D., Ng, D. K. & Siu, G. Positive selection induces CD4 promoter and enhancer function. Int Immunol 9, 877–887, doi:10.1093/intimm/9.6.877 (1997).

4 Sorensen, E. W. et al. CXCL10 stabilizes T cell-brain endothelial cell adhesion leading to the induction of cerebral malaria. JCI Insight 3, doi:10.1172/jci.insight.98911 (2018).

5 Wolf, A. I. et al. Plasmacytoid dendritic cells are dispensable during primary influenza virus infection. J Immunol 182, 871–879, doi:10.4049/jimmunol.182.2.871 (2009).

6 Abtin, A. et al. Perivascular macrophages mediate neutrophil recruitment during bacterial skin infection. Nat Immunol 15, 45–53, doi:10.1038/ni.2769 (2014).

7 FastQC. <https://www.bioinformatics.babraham.ac.uk/projects/fastqc/> (

8 RSEM. <https://bmcbioinformatics.biomedcentral.com/articles/10.1186/1471-2105-12-323> (

9 Triwise. <https://saeyslab.github.io/triwise/> (

10 van de Laar, L. et al. Yolk Sac Macrophages, Fetal Liver, and Adult Monocytes Can Colonize an Empty Niche and Develop into Functional Tissue-Resident Macrophages. Immunity 44, 755–768, doi:10.1016/j.immuni.2016.02.017 (2016).

11 Robinson, M. D., McCarthy, D. J. & Smyth, G. K. edgeR: a Bioconductor package for differential expression analysis of digital gene expression data. Bioinformatics 26, 139–140, doi:10.1093/bioinformatics/btp616 (2010).

12 Limma. <https://bioconductor.org/packages/release/bioc/html/limma.html> (

13 Ritchie, M. E. et al. limma powers differential expression analyses for RNA-sequencing and microarray studies. Nucleic Acids Res 43, e47, doi:10.1093/nar/gkv007 (2015).

14 Luis, T. C. et al. Initial seeding of the embryonic thymus by immune-restricted lympho-myeloid progenitors. Nat Immunol 17, 1424–1435, doi:10.1038/ni.3576 (2016).

15 Bolden, J. E. et al. Identification of a Siglec-F+ granulocyte-macrophage progenitor. J Leukoc Biol 104, 123–133, doi:10.1002/JLB.1MA1217-475R (2018).

16 Bolger, A. M., Lohse, M. & Usadel, B. Trimmomatic: a flexible trimmer for Illumina sequence data. Bioinformatics 30, 2114–2120, doi:10.1093/bioinformatics/btu170 (2014).

17 Dobin, A. et al. STAR: ultrafast universal RNA-seq aligner. Bioinformatics 29, 15–21, doi:10.1093/bioinformatics/bts635 (2013).

18 Liao, Y., Smyth, G. K. & Shi, W. featureCounts: an efficient general purpose program for assigning sequence reads to genomic features. Bioinformatics 30, 923–930, doi:10.1093/bioinformatics/btt656 (2014).

19 RCoreTeam. R: A language and environment for statistical computing. R Foundation for Statistical Computing., <https://www.R-project.org/> (2018).

20 Love, M. I., Huber, W. & Anders, S. Moderated estimation of fold change and dispersion for RNA-seq data with DESeq2. Genome Biol 15, 550, doi:10.1186/s13059-014-0550-8 (2014).

21 Johnson, W. E., Li, C. & Rabinovic, A. Adjusting batch effects in microarray expression data using empirical Bayes methods. Biostatistics 8, 118–127, doi:10.1093/biostatistics/kxj037 (2007).

22 Walker, W. L. et al. Empirical Bayes accomodation of batch-effects in microarray data using identical replicate reference samples: application to RNA expression profiling of blood from Duchenne muscular dystrophy patients. BMC Genomics 9, 494, doi:10.1186/1471-2164-9-494 (2008).

23 Wald, A. Sequential Tests of Statistical Hypotheses. Ann. Math. Statist 16, 117–186, doi:doi:10.1214/aoms/1177731118 (1945).

24 Benjamini, Y. & Hochberg, Y. Controlling the False Discovery Rate: A Practical and Powerful Approach to Multiple Testing. Journal of the Royal Statistical Society: Series B (Methodological) 57, 289–300, doi:10.1111/j.2517-6161.1995.tb02031.x (1995).

25 Kanungo, T. et al. An efficient k-means clustering algorithm: analysis and implementation. IEEE Transactions on Pattern Analysis and Machine Intelligence 24, doi:10.1109/TPAMI.2002.1017616 (2002).

26 Yu, G., Wang, L. G., Han, Y. & He, Q. Y. clusterProfiler: an R package for comparing biological themes among gene clusters. OMICS 16, 284–287, doi:10.1089/omi.2011.0118 (2012).

27 Jolliffe, I. T. Principal component analysis. 2nd edn, (Springer, 2002).

28 Mass, E. et al. Specification of tissue-resident macrophages during organogenesis. Science 353, doi:10.1126/science.aaf4238 (2016).

29 Leek, J. T., Johnson, W. E., Parker, H. S., Jaffe, A. E. & Storey, J. D. The sva package for removing batch effects and other unwanted variation in high-throughput experiments. Bioinformatics 28, 882–883, doi:10.1093/bioinformatics/bts034 (2012).

30 Hoffman, G. E. & Schadt, E. E. variancePartition: interpreting drivers of variation in complex gene expression studies. BMC Bioinformatics 17, 483, doi:10.1186/s12859-016-1323-z (2016).

31 Vennin, C. et al. Transient tissue priming via ROCK inhibition uncouples pancreatic cancer progression, sensitivity to chemotherapy, and metastasis. Sci Transl Med 9, doi:10.1126/scitranslmed.aai8504 (2017).

32 Mazo, I. B. et al. Hematopoietic progenitor cell rolling in bone marrow microvessels: parallel contributions by endothelial selectins and vascular cell adhesion molecule 1. J Exp Med 188, 465–474 (1998).

33 Shaw, T. N. et al. Perivascular Arrest of CD8+ T Cells Is a Signature of Experimental Cerebral Malaria. PLoS Pathog 11, e1005210, doi:10.1371/journal.ppat.1005210 (2015).

34 Austyn, J. M. & Gordon, S. F4/80, a monoclonal antibody directed specifically against the mouse macrophage. Eur J Immunol 11, 805–815, doi:10.1002/eji.1830111013 (1981).

35 Holmes, R. & Zúñiga-Pflücker, J. C. The OP9-DL1 system: generation of T-lymphocytes from embryonic or hematopoietic stem cells in vitro. Cold Spring Harb Protoc 2009, pdb.prot5156, doi:10.1101/pdb.prot5156 (2009).

36 Helft, J. et al. GM-CSF Mouse Bone Marrow Cultures Comprise a Heterogeneous Population of CD11c(+)MHCII(+) Macrophages and Dendritic Cells. Immunity 42, 1197–1211, doi:10.1016/j.immuni.2015.05.018 (2015).

37 Fraser, S. T., Isern, J. & Baron, M. H. Maturation and enucleation of primitive erythroblasts during mouse embryogenesis is accompanied by changes in cell-surface antigen expression. Blood 109, 343–352, doi:10.1182/blood-2006-03-006569 (2007).

38 Pai, S. et al. Real-time imaging reveals the dynamics of leukocyte behaviour during experimental cerebral malaria pathogenesis. PLoS Pathog 10, e1004236, doi:10.1371/journal.ppat.1004236 (2014).

39 Sierro, F. et al. A Liver Capsular Network of Monocyte-Derived Macrophages Restricts Hepatic Dissemination of Intraperitoneal Bacteria by Neutrophil Recruitment. Immunity 47, 374–388.e376, doi:10.1016/j.immuni.2017.07.018 (2017).

